# Control of cell division by an *Acinetobacter baumannii* protein with a nucleotidyl cyclase-like domain

**DOI:** 10.64898/2026.04.03.716176

**Authors:** Andrew Farinha, Mark W Soo, George Minasov, Nicole L Inniss, Ludmilla Shuvalova, Vimisha Dharamdasani, Jessica Finkler, Oliver Stearns, Trisha Shenoy, Christopher Kim, Karla J F Satchell, Edward Geisinger

**Author notes:** Contributed equally (Andrew Farinha, Mark W Soo).

## Abstract

*Acinetobacter baumannii* has diverged substantially from model γ-proteobacteria, complicating our understanding of the antibiotic-resistant pathogen. A major area of divergence is cell division. *A. baumannii* lacks widely conserved division enzymes such as FtsEX and relies on gene products with no similarity to well-characterized proteins. Key among these is AdvA, which we previously identified by Tn-seq as essential for division. AdvA comprises an N-terminal membrane/periplasmic region linked to an unannotated cytoplasmic domain, and most *advA* transposon insertions were lethal except within the linker. How AdvA functions and why insertions showed this positional bias were unclear. Here we combine genetic, localization, two-hybrid, and structural analyses to define roles for AdvA domains in cell division. Removing the N-terminal region or partially truncating the cytoplasmic domain caused lethal division defects. The lethal division block in AdvA-deficient bacteria was associated with failure to recruit early and late divisome proteins to Z-rings but was suppressed by mutations in late septal synthesis genes (*ftsB* and *ftsW*). The cytoplasmic domain bound the early Z-ring protein ZipA, while the N-terminal region mediated dimerization, and both contributed to Z-ring localization. Finally, the crystal structure of the cytoplasmic domain revealed a highly divergent adenylyl/guanylyl cyclase-like fold lacking canonical catalytic and dimerization sites and possessing unusual features, including a positively charged tip and extra C-terminal helix, critical for ZipA binding and division. The essential C-terminal addition helps explain the transposon and truncation effects and facilitated identification of a distant homolog in *Pseudomonas aeruginosa*. These results reveal an atypical cyclase-like protein governing bacterial division.

**Importance:** The multidrug-resistant sepsis pathogen *Acinetobacter baumannii* poses an urgent threat to public health. How *A. baumannii* accomplishes cell division without key conserved factors is not well understood, but this information could lead to improved antimicrobial strategies. We demonstrate that a protein (AdvA) bearing a previously unrecognized structure has a critical role in cell division in the pathogen. The structure resembles nucleotide cyclase enzymes, but it has lost their typical catalytic properties and instead uses noncanonical sites and structural additions to enable assembly of the cell division machine. This atypical domain is found in other pathogens such as *Pseudomonas aeruginosa*, in which it is also connected to cell division. This work opens new avenues to understand division in multidrug-resistant hospital-acquired bacteria.

## Introduction

*Acinetobacter baumannii* is a Gram-negative γ-proteobacterium that causes a wide range of challenging multidrug-resistant (MDR) diseases in healthcare settings, commonly manifesting as bloodstream infections, pneumonia, wound infections, and sepsis [1]. These diseases affect the most critically ill patients and result in increased hospital stays, medical care costs, and risk of mortality [1]. Resistance to formerly “last-line” antibiotics such as carbapenems is now widespread, leaving few effective treatment options [2]. This problem only worsened during the COVID-19 pandemic, which coincided with increased rates of hospital-acquired carbapenem-resistant *A. baumannii* infections [3, 4]. Due to its broad resistance and ability to spread, *A. baumannii* has been designated an urgent threat to public health and a top priority for research and development of new therapies by the Centers for Disease Control and the World Health Organization [5, 6]. Efforts to inhibit the pathogen will be boosted by a better understanding of its essential cell biology, including the fundamental process of bacterial division, which has the potential to provide multiple essential antimicrobial targets [7].

Our understanding of Gram-negative cell division has derived from research in *Escherichia coli* and other model organisms. The canonical pathway follows three main stages: (1) midcell placement of tubulin-like FtsZ filaments to form the Z-ring; (2) ordered assembly of the divisome, the dynamic multiprotein complex that carries out division; and (3) divisome-mediated constriction and septation [8, 9]. Transitions between these stages depend on the ABC transporter-like complex FtsEX that is conserved across a wide range of bacteria [10]. In stage 1, Z-rings are stabilized and membrane-anchored by two proteins, ZipA and the actin homolog FtsA, and recruit FtsEX [11, 12]. Binding of FtsX to FtsA modulates FtsA oligomerization, likely creating the signal to begin stage 2 protein recruitment, including arrival of FtsK, septal regulator complex FtsQLB, FtsW and FtsI peptidoglycan synthases, and the stage 3 activator FtsN [10, 13]. Stage 3 likewise involves FtsEX, which activates peptidoglycan hydrolysis by amidases and their regulatory partners; this processing triggers FtsN-dependent feedback loops that further activate septal wall synthesis by FtsWI, ultimately driving septation and cell separation [10, 14].

While several aspects of the above division paradigm are likely to apply to *A. baumannii*, the pathogen shows notable differences, despite belonging to the same γ-proteobacterial class as *E. coli*. First, in *A. baumannii*, several divisome proteins are either absent or show substantial evolutionary divergence from their *E. coli* counterparts. Most notably, *A. baumannii* has no orthologs of FtsEX [15, 16]. This complex is conditionally, rather than strictly, essential in *E. coli*, which may explain why it can be lost in *Acinetobacter*. The pathogen also lacks homologs of several Z-ring regulators (ZapBCD, SlmA, SulA) as well as amidases AmiA, B, and C [16, 17]. Although an *A. baumannii* ortholog of FtsN has been identified (ACX60_RS17385/GO593_07290), it shows extensive divergence from the *E. coli* prototype (16% identity [16]) and an alternative annotation for the *A. baumannii* gene (*spor*) has been proposed [18]. Moreover, *A. baumannii* versions of ZipA, FtsQ, and FtsL have greatly diverged (26-27% identity with *E. coli* orthologs) [16]. Second, *A. baumannii* encodes a number of atypical proteins that have no close similarity to any known protein in other organisms yet are linked to cell division [15, 19, 20]. These observations indicate that distinct mechanisms have evolved to drive cell division in *A. baumannii* and highlight the need for research to understand and exploit this essential process for controlling the pathogen.

Central among the *A. baumannii*-specific proteins is AdvA. We discovered AdvA through transposon-sequencing (Tn-seq) gene-antibiotic profiling, which revealed a phenotypic signature defined by hypersusceptibility to fluoroquinolone antibiotics tightly connecting AdvA to division pathways [15]. AdvA depletion resulted in dramatic terminal filamentation and blocked growth, demonstrating the essential nature of the protein [15, 21]. Analysis of the insertion sites in viable mutants within the Tn-seq libraries revealed a highly biased distribution in *advA*, with transposons tolerated only at a specific region roughly halfway into the gene (Fig. 1A, purple and green bars). These insertions would create *advA* hypomorphs producing a partial protein ending at residues 171 or 208-244 (of 435 total), either before or after a central predicted coiled coil (CC). The region N-terminal to these sites is predicted to contain two transmembrane (TM) helices flanking a periplasmic domain [15]. The absence of viable insertions within this region is consistent with an essential function for the N-terminal domains of AdvA. The paucity of viable transposon insertions downstream of residue 244 was surprising, however, because all such insertions would produce larger proteins with intact N-terminal domains. This result suggests that partial C-terminal fragments interfere with AdvA function. Nothing was known about the C-terminal domain other than its predicted cytoplasmic localization.

**Fig 1.**
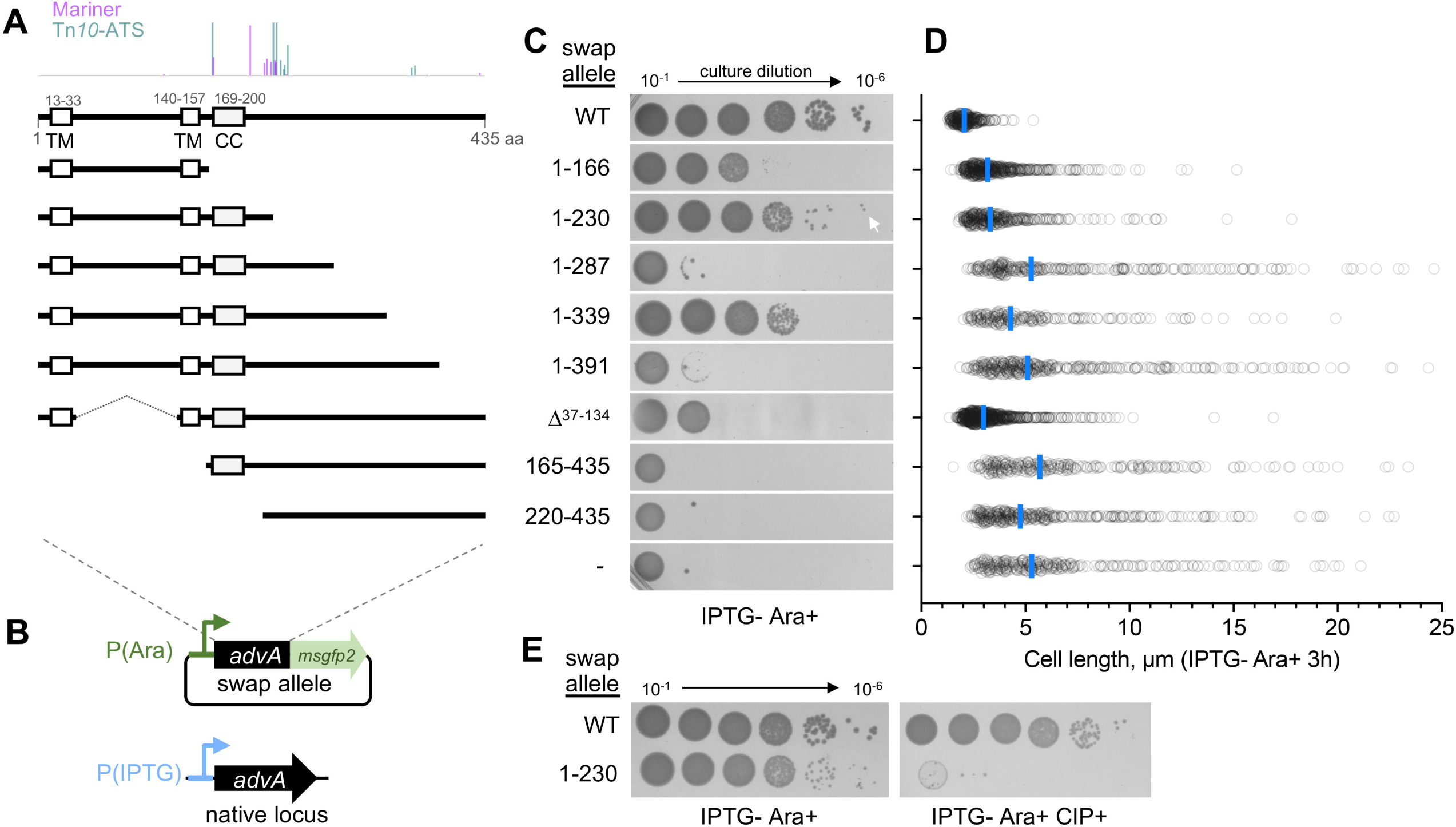
AdvA domain requirements for *A. baumannii* growth and division. (A) Diagrams of the AdvA proteins tested in gene-swap experiments. Bar plot above indicates the corresponding positions of transposon insertions in *advA*. Bar height corresponds to Tn-seq read abundance [21]. Absence of a bar suggests the position is intolerant of transposon insertion. Labels for each construct are given in the adjacent panel; numbers refer to AdvA residues. (B) Diagram of the gene-swap strategy. Swap alleles are fused to msGFP2 and expressed via Ara induction [P(Ara)-*advA-msGFP2*], while endogenous *advA* is under IPTG control at its native chromosomal site [P(IPTG)-*advA*]. (C) Colony growth of *A. baumannii* AdvA swap strains on solid IPTG 0, Ara 0.5% medium. - indicates strain contained an insertless vector (no swap allele). White arrow indicates efficient but smaller colony growth with 1-230 mutant. (D) Cell length in the *A. baumannii* AdvA swap strains from C. Strains were imaged by phase-contrast microscopy after 3 h in IPTG 0, Ara 0.5% medium, and cell length determined via MicrobeJ (n ≥ 232 cells per strain, pooled from 2 independent cultures). Each symbol represents one cell; blue line indicates median. P<0.0001 when comparing each swap allele to WT control by Kruskal-Wallis test. (E) Colony growth of WT and AdvA^1-230^ swap strains on solid IPTG 0 Ara 0.5% medium with or without CIP at 0.0625µg/ml. Images representative of n=3 independent cultures (C, E).

Based on its essentiality and localization to sites of active division [15], we hypothesized that AdvA makes contact with conserved core divisome proteins in ways critical to their function. Consistent with this idea, a recent study detected two-hybrid interactions between AdvA (referred to as Aeg1) and ZipA and FtsN, and identified amino acid changes in FtsA (E202K) and FtsB (E65A) that bypassed AdvA essentiality [22], suggesting that AdvA may function through these core proteins. More work was needed to define the potential binding and controlling functions of AdvA in cell division, clarify the basis for positional effects of transposons in the gene, and understand its structure that has eluded annotation.

In this paper, we address the above problems and reveal the integral role of AdvA and its domains in assembly of the *A. baumannii* divisome. We confirm the N-terminal TM/periplasmic domains of AdvA are essential and mediate dimer formation, and show that the cytoplasmic C-terminal domain influences these N-terminal functions and binds to the early protein ZipA. AdvA is required to assemble the WT divisome, but we identified bypass suppressors in divisome components that function in septum formation, linking AdvA to control of septal synthesis. Finally, we show by X-ray crystallography that the AdvA cytoplasmic domain adopts a three-dimensional fold resembling adenylyl and guanylyl cyclase (AC/GC) enzymes but with major differences not before seen in this enzyme class, including a critical terminal structure that helps explain positional transposon effects in *advA*. Collectively, these findings establish AdvA as an atypical protein central to FtsEX-independent divisome assembly and reveal a cyclase-like fold with an unexpected role in bacterial division.

## Results

### Truncation analysis confirms the positional effects of *advA* insertions and reveals essential protein domains

To examine the basis for the unusual distribution of sites in *advA* tolerating insertional inactivation (Fig. 1A), we engineered a series of *advA* truncation variants and tested them using a dual-promoter gene-swapping strategy. Endogenous *advA* was placed under isopropyl β-D-1-thiogalactopyranoside (IPTG) control at its native locus [P(IPTG)-*advA*], while engineered *advA* alleles fused to GFP were expressed from the arabinose (Ara) promoter via a plasmid [P(Ara)-*advA*-*msGFP2*], such that culturing in IPTG-Ara+ medium swaps endogenous AdvA for the AdvA-GFP construct (Fig. 1B). Control strains behaved as expected, showing efficient growth only when WT AdvA was supplied from either locus (Fig. S1A). We used this system to examine a variety of AdvA mutants: C-terminal truncations ending after TM2 or the CC (aa 1-166 or 1-230, corresponding to positions of most of the viable insertional mutants), as well as three additional downstream locations (1-287, 1-339, or 1-391); cytoplasmic fragments with or without the CC (165-435 and 220-435); and a variant lacking the predicted periplasmic domain by substitution with a short linker (Δ^37-134^) (Fig. 1A). All gene swap strains showed the predicted IPTG-dependent growth pattern on control Ara-media (Fig. S1B). In addition, western blotting confirmed that Ara induction of all the AdvA-GFP fusions yielded intact proteins, with all but one (165-435) at levels reasonably close to the WT control (Fig. S2A).

Analysis of the C-terminally truncated mutants on IPTG-Ara+ swap medium was consistent with the Tn-seq results. First, the 1-230 variant corresponding to the portion of the protein produced in most of the viable transposon mutants enabled nearly 100% colony formation efficiency on the swap medium, albeit with smaller colony size compared to the WT control (Fig. 1C, white arrow). Second, the other C-terminal truncations did not enable efficient colony formation, agreeing with the dearth of viable transposon insertion sites outside of residue positions 208-244 [15] (Fig. 1A,C). Of note, the 1-166 and 1-339 truncations did support partial growth at high seeding densities but not at 10^-5^ and 10^-6^ dilution, consistent with presence of a small number of viable transposon insertions at corresponding positions in *advA* in the Tn-seq data (Fig. 1A,C). To confirm that the ineffective truncations (1-287 and 1-391) were not affected by GFP fusion, we constructed and tested untagged versions. These were just as deficient at supporting growth as the tagged constructs (Fig. S3A). Third, we tested the 1-230 truncation for hypersusceptibility to the fluoroquinolone ciprofloxacin (CIP), a phenotype characteristic of *advA* hypomorphs including the viable transposon mutants examined in Tn-seq screens [15]. The 1-230 truncation was indeed unable to support colony formation during low-level CIP exposure (i.e., at a dose below the MIC for WT *A. baumannii*; Fig. 1E). Together, these results confirm the phenotypes previously identified by Tn-seq and indicate that C-terminal, cytoplasmic segments of AdvA beyond the CC have the potential to negatively modulate the essential function of the N-terminal region of the protein.

To further examine the importance of the N-terminal domains for AdvA function, we tested additional variants lacking the TM helices and/or periplasmic domain. The two N-terminal truncations lacking all of these domains failed to support growth during gene swap, whether or not the CC region was included (165-435 and 220-435; Fig. 1A,C). Similarly, Δ^37-134^ lacking only the periplasmic domain also failed to rescue colony growth (Fig. 1A,C). These results further support the idea that the N-terminal TM and periplasmic domains of AdvA provide functions essential for *A. baumannii* growth.

Lastly, we considered the possibility that the truncations that lacked complementing function might show dominant negative effects. None of the non-complementing variants showed any growth inhibiting activity, however, when overexpressed in WT *A. baumannii* harboring native *advA* (Fig. S3B), suggesting the fragments impair AdvA function only within the same molecule.

### AdvA midcell localization depends on the TM helices and is enhanced by the cytoplasmic domain

To connect the variant growth phenotypes to effects on cell division as well as AdvA localization, we examined gene swap bacteria by microscopy. Western blot analysis revealed that culturing in IPTG-Ara+ medium led to progressive depletion of endogenous WT AdvA, with ∼40% and 10% remaining after 1 and 3 h (T1 and T3), respectively (Fig. S4). We used these time points to examine the cell morphology effects of AdvA variants, focusing on development of increased cell length or filamentation, which is indicative of a block in cell division. At T1, all test strains including the control lacking a swap allele displayed cell length roughly similar to the cells expressing WT AdvA-GFP, indicating that ∼40% residual AdvA levels were sufficient to prevent obvious division block (Fig. S1C). A different situation was seen with the more extensive depletion of native AdvA at T3. Compared to the control strain expressing a WT swap allele, the insertless vector control showed significantly increased median cell length (2 vs >5 µm), as did all AdvA truncations (Fig. 1D). The extent of these cell division defects tended to match the growth defects of each variant in Fig. 1C. For instance, 1-166, 1-230, and 1-339 (growth-complementing) caused relatively modest increases in cell length compared to the 1-287, 1-391 and cytoplasmic fragments (growth-non-complementing), although some cell chaining was also evident in the former set (see Fig. 2B, which shows representative images from Fig. 1D analysis). A notable exception was the Δ^37-134^ variant, which caused a relatively modest cell length defect (∼1.5x increased median length vs WT) despite being ineffective at permitting colony growth on solid medium (Fig. 1D, Fig. 2B). After an additional culture passage in swap medium, the Δ^37-134^ mutant maintained a similar median cell length defect (Fig. S1D).

**Fig 2.**
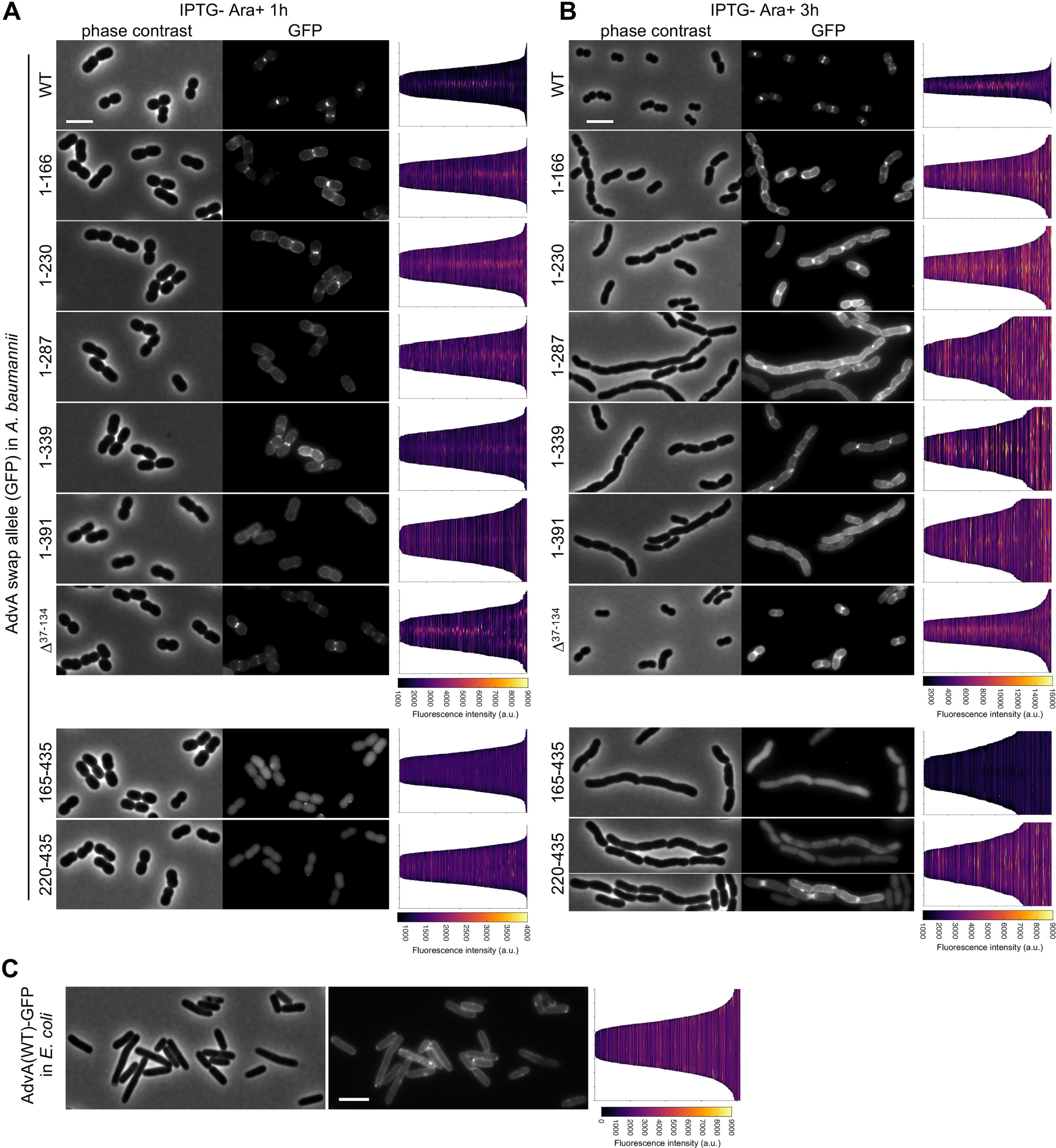
AdvA domain requirements for midcell localization. (A-B) Localization in *A. baumannii*. Swap strains harboring the indicated AdvA-msGFP2 construct were grown 1 (A) and 3 (B) h in swap medium (0.5% Ara, no IPTG) and analyzed by phase-contrast and fluorescence microscopy. (C) Localization in *E. coli.* DH5*a* harboring the same AdvA(WT)-msGFP2 plasmid as in A-B was analyzed after 2 h with 0.1% Ara. Given are representative phase-contrast (left) and fluorescence (center) micrographs and demographs (n≥232 cells per strain, pooled from 2 independent cultures) showing AdvA-msGFP2 fluorescence intensity along medial axes of bacteria (right). In the demographs, the y-axis indicates cell position from -4 to +4 µm. Scale bar, 4 µm.

AdvA localizes to midcell sites of division in *A. baumannii* [15]. We reasoned that a feature of the essentiality of the AdvA N-terminal TM and periplasmic domains was their ability to promote proper localization to these midcell sites. We examined AdvA-GFP localization in the above samples by fluorescence microscopy. WT AdvA-GFP showed the expected localization at midcell and sites of constriction at both T1 and T3 (Fig. 2A, B). In general, the localization of AdvA truncation variants again tended to match their growth effects. 1-166, 1-230, and 1-339 each showed localization to midcell/constriction sites at T1, and this pattern continued at T3 albeit much less efficiently than WT; 1-287 and 1-391 showed mainly peripheral signals with poor midcell localization; and the C-terminal fragments (165-435 and 220-435) yielded nonlocalized cytoplasmic fluorescence, although some diffuse signals at constriction sites were seen sporadically with 220-435 in filamentous cells at T3 (Fig. 2A, B). Interestingly, Δ^37-134^ lacking the periplasmic domain localized to midcell much more efficiently than the other nonfunctional truncations at both T1 and T3, though with increased peripheral signal relative to WT, despite similarly failing to promote colony formation (Fig. 2A, B). Δ^37-134^ differs from the 165-435 construct effectively only by addition of the 2 TM domains. Collectively these results suggest that (1) the TM domains of AdvA are required for localization to division sites, (2) the intact cytoplasmic domain enhances the efficiency of this localization, and (3) the periplasmic domain has an essential function largely separable from midcell localization. Moreover, they show that none of the truncated fragments tested was sufficient to completely phenocopy the growth, cell morphology, and localization conferred by WT AdvA.

Of note, we found that midcell localization of AdvA does not extend to *E. coli*. Full-length AdvA-GFP heterologously expressed in this host localized to the cell periphery, without enrichment at midcell or constriction sites (Fig. 2C). Localization to sites of division thus likely depends on specific molecular determinants of the division apparatus that have evolved in *A. baumannii*.

### AdvA is required for assembly of the divisome

Given the essentiality of AdvA for cell division and its midcell localization, we tested the hypothesis that the protein functions in *A. baumannii* divisome assembly. First, we determined the degree to which divisome protein recruitment to the Z-ring requires AdvA. To this end, we used a dual-label fluorescence strategy enabling concurrent tracking of Z-rings and divisome proteins in cells with or without AdvA. We located Z-rings by using a chromosomal *zapA*-mCherry fusion [23] and tracked a panel of divisome proteins (ZipA, FtsK, -B, -W, and -N) as plasmid-borne, Ara-inducible GFP fusions (Fig. 3A). To analyze the effect of stringent AdvA depletion while cells grew exponentially, we used the conditional P(IPTG)-*advA* allele with IPTG-sucrose (5%) medium, a permissive condition that supports growth despite AdvA depletion (Fig. S5A,C). This permissive effect depends on leaky expression from P(IPTG)-*advA*, since it was blocked by an additional copy of *lacI* or by highly efficient CRISPRi of *advA* [21] (Fig. S5A,B). We passaged cells for 4 h in IPTG-sucrose medium (an initial 3 h passage followed by a second 1 h passage with Ara added to transiently induce the GFP fusions; Fig. S5C) before fluorescence microscopy. In a WT AdvA^+^ control strain background, midcell localization of ZapA-mCherry and all divisome protein-GFP fusions was observed, supporting the utility of this strategy for locating Z-rings and divisome proteins (Fig. 3B).

**Fig 3.**
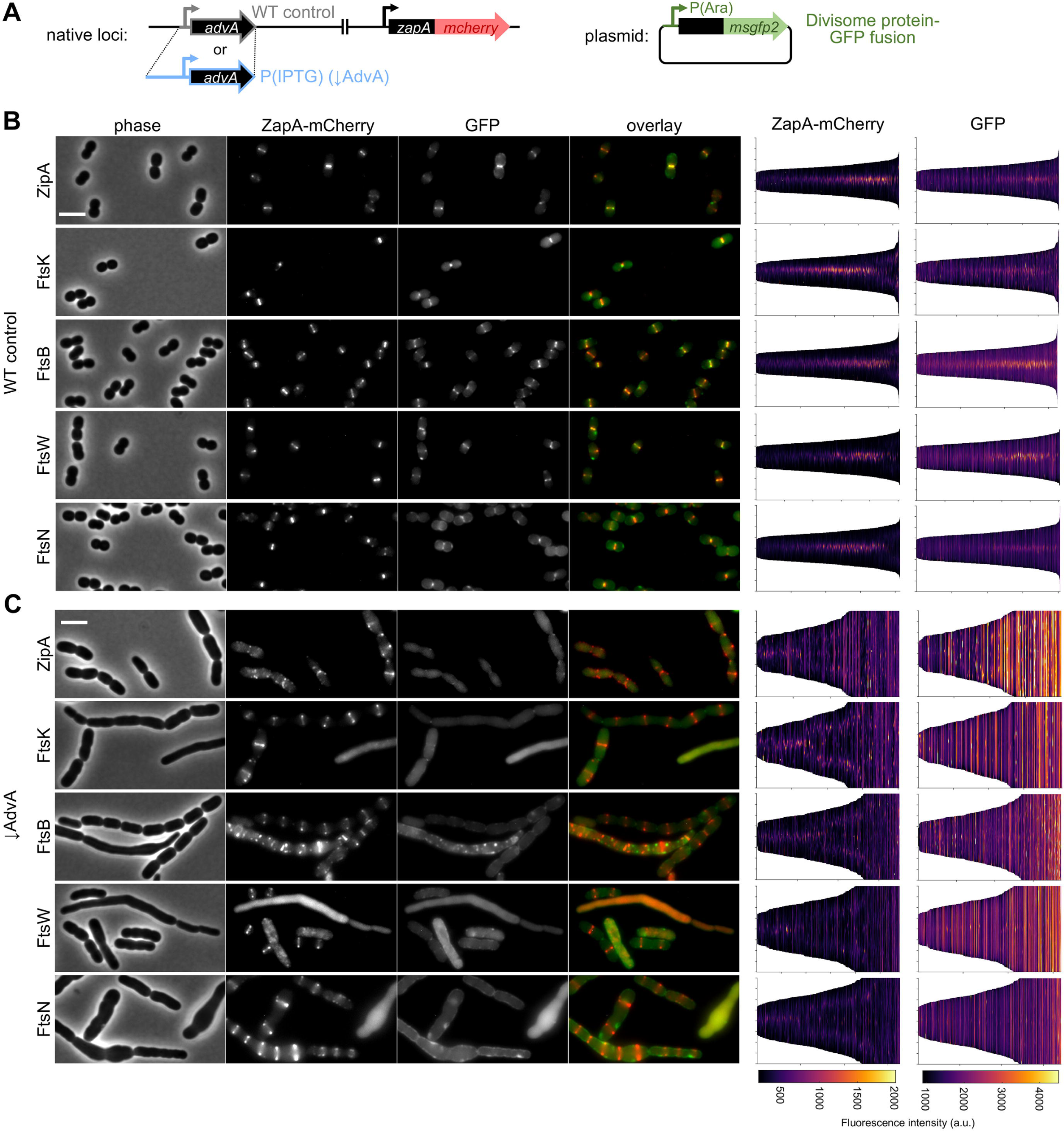
AdvA depletion disrupts recruitment of core divisome proteins to Z-rings. (A) Diagram shows the strategy for tracking localization of division proteins during AdvA depletion in *A. baumannii*. AdvA-depletion test strains contained the chromosomal IPTG-dependent *advA* allele as in Figs 1-2, while control strains contained the native allele. Strains contained a *zapA-mCherry* fusion at the *zapA* locus to label Z-rings and a plasmid-borne, Ara-inducible divisome gene fusion to GFP [P(Ara)-X-*msGFP2*] to track divisome protein localization. (B-C) Representative phase-contrast and fluorescence micrographs (left) and fluorescence profile demographs (right) for WT control (B) and AdvA-depleted test strains (C) as in Fig. 2 (n≥206 cells per strain, pooled from 2 independent cultures). Strains were passaged in IPTG-sucrose 5% medium for 4 h total culture time to allow stringent depletion of AdvA. Division protein-GFP fusions were induced with Ara in the final 1 h of culture before microscopy. Scale bar, 4 µm.

By contrast to the results with WT, AdvA depletion caused profound defects in the ability of both early and late divisome proteins to properly recruit to division sites. In this condition, cell filamentation and chaining occurred, and while a portion of cells showed no localized ZapA-mCherry signal, many cells retained clear ring patterning (Fig. 3C). This signal designating Z-rings can be seen in the demographs in the form of a ZapA-mCherry midline signal in shorter cells and V-patterns in progressively longer/chaining cells. In such cells that had detectable Z-rings, none of the tested divisome proteins efficiently co-localized, indicating defects in the earliest stages of ordered divisome protein recruitment in the absence of AdvA (Fig. 3C). These results support the conclusion that AdvA is a key member of the *A. baumannii* divisome that is required to appropriately assemble the structure and allow cell division.

### The AdvA requirement is bypassed by amino acid changes in FtsB and FtsW

Despite their essentiality under WT conditions, certain *E. coli* core divisome proteins become dispensable when activating mutations alter other components, providing insights into control mechanisms of the divisome. These include mutations in *ftsA* bypassing essentiality of ZipA [24], *ftsL* and *ftsB* bypassing FtsN [25, 26], and *ftsW* bypassing FtsEX [13]. In *A. baumannii*, *ftsA*(E202K) and *ftsB*(E65A) bypass the growth defect of AdvA depletion [22]. We sought additional mutations within division or other loci that could bypass the essentiality of AdvA, potentially informing its roles in divisome function.

To obtain such mutations, we took advantage of the P(IPTG)-*advA-GFP lacI*_2x_ strain and isolated spontaneous IPTG-independent mutants bypassing AdvA. The extra *lacI* served to limit off-target mutations affecting *advA* repression, while the GFP tag enabled secondary screens to exclude them. This yielded two distinct suppressors, both mapping in late divisome genes: *ftsB*(D68G) and *ftsW*(A213V) (Fig. 4A). The affected residues are located in periplasmic (FtsB) and TM (FtsW) sites of the divisome (Fig. S6A). To validate their suppressor functions, we re-engineered each mutation in a different background allowing depletion of AdvA by anhydrotetracycline (aTc)-induced CRISPRi [21]. CRISPRi of *advA* was lethal in an otherwise WT strain but was no longer lethal in the *ftsB*(D68G) or *ftsW*(A213V) mutants, which grew with efficiencies similar to the controls (no induction or CRISPRi with a non-targeting guide RNA, Fig. 4B). The mutations also largely reversed the extreme filamentation characteristic of AdvA depletion, reducing median cell length to levels approaching that of the AdvA-replete controls (Fig. 4C, D). These results validate *ftsB*(D68G) and *ftsW*(A213V) as bona fide *advA* suppressors. The suppressors also showed selective rescue of ZipA, but not FtsK, depletion in analogous CRISPRi colony growth experiments (Fig. S6B). Given the roles of FtsB and FtsW in septal cell wall synthesis and its regulation [13, 25, 27, 28], these findings raise the possibility that AdvA contributes to controlling septal synthesis in addition to recruiting the synthase proteins to midcell, potentially in coordination with ZipA.

**Fig 4.**
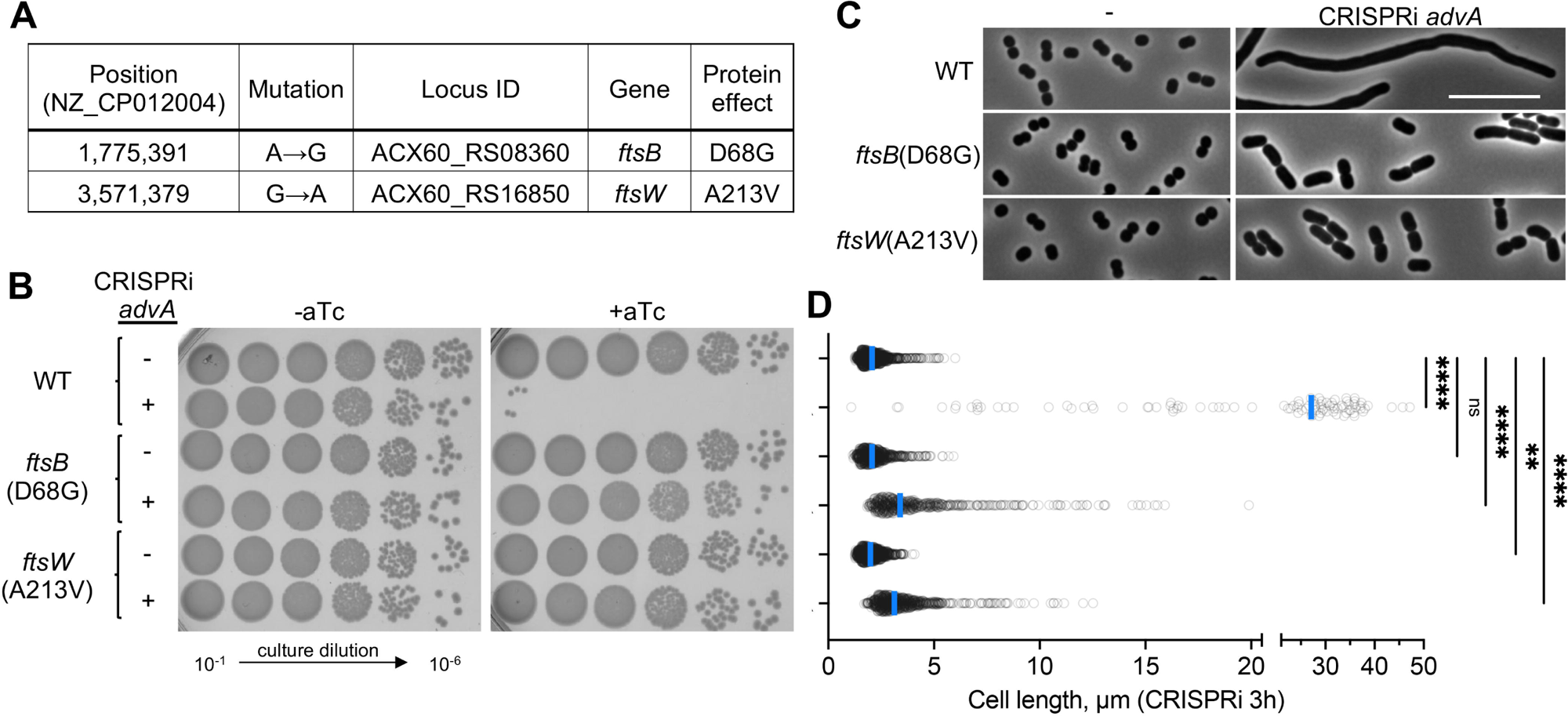
Suppressor mutations in *ftsB* and *ftsW* bypass *advA* essentiality. (A) AdvA^-^suppressors isolated in AFA29 [P(IPTG)*-advA-GFP lacI*_2x_] using non-permissive IPTG-medium. Mutations were identified by breseq analysis. (B) Validation of AdvA^-^suppressors. *ftsB* and *ftsW* mutations were re-constructed in a CRISPRi parental strain. Strains harbored non-targeting control (-) or *advA*-targeting (+) sgRNA. Serial dilutions were spotted on solid medium with or without CRISPRi inducer (aTc, 50ng/ml). Images representative of n=3 independent cultures. (C and D) Suppression of the AdvA^-^division defect. CRISPRi strains from B were cultured for 3 h with aTc and imaged by phase-contrast microscopy. (C) Representative phase-contrast micrographs. Scale bar, 10 µm. (D) Analysis of cell lengths as in Fig. 1D (n ≥ 98 cells per strain, pooled from 2 independent cultures). One data point for WT/CRISPRi-*advA* is not shown (length > 50 µm). ** P<0.01, **** P<0.0001, ns P>0.05 when comparing each strain to WT/CRISPRi-control by Kruskal-Wallis with Dunn’s multiple comparisons test.

### AdvA self-associates and binds ZipA

To further test the hypothesis that AdvA functions in divisome assembly, we examined its self-association and interactions with core divisome components of *A. baumannii*. To this end we used the bacterial adenylate cyclase (AC)-based two-hybrid (BACTH) system [29, 30], in which proteins are fused to one of two fragments from *Bordetella pertussis* CyaA (T18 and T25). Interaction between the proteins reconstitutes AC activity, producing cyclic AMP (cAMP) and a cAMP-receptor protein (CRP)-dependent reporter signal in *E. coli*. We focused on WT AdvA and three truncations: 1-166 (TM and periplasmic domains), 1-230 (TM, periplasmic, and CC domains), and 165-435 (CC and remaining cytoplasmic domains) (Fig. 5A). We fused the C-terminus of each fragment to T18 and T25. Western analysis of the T18 fusion proteins confirmed that each was produced intact at levels similar to WT (Fig. S7A). As a control, we showed that no individual AdvA construct yielded any detectable BACTH signal when expressed alone (i.e., with the insertless complementary vector), indicating that AdvA itself does not raise cAMP levels (Fig. 5A). We tested the ability of AdvA to self-interact by co-expressing both the T18 and T25 fusions. Reporter activity consistent with homodimerization was detected for the WT, 1-166, and 1-230 fragments, while no evidence of self-association was detected for the 165-435 cytoplasmic fragment (Fig. 5A). The N-terminal membrane-tethered portion of AdvA thus appears to be sufficient for dimerization, even in the absence of the cytoplasmic domain or CC.

**Fig 5.**
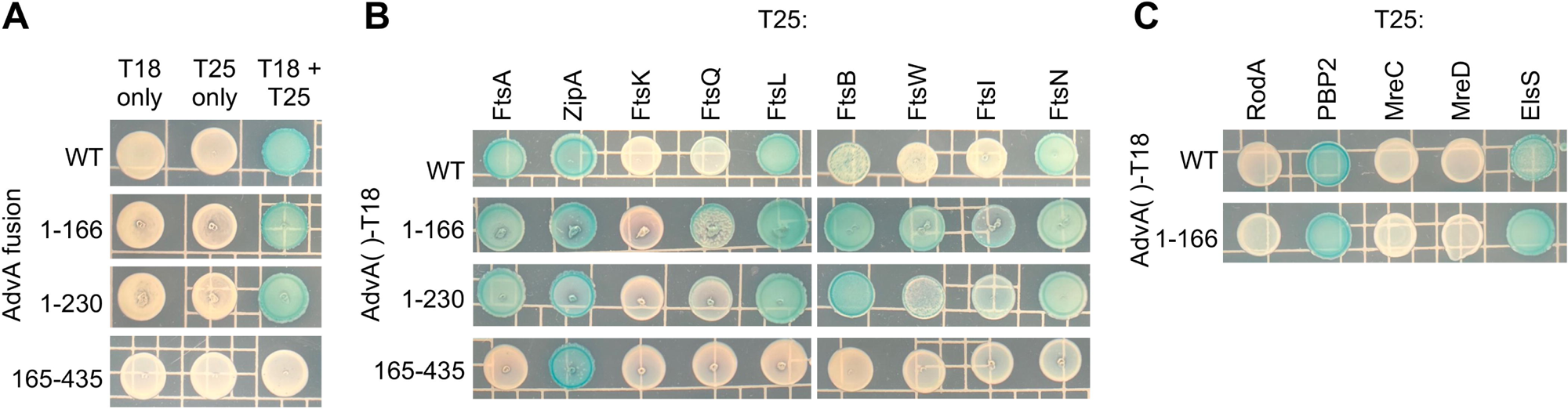
Two-hybrid analysis of AdvA dimerization and divisome protein interactions. Two-hybrid analysis was performed in *E. coli*, with reporter color evaluated after growth on solid indicator medium. (A) Analysis of AdvA homodimerization with the indicated WT or truncated construct fused to T25 or T18 fragment of *B. pertussis* CyaA. In first and second columns, strains contained the insertless vector with the reciprocal CyaA fragment. (B-C) Analysis of AdvA-T18 interactions with T25 fusions to the indicated *A. baumannii* division protein (B) or control membrane protein (C). Images representative of n=3 independent cultures.

We next examined AdvA interactions with core divisome proteins. We constructed a panel of T25 fusions to 9 core *A. baumannii* divisome components (ZipA and FtsA, -K, -Q, -L, - B, -W, -I, and -N); FtsZ was not analyzed because we were unable to construct a CyaA fusion that showed any interaction with its predicted partners, *A. baumannii* FtsA and ZipA [31]. As a control, we also tested a set of membrane proteins not associated with division—previously constructed BACTH fusions to Rod system components RodA, PBP2, MreC, or MreD; and to ElsS, a protein of unknown function linked to the Rod system by phenotypic signatures [15].

When the candidate divisome panel was coexpressed with T18 fusions to AdvA, we detected two-hybrid signals between AdvA^WT^ and FtsA, ZipA, FtsL, and FtsN, as well as a possible weak signal with FtsB (Fig. 5B), several of which overlap with previous findings [22]. The 1-166 and 1-230 AdvA fragments maintained positive two-hybrid signals with the same set of divisome proteins; in addition, the previously weak signal with FtsB appeared to intensify and additional possible weak signals were also observed with FtsW, FtsI, and FtsQ (Fig. 5B). Of note, FtsK showed no BACTH signal with any of the AdvA fusions, but it was functional in that it interacted with complementary fusions to FtsQ, its predicted divisome recruit by analogy with *E. coli* [32] (Fig. S7C). With the control membrane proteins, AdvA and AdvA^1-166^ did not interact with RodA, MreC, or MreD, but they showed a positive two-hybrid signal with PBP2 and ElsS (Fig. 5C). Non-specific signals have been observed previously between membrane proteins in the BACTH assay [33]. Therefore, the signals generated between the transmembrane AdvA constructs (WT, 1-166, 1-230) and membrane-bound divisome proteins (FtsA, ZipA, FtsL, FtsN) should be viewed only as candidate interactions requiring validation.

By contrast with the AdvA transmembrane N-terminal fragments, the cytoplasmic AdvA 165-435 region yielded a two-hybrid signal only with ZipA (Fig. 5B). As the AdvA 165-435 fragment is soluble, its signal with ZipA is unlikely to arise from membrane co-tethering and more likely reflects a specific interaction, which we further test in the structural modeling below. These results are consistent with AdvA binding the early Z-ring protein ZipA.

### The AdvA cytoplasmic domain adopts a noncanonical AC/GC-like structure

To understand how the unannotated cytoplasmic region contributes to AdvA function, we purified recombinant cytoplasmic AdvA (residues 158-435, Fig. 6A) and determined its structure by X-ray crystallography. The crystal structure covered AdvA residues 226-420, possibly from partial hydrolysis during crystallization. The 1.65 Å structure revealed a 3D fold comprised of six α-helices and two short η-helices surrounding a central seven-stranded β-sheet (Fig. 6B).

**Fig 6.**
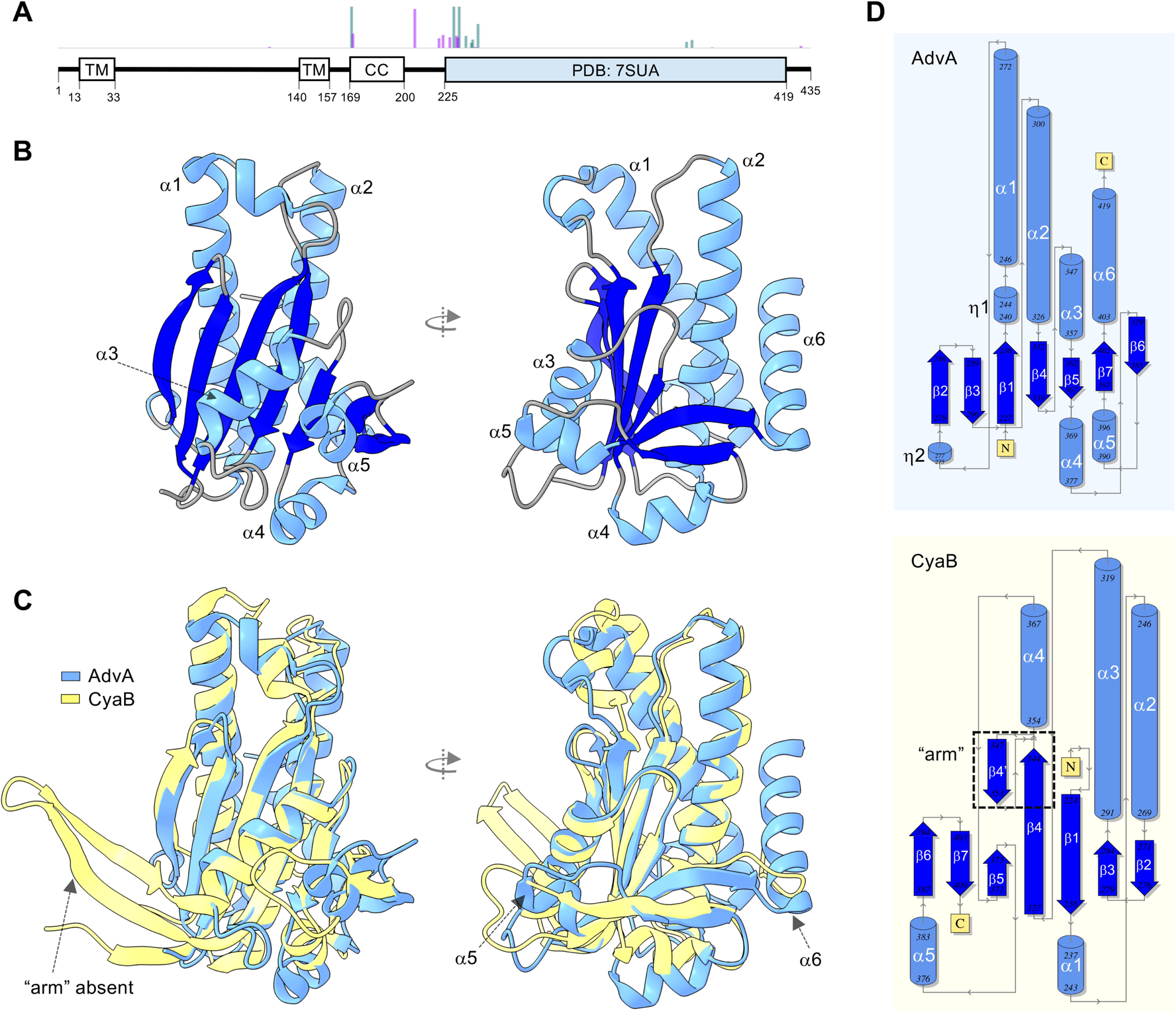
AdvA C-domain crystal structure reveals an atypical AC/GC-like domain with distinct topological features. (A) Linear diagram highlighting AdvA features. Blue shading denotes region for which structure was solved by X-ray crystallography (residues 226-420, PDB: 7SUA). Bar plot above the diagram indicates positions of Tn-seq reads in *advA* (purple, Mariner; green, Tn*10*-ATS) as in Fig. 1A. (B) X-ray crystal structure of AdvA^226-420^ (PDB: 7SUA). Helices are numbered and colored light blue; strands are medium blue; loops are gray. (C) Structural alignment of AdvA^226-420^ with the AC domain of *P. aeruginosa* CyaB (PDB: 3R5G). Key differences between these structures are noted by arrows: AdvA-C lacks the arm domain present in AC/GC enzymes and contains two α-helices not present in most AC/GC enzymes. (D) Protein topology maps of AdvA^226-420^ (top) and CyaB (bottom) generated by PDBsum. Cylinders denote α-helices, large arrows denote β-strands, and small arrows indicate directionality of the protein chain. Boxed is the arm domain of CyaB (bottom) that is absent from AdvA.

Structural similarity searches of the Protein Data Bank (PDB) using the DALI server [34] identified striking resemblance to the cyclase homology domain (CHD) of type III nucleotidyl cyclases (NCs), including the widespread class III ACs and cyclic GMP (cGMP)-producing guanylyl cyclases (GCs) [35]. Almost all of the top structural matches were AC and GC enzymes. For instance, the AdvA^226-420^ structure aligns closely with the AC domain of *Pseudomonas aeruginosa* CyaB [36](Dali Z-score 16.2, RMSD of 2.7 Å over 160 aligned residues, Fig. 6C). Despite the structural similarity, the aligned residues showed very low primary sequence identity (10%). Indeed, while AdvA^226-420^ maintains the characteristic core architecture of ACs such as CyaB, topology analysis revealed that their secondary structure elements have different connectivity (Fig. 6D). These observations are consistent with large evolutionary divergence between AdvA and the canonical AC enzyme.

The AdvA^226-420^ structure has a number of notable features dissimilar from CyaB and other CHD structures. First, AdvA lacks the “arm” formed by 2 antiparallel β-strand extensions in almost all AC and GC enzymes (Fig. 6C). The arm mediates enzyme dimerization, which is critical to formation of the active site [35]. Second, AdvA^226-420^ contains two additional α-helices (α5 and α6) not present in canonical AC/GC enzymes. α5 is situated in a region that is usually a loop in AC/GC enzymes, while α6 (residues 403-419) extends beyond the typical C-terminus of AC/GCs along the surface roughly opposite that of the canonical dimerization arm (Fig. 6C).

This additional terminal helix is absent from all known NC enzymes. Given the lack of the dimerization arm in AdvA, we used AlphaFold to model how the domain may dimerize and assemble relative to well-characterized CHD dimers [37]. Whereas AC/GC enzymes such as CyaB form an arm-mediated, wreath-like dimer with a shared central cleft providing intermolecular catalytic sites (Fig. S8A) [35, 36], all of our structural models of AdvA dimers (of the CHD-like domain alone and of the full-length protein) were low-confidence and placed the CHD-like domains in arrangements lacking this organization (Fig. S8B,C; Table S4). Notably, the full-length model positioned the CHD-like domain adjacent to the CC via an unstructured linker, with no confident interface between two CHD-like domains. Instead, dimer contacts were made along the CC, TM, and periplasmic regions (interface predicted aligned error [PAE] minimum of 3.3 Å despite the low overall model confidence; Table S4, Fig. S8C), consistent with the two-hybrid data for the N-terminal fragments (Fig. 5A). Together with the absence of cytoplasmic fragment self-interaction in the two-hybrid assays (Fig. 5A, AdvA^165-435^), these observations argue against the CHD-like domains forming prototypical, arm-mediated catalytic dimers.

Analysis of the residues of AdvA^226-420^ that share positions with the active sites of AC and GC enzymes revealed further differences between AdvA and canonical cyclases. We used DALI to compare the structures of AdvA^226-420^, CyaB, and another structural homolog, the GC Cya2 from cyanobacterium *Synechocystis PCC6803* [38] (PDB: 2w01; RMSD 2.5 Å over 160 residues aligned with AdvA). CyaB and Cya2 show a number of matching conserved residues, several of which serve as the AC/GC active sites [39] (Fig. S9A, positions indicated by #).

Strikingly, of the 8 AC/GC active site residues, only 2 (R334 and R354) are identical in AdvA, with AdvA either containing distinct residues at the other positions or lacking the position altogether (e.g., T351 on the arm of CyaB) (Fig. S9A,B). In contrast to the residues conserved in the AC/GC protein family, the AdvA family has a distinct spectrum of conserved sites. A region with a prominent number of such sites comprises positions 235-241 and 320-329 (Fig. S9A), which together map to an exposed tip in the AdvA 3D fold involving the C-terminal end of helix α2 and nearby loops (Fig. S9B). Residues at these positions in AdvA are completely dissimilar to those in CyaB and Cya2, and 320-329 in particular corresponds to a region of low conservation in the cyclase proteins.

The absence of most AC/GC active site residues and the critical dimerization arm suggested that AdvA is unlikely to have canonical AC or GC activity. Our findings from multiple *in vivo* and *in vitro* tests agree with this prediction. First, recall that no AdvA construct showed an AC reporter signal when expressed individually in CRP-based BACTH assays (see Fig. 5A). Second, neither full-length AdvA nor two cytoplasmic CHD-containing fragments showed a clear reporter signal in an *E. coli* dual CRP/CRP_G_ reporter system detecting both AC and GC activity [40], in contrast to the clear signals of control enzymes CavA from *A. baumannii* (AC) [41] and GuaA from *Azospirillum* (GC) [40] (Fig. S10A). Third, an *A. baumannii* mutant lacking the CHD-like domain showed no discernible defect in cAMP/cGMP ELISAs (Fig. S10B). Fourth, purified AdvA^158-435^ yielded no detectable pyrophosphate or phosphate release in assays against ATP, GTP, CTP, or UTP (Fig. S10C,D), consistent with absence of NC as well as NTPase activity. Finally, mutating AdvA residues aligning with canonical AC/GC catalytic sites (R334, R354, D283, E350; Fig. S9) did not affect colony formation or CIP susceptibility after gene swap (Fig. S11), consistent with these sites having little importance for AdvA function in division.

Together, these results support the idea that AdvA is not a traditional AC/GC enzyme, showing essentially no canonical NC activity at least under the conditions tested. Instead, the protein contains an atypical cyclase-like structure with numerous features that distinguish it from known enzymes and that may provide distinct mechanisms to control cell division.

### The terminal helix and positively charged tip of the CHD-like domain have critical functions

The extra α6 helix is an unusual addition to the AC/GC fold. Its conservation across the AdvA family suggests the helix may have an important functional role, an idea supported by the previously tested 1-391 variant, which had a truncation site near this helix abolishing its complementing activity. To further test this prediction, we examined an additional truncated *advA* allele deleting the α6 helix (AdvA^1-403^-GFP) (Fig. 7A). The resulting protein was produced intact and at levels equivalent to WT (Fig. S2B), but had dramatic effects on *A. baumannii* division after gene swap. AdvA^1-403^-GFP was completely unable to support colony formation (Fig. 7B) or prevent cell elongation (Fig. 7C,E), in contrast to the parallel WT control allele which supported high-efficiency colony formation and short-rod shape. The AdvA^1-403^ growth defect was observed even when testing an untagged construct (Fig. S3A). In addition, AdvA^1-403^-GFP was defective at midcell localization at 1 and 3 h (Fig. 7D,E) mimicking the results with the 1-287 and 1-391 truncations (see Fig. 2). Lastly, we analyzed AdvA^1-403^ two-hybrid interactions after fusion to CyaA T18 and T25. The AdvA^1-403^-T18 hybrid was produced intact at levels similar to the WT construct (Fig. S7A). AdvA^1-403^ did not form homodimers; however, AdvA^1-403^-T25 yielded a strongly positive BACTH signal when paired with AdvA^WT^-T18, consistent with an ability to form heterodimers (Fig. 7F). Because this signal was weaker in the reciprocal - T18/-T25 pairing, possibly due to differences in vector copy number [30], we tested both orientations for interactions with the *A. baumannii* divisome proteins. AdvA^WT^-T25 reproduced the divisome signals seen with AdvA^WT^-T18 (Fig. 7G). By contrast, AdvA^1-403^ showed no interactions with any divisome protein in either fusion arrangement, apart from a very faint possible signal between AdvA^1-403^-T25 and ZipA-T18 (Fig. 7G, white arrow). The results reveal a critical role for the unusual extended C-terminus of the CHD-like domain in enabling AdvA dimerization, ZipA binding, and the ability of its N-terminal region to generate two-hybrid signals with divisome proteins. More broadly, the *a*6 helix appears essential to AdvA midcell recruitment and division functions.

**Fig 7.**
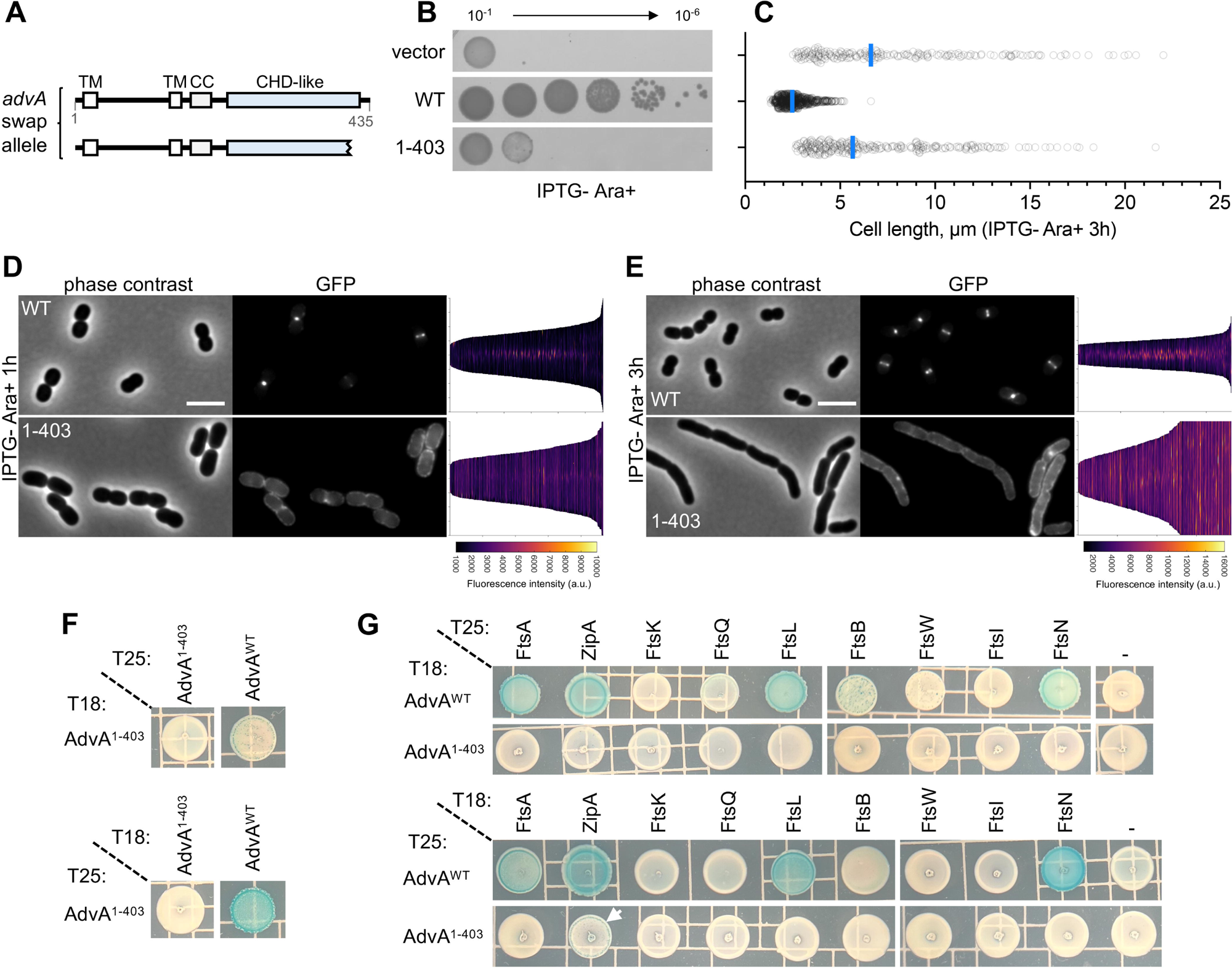
The atypical terminal helix of the cyclase-like domain is critical for AdvA function. (A) Protein map of AdvA^WT^ and AdvA^1-403^ shown as in Fig 1A. (B) AdvA^1-403^ fails to support *A. baumannii* colony growth. Colony growth was examined after gene swap with the indicated AdvA construct fused C-terminally to msGFP2 as in Fig 1B. (C-E) AdvA^1-403^ mislocalizes and causes dramatic cell elongation. The indicated *advA*-*msGFP2* gene swap strains were imaged after 1 and 3 h in IPTG-Ara 0.5% medium by phase-contrast and fluorescence microscopy (n≥206 cells per strain, pooled from 2 independent cultures). (C) Cell lengths from phase contrast images (3 h) analyzed as in Fig 1D, P<0.0001 when comparing vector or 1-403 to WT control by Kruskal-Wallis test. Representative micrographs and demographs of AdvA-msGFP2 subcellular fluorescence are given for 1 h (D) and 3 h (E) time points. (F and G) BACTH analysis in *E. coli* of AdvA^1-403^ dimerization (F) and interactions with *A. baumannii* core divisome proteins (G). - indicates two insertless T18 and T25 vectors. White arrow indicates partial reporter signal with the AdvA^1-403^-T25, ZipA-T18 pairing. Images representative of n=3 independent cultures (B, F-G).

We also examined the importance of the conserved tip of the CHD-like domain. This region includes multiple basic residues, including K239 and R324, which contribute to a prominent positively charged surface surrounding a short cleft (Fig. 8A). We mutated K239 and R324 to alanine and examined mutant complementation and localization phenotypes after gene swap. While both mutant alleles supported *A. baumannii* colony growth on IPTG-Ara+ medium lacking antibiotics, the R324A mutant showed hypersusceptibility to CIP, mimicking the 1-230 truncation which entirely removes the AC/GC-like domain (Fig. 8B). Consistent with its colony growth effects, in antibiotic-free gene-swap medium R324A caused a subtle but significant increase in cell length (Fig. 8C, D). R324A also slightly reduced the efficiency of AdvA midline localization, showing increased peripheral signal compared to the WT control (Fig. 8D). We hypothesized that the susceptibility and localization phenotypes of the R324A mutation in the CHD-like domain are connected to altered ability to interact with the cytoplasmic divisome. To test this, we examined how R324A affects interactions with ZipA, the sole protein found to interact with the AdvA soluble cytoplasmic region (165-435, Fig. 5B). While full-length AdvA^R324A^ retained a BACTH signal with ZipA, the R324A mutation in the cytoplasmic fragment (165-435) largely eliminated this interaction (Fig. 8E). We confirmed this result with fusions to CyaA T18 and T25 fragments in both orientations (Fig. 8E) and showed that the reduced signal is not due to AdvA instability, as protein levels for the T18 fusions to both R324A variants were similar to their respective WT controls (Fig. S7B). These results identify a single key residue in the positively charged tip of AdvA that is important for ZipA interaction, efficient bacterial division, and fluoroquinolone resistance.

**Fig 8.**
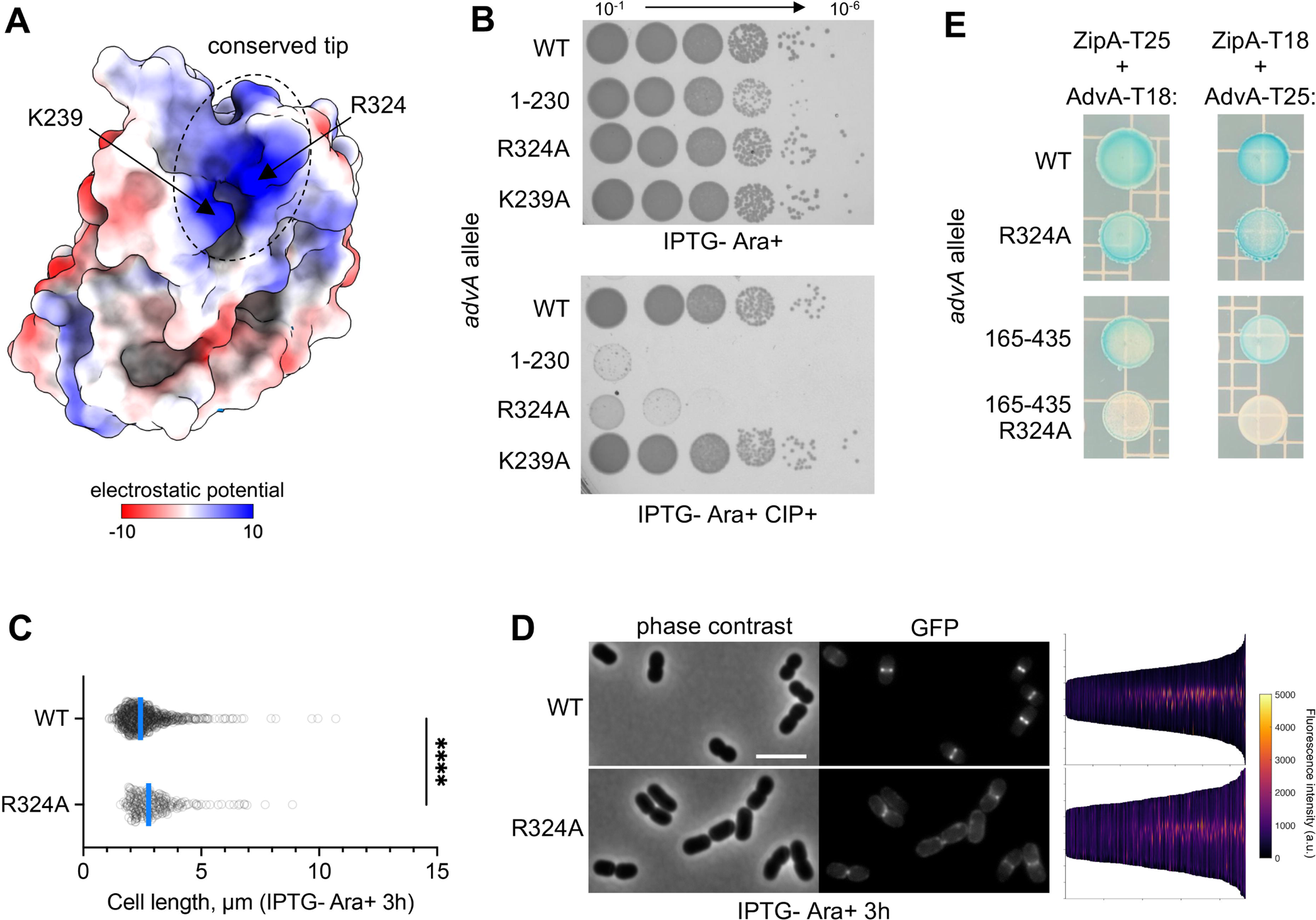
AdvA residue R324 is important for ZipA binding, efficient cell division, and fluoroquinolone resistance. (A) Electrostatic surface potential of AdvA^226-420^ showing the positively charged tip region (head-on view) and positions of residues K239 and R324. (B) AdvA K239 and R324 were each mutated to alanine, C-terminally fused to msGFP2, and tested for effect on *A. baumannii* colony formation after allele swap, alongside reference alleles (*advA* WT and 1-230). Colony growth was assessed on solid Ara 0.5% medium with or without CIP (0.1 µg/ml). (C-D) Division and subcellular localization defects of AdvA^R324A^. Gene swap strains producing the indicated AdvA-msGFP2 construct were cultured with Ara 0.5% (no CIP) for 3 h and analyzed by phase-contrast and fluorescence microscopy (n ≥ 260 cells per strain, pooled from 3 independent cultures). (C) Analysis of cell lengths as in Fig. 1D. One data point for WT allele is not shown (length > 15 µm). **** P<0.0001, Mann-Whitney test. (D) Representative micrographs and demographs of subcellular fluorescence. Scale bar, 4 µm. (E) Effect of R324A on AdvA interaction with ZipA tested via BACTH assay. AdvA variants (left labels) and ZipA were fused in both orientations to CyaA fragments. Images representative of n=3 independent cultures (B, E).

### Structural modeling supports interaction of the AdvA cyclase-like domain with ZipA, with predicted contacts mapping to the charged tip and terminal helix

We used computational structural modeling with AlphaFold3 to gain insight into how AdvA binds ZipA in the cytoplasm. The prediction of the complex between AdvA and ZipA cytosolic domains was of high confidence, with predicted template modeling (pTM) score of 0.92, interface pTM (ipTM) score of 0.89, and low minimum PAE (1 Å) (Fig. 9). Strikingly, the predicted interface centered on the positively charged tip and α6 helix of AdvA, the two key features of its cyclase-like domain. In the model, R324 and other positively charged tip residues are predicted to form salt bridges with acidic residues in ZipA, while the α6 helix makes contacts along part of the ZipA disordered linker (Fig. 9). This analysis lends further support for the AdvA-ZipA interaction and highlights potential functions for R324 and the essential α6 helix in interfacing with an early component of the divisome.

**Fig. 9.**
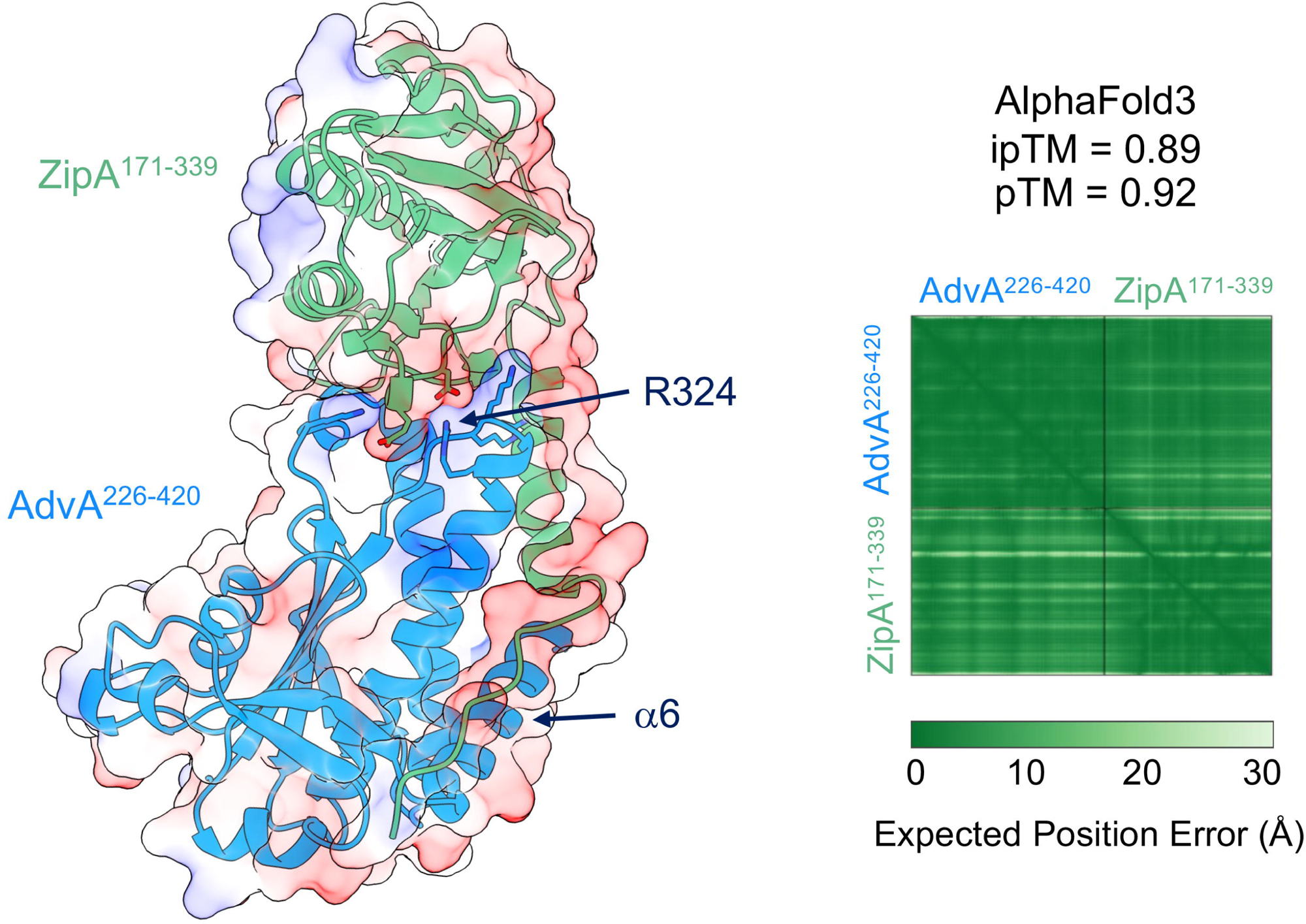
Structural prediction of the interface between the AdvA cyclase-like domain and ZipA. AlphaFold3 structural model of the cytoplasmic AdvA-ZipA complex. Ribbon diagrams for *A. baumannii* AdvA^226-420^ and *A. baumannii* ZipA^171-339^ are given in light blue and green, respectively (left). Side chains are shown for basic residues in the conserved tip of AdvA and acidic residues in the interacting surface of ZipA, and R324 and α6 of AdvA are labelled. Surface overlays show electrostatic potential (blue, positive charge; red, negative charge). ipTM and pTM scores and plot of the predicted alignment error (PAE) showing the predicted uncertainty in Å for all residue pairs are given at right. Low PAE (dark green) indicates highly confident relative domain positioning.

In addition to ZipA, we performed AlphaFold3 modeling of the interactions between the AdvA N-terminal region and candidate inner-membrane/periplasmic divisome hits identified in BACTH screens (FtsN and FtsL). FtsN was modeled with or without its N-terminal SPOR domain, and FtsL was modeled together with FtsB, as the two proteins form a co-dependent heterodimer in *E. coli* [42, 43]. In contrast to ZipA, the interfaces between AdvA and these proteins were not confidently modeled, showing low ipTM scores (≤0.25) and high interface PAE (≥8 Å), while internal control interfaces in the same predictions (the AdvA N-terminal dimer and the FtsL-FtsB heterodimer) were of higher confidence (Table S4). We did not model FtsA, since the filament-forming, membrane-tethered structure of the actin homolog is not well represented by AlphaFold3. Together, these observations support the above notion that the BACTH signals between membrane proteins should be regarded as candidate interactions requiring additional confirmation. Further, they distinguish the membrane-region signals from the AdvA-ZipA interaction, which is supported by both structural modeling and mutational analysis.

### Sequence and structural comparisons reveal close and distant AdvA homologs

BLASTp analysis of the full-length protein sequence revealed that closely related homologs of *A. baumannii* AdvA are restricted to the Moraxellaceae family of γ-proteobacteria (Fig. 10A). As expected, orthologs with highest similarity were in *Acinetobacter* (sequence identity ranging from 51-97%), while other genera, including *Psychrobacter* and *Moraxella*, showed more divergence (22-41% identity with *A. baumannii* AdvA). Regarding the *Acinetobacter* AdvA orthologs, a stretch of low sequence conservation and/or gaps was seen in the linker connecting the CC and CHD-like domains (between residues 210-230), which corresponds well to the *advA* sites enriched in viable transposon insertions (Fig. 10A). In accord with the previous analysis of the CHD-like domain, virtually all of the close orthologs (126/128) had arginine at AdvA residue position 324, and multiple α6 helix residues were conserved (Fig. 10A). The 13 most C-terminal residues of AdvA predicted to form an unstructured tail after α6 were either poorly conserved or absent from the vast majority of orthologs.

**Fig 10.**
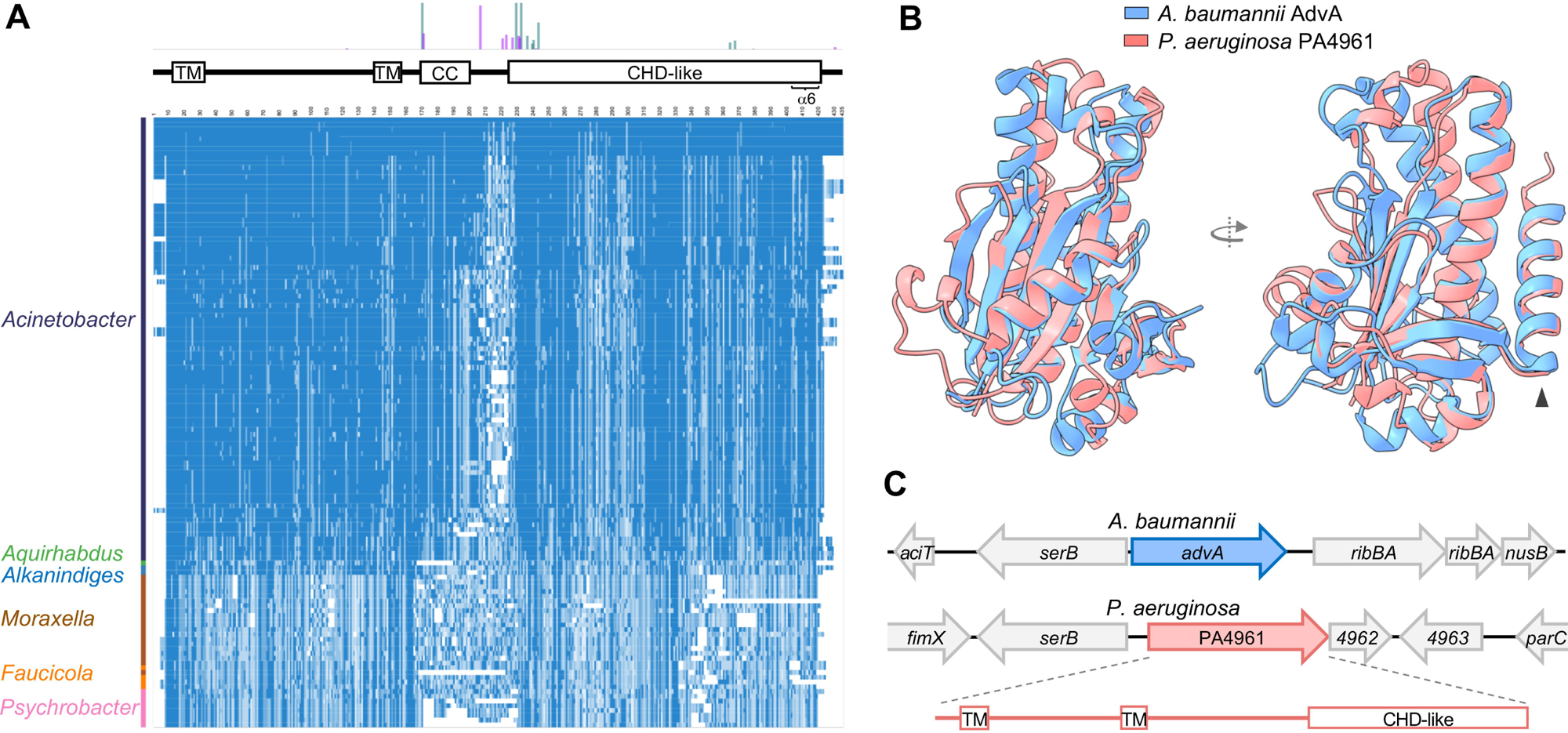
Analysis of AdvA homologs. (A) Heatmap shows amino acid sequence similarity of close AdvA homologs. *A. baumannii* AdvA is the first row, followed by sequence homologs from Refseq prokaryote representative genomes, using BLASTp with filters E<0.01, coverage 75-100. Taxon is indicated by label at left. Within heatmap, blue shading represents BLOSUM similarity score (darker blue, higher score/conservation), and white indicates gap. AdvA protein diagram and *advA* Tn-seq mutant abundance (as in Fig. 1A) are shown above the heatmap. (B) Structural alignment of AdvA^226-420^ (PDB: 7SUA) with AlphaFold model of the CHD-like domain of *P. aeruginosa* PA4961 (residues 324-512; pTM 0.91). Arrowhead denotes structurally conserved C-terminal helix. (C) Genomic regions of *A. baumannii advA* and *P. aeruginosa* PA4961. PA4961 predicted protein features are shown below the gene map.

To identify possible remote homologs of AdvA beyond the Moraxellaceae not evident through sequence comparisons, we performed an expanded search for proteins whose structures align with the AdvA^226-420^ crystal structure. We required that candidate structural homologs contained a C-terminal helix matching that of AdvA and lacked the dimerization arm of AC/GCs. As no protein in the PDB passed these criteria, we used Foldseek to search the AlphaFold Protein Structure Database [44]. This led to the identification of PA4961 from *P. aeruginosa* PAO1. PA4961 is an uncharacterized protein annotated as a histidine kinase (HK). The predicted structure of the C-terminal segment of the protein (residues 324-512) significantly matches AdvA^226-420^ (Fig. 10B, RMSD of 1.115 Å for 78 pruned atom pairs and 4.995 Å across all 172 pairs) and resembles a CHD, not a HK catalytic domain. Like AdvA, PA4961 has the unusual property of lacking the canonical AC/GC catalytic residues and dimerization arm while containing an extra C-terminal α-helix. Overall, PA4961 and AdvA have low amino acid sequence identity (12%) but have the same predicted TM topology with analogous periplasmic domains, and both genes share a genomic context oriented divergently from *serB* (Fig. 10C).

Interestingly, although few studies have examined PA4961, genome-wide screens have linked mutations in the locus (or its PA14 equivalent PA14_65570) to defective fitness in bacteriological medium [45], increased antibiotic susceptibility [46, 47], and cell filamentation [48]. AdvA and PA4961 are thus likely to be distant homologs and examples of a new type of bacterial control protein employing a CHD-like fold.

## Discussion

In these studies, we have defined key functional and structural features of AdvA, a distinctive essential protein that bears no sequence resemblance to known proteins and is critical to *A. baumannii* cell division. Our Z-ring recruitment and bypass suppressor data support the conclusion that AdvA essentiality is based on at least two key functions, assembling and helping activate the multicomponent divisome in the microorganism. We revealed by truncation analysis that AdvA contains N- and C-terminal regions with different contributions to *A. baumannii* viability and divisome interactions. The N-terminal periplasmic and TM domains are essential for viability, mediate dimerization, and show candidate interactions with early and late divisome proteins. By contrast, the cytoplasmic C-terminal domain is largely dispensable for viability but required for efficient cell division and fluoroquinolone resistance, and interacts with the early Z-ring protein ZipA. Removing a small segment from the end of this domain, however, had the striking effect of blocking AdvA divisome interactions and cell division. These results suggest that the cytoplasmic domain can modulate the activities of the protein, potentially through conformations that recapitulate effects of truncations. To investigate its control mechanisms, we solved the crystal structure of the cytoplasmic domain, which revealed a 3D fold resembling AC/GC enzymes with crucial differences—absence of canonical active sites and presence of atypical structural elements essential to division and CIP susceptibility. These results thus link cell division functions of AdvA to unique features of its structure.

Our integrated data support a model for the critical role of AdvA in *A. baumannii* divisome assembly (Fig. 11). (i) AdvA self-associates through its N-terminal region (supported by two-hybrid and structural modeling) and appears to function as a dimer. (ii) AdvA localizes to Z-rings early in divisome formation likely via recruitment by ZipA (although interaction with FtsZ has not been ruled out). The AdvA CHD-like domain makes key cytoplasmic contacts with ZipA through its positively charged tip (including R324) and α6 helix. The domain lacks canonical catalytic sites but retains the overall cyclase fold, which could permit binding of a cytoplasmic ligand that influences AdvA function. (iii) Detection of ZipA by the CHD-like domain may promote recruitment of additional divisome proteins. This recruitment could directly involve AdvA through its N-terminal domains, or, alternatively, AdvA may indirectly trigger recruitment by modulating FtsA through the ZipA interaction. By analogy with *E. coli* [10, 49], modulation of FtsA may drive recruitment of subsequent divisome proteins that promote synthesis of the septal wall. (iv) Suppressor mutations affecting septal synthesis proteins FtsB and FtsW bypass AdvA essentiality and to a large degree rescue the AdvA^-^cell division defect. These mutations have analogy with similar *E. coli* suppressors in FtsB and FtsW that hyperactivate division [25, 27, 28]. Because they likely lead to a hyperactive septal peptidoglycan synthase, these mutations could bypass AdvA either by substituting for a checkpoint or activating function normally provided by AdvA, or simply by driving division sufficiently to rescue a divisome crippled by AdvA deficiency. The specificity of the suppressors for AdvA and ZipA, but not FtsK, argues against the latter possibility of nonspecific divisome rescue. Further work is needed to define a possible role for the ZipA-AdvA interaction in divisome control, and to resolve the broader molecular contacts governing activation of septal synthesis in *A. baumannii*.

**Fig 11.**
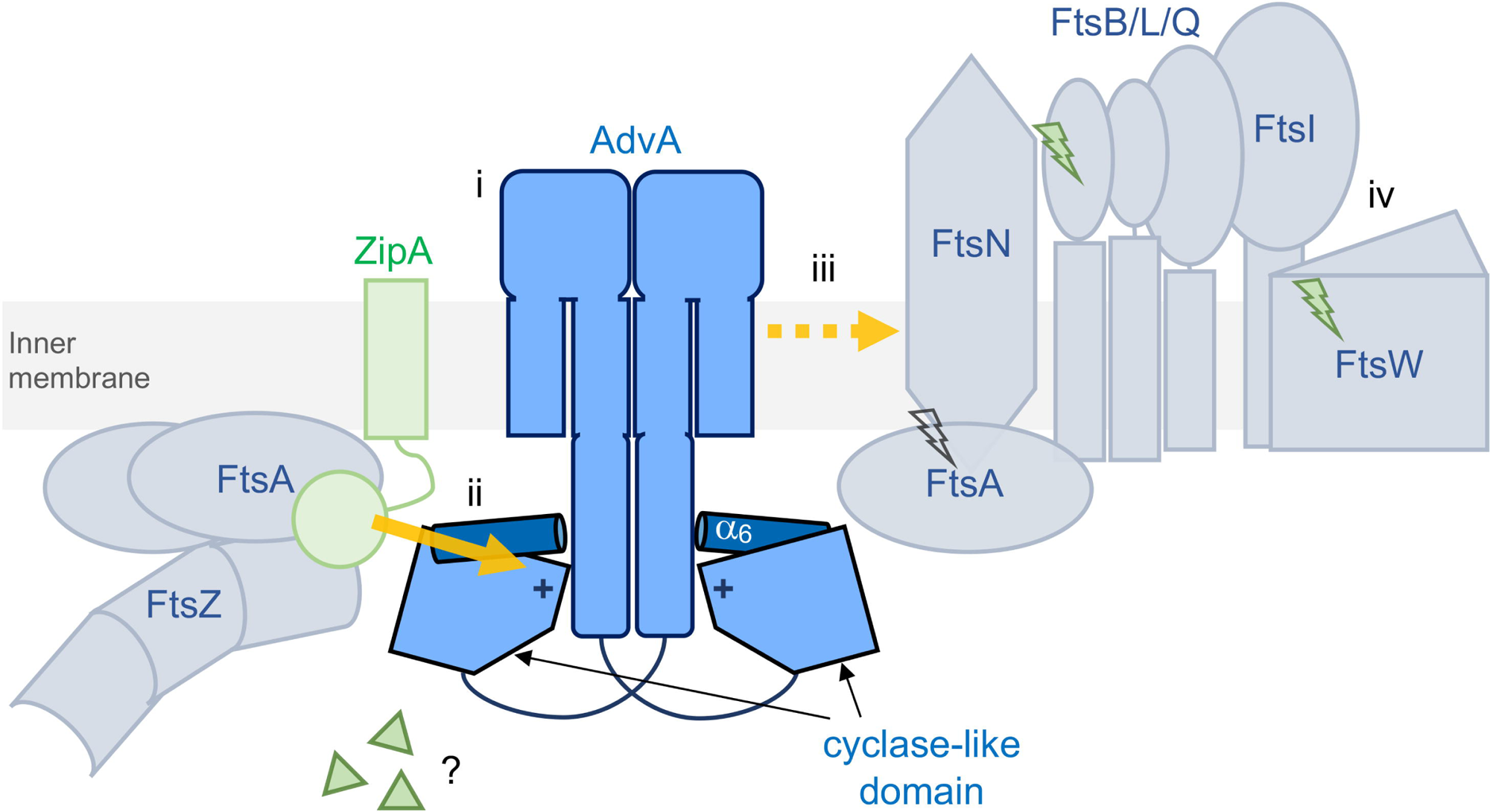
Model for AdvA-dependent divisome formation. Model depicts roles of AdvA in divisome construction and control to drive cell division independent of FtsEX, which is absent from *Acinetobacter*. (i) AdvA acts as a dimer. (ii) The protein is likely recruited early to midcell through binding ZipA (yellow arrow). ZipA binding is mediated by the positively charged tip (+) and C-terminal α6 helix (dark blue) of the cytoplasmic AdvA CHD-like domain. AdvA variants lacking the CHD-like domain still show some midcell localization, albeit inefficient, suggesting that additional contacts may contribute to AdvA localization. The cyclase-like structure raises the possibility that other cytoplasmic signals such as nucleotide-related molecules (triangles) may modulate AdvA CHD function. (iii) AdvA enables late divisome assembly. Detection of ZipA by the AdvA CHD-like domain may favor conformations of the rest of the AdvA protein, or of ZipA, that promote recruitment of additional divisome components (yellow dotted arrow). Candidate interactions with late divisome components require experimental confirmation, and other potential targets of the AdvA periplasmic domain related to division are possible. (iv) Divisome assembly ultimately leads to activation of the FtsWI septal synthase. Suppressor mutations in FtsB and FtsW identified in this study (green lightning) bypass loss of AdvA, suggesting that AdvA may be involved in the activation pathway. An AdvA bypass suppressor in FtsA was also identified in [22] (open lightning). The terminal α-helix of AdvA (dark blue) is essential for AdvA function and cell division, likely by stabilizing a state competent for dimerization, divisome protein interactions, and Z-ring recruitment.

Overall, AdvA’s properties of driving divisome assembly in the WT and being bypassed by mutations affecting septal synthases bear resemblance to features of the widely conserved FtsEX system, which is absent from *Acinetobacter*. Notably, all of the taxa harboring close AdvA orthologs (Fig. 10A) also lack FtsEX. These data are consistent with the possibility that AdvA performs functions substituting for the system, albeit through different molecular interactions, since AdvA binds ZipA, which is not a known FtsEX partner. More work is needed to define the extent to which AdvA may replace FtsEX functions in divisome assembly and activation, as well as determine whether AdvA has a role, like FtsEX, in regulating septal cell wall hydrolysis [14, 17].

*A. baumannii* cell division appears to hinge on the critical α6 helix of the AdvA cytoplasmic domain, based on the dramatic phenotypes of the 1-403 mutant (blocked homodimerization, divisome interactions, midcell localization and ultimately division). It is intriguing that the α6 region has such global effects, when two-hybrid results indicate the entire cytoplasmic domain binds just one core divisome protein (ZipA) and the domain is predicted to provide minimal self-interaction surface. We propose a two-fold explanation for the striking defects of AdvA^1-403^. First, the CHD-like domain may contain an inhibitory function that is normally suppressed by α6; for instance, α6 could act as a “latch” between AdvA subunits, enabling the remainder of the protein to stably adopt a dimeric state competent for its divisome functions. The inability of AdvA^1-403^ to form dimers may be the basis for most of the binding defects of the protein, if binding requires dimeric AdvA surfaces or conformations. Second, independent of dimer formation, α6 may be critical for interaction with ZipA and thus for AdvA recruitment to midcell division sites. This idea is supported by the two-hybrid data showing that ZipA interacts with AdvA^165-435^ but not AdvA^1-403^, despite neither having the ability to homodimerize (Figs. 5 and 7), and by the structural model (Fig. 9). Notably, the critical requirement in *A. baumannii* for the C-terminus of AdvA containing α6 helps to explain the unusual positional effects of transposon insertions in *advA* on Tn-seq fitness [15]. Insertions after the possibly inhibitory sites in the CHD-like domain producing truncations lacking α6 would likely yield globally inactive forms of the protein mimicking AdvA^1-403^ and blocking division; whereas insertions in the linker upstream of the domain would avoid such inhibitory latchless cytoplasmic fragments, yielding dimeric N-terminal domains that provide hypomorphic binding functions.

We have uncovered a highly divergent variant domain related to the type III NC family, with a core architecture most closely resembling the CHD of AC/GC enzymes but without the typical catalytic residues, dimerization arm, or topology. Combined with the lack of AC/GC activity in multiple assays, our data strongly suggest that the AdvA domain is not an AC or GC. Diguanylate cyclases (DGCs) are also members of the type III NC family, containing the same core fold as AC/GCs while lacking a dimerization arm [35, 50], and we considered a possible relationship with these enzymes. AdvA bears lower resemblance to DGCs than AC/GCs, however, with fewer aligning residues (120 vs 160) and lower DALI Z-score (10.9 vs 16.3) when comparing the closest DGC and AC/GC matches to AdvA. The absence of a GGDEF motif or PPi release from nucleotides *in vitro* further suggests that AdvA is not a DGC. AdvA^158-435^ also lacks appreciable NTPase activity *in vitro*, at least under the conditions tested, together raising the possibility that the protein uses mechanisms independent of NTP hydrolysis. By contrast, our mutational data indicate the domain is critical to division efficiency and can globally modulate AdvA self and divisome interactions. Why would *Acinetobacter* evolve a variant NC protein for such a role? The domain may have a yet unidentified enzymatic activity, in the context of the complete protein, that influences divisome interactions. Alternatively, as suggested above, AdvA may use the CHD-like fold to sense nucleotide-related ligands important for control of the *A. baumannii* cell cycle, or to exploit interactions with proteins that are traditional binding partners of ACs, such as G-proteins [35]. Experiments examining these alternative mechanisms are in progress.

Our results show that *A. baumannii* relies on a divergent NC-related architecture to enable a distinct strategy for controlling cell division. The CHD-like fold may be used by other pathogens including *P. aeruginosa*, which possesses a distant AdvA ortholog also linked to antibiotic resistance and cell division. Interestingly, unlike taxa with close AdvA orthologs, *P. aeruginosa* employs the widespread FtsEX system in its division cycle [51]. How the two pathogens have evolved to use AdvA-family proteins in distinct ways is a key question for future studies. The essential function of the AdvA periplasmic domain is also unknown. This region is predicted by AlphaFold3 to adopt a PAS-like fold (Fig. S8C), a module often mediating sensing or protein interactions. Defining the potential interactions of this domain is an additional question for future work, as is resolving the conformational dynamics of AdvA involving the cyclase-like domain during the division process. Improving our understanding of atypical division proteins such as AdvA may inform strategies for blocking division and enhancing antibiotic susceptibility of MDR nosocomial pathogens.

## Materials and methods

### Bacterial strains, growth conditions, and antibiotics

Bacterial strains used in this work are described in Table S1. *A. baumannii* strains were derivatives of ATCC 17978. *Escherichia coli* strains were DH5α, DH5α(λpir), or XL1-Blue for cloning; BTH101 for BACTH; and BL21[DE3] for protein purification and AC/GC reporter assay. All cultures were in lysogeny broth (LB) (10g/L tryptone, 5g/L yeast extract, 10g/L NaCl) at 37℃ unless otherwise indicated. Antibiotics were used for isolation of strains with plasmids as follows: tetracycline (Tc, 5-10µg/ml), kanamycin (Km, 10µg/ml), carbenicillin (Cb, 50µg/ml) for *A. baumannii*; Tc (10µg/ml), Km (30-50µg/ml), Cb (50-100µg/ml), ampicillin (Amp, 50-200 μg/ml), and/or chloramphenicol (Cm, 25µg/ml) for *E. coli*. Liquid cultures were aerated by shaking in flasks or by rotation on a tube roller drum. Medium included 100µM IPTG for maintaining P(IPTG)-*advA* strains and with 5-10% sucrose for maintaining the *advA*(1-230) strain.

### Molecular cloning and strain construction

Plasmids and synthetic DNA oligos/gene fragments used in this study are listed in Tables S2 and S3, respectively. Oligos and gBlocks were purchased from IDT. Blunt-ended PCR products (Q5 High-Fidelity DNA polymerase, NEB) and gBlocks were cloned into the HincII site of pUC18 and sequenced (Genewiz, Quintara Biosciences) before subcloning. Plasmids were introduced into *A. baumannii* by electroporation.

#### Allelic exchange

For chromosomal P(IPTG)*-advA*, a ∼3kb fragment including *lacIq-*P_T5*lac*_*-advA* was amplified from pAFE43 with 5’ SphI and 3’ SalI sites, and ∼1kb sequence upstream of the *advA* ORF was amplified from gDNA with 5’ NotI and 3’ SphI sites; the fragments were digested and fused via 3-way ligation into NotI and SalI sites of pSR47S, resulting in pAFE104. For chromosomal *zapA-mCherry*, a ∼800bp fragment including *A. baumannii zapA* and upstream sequences was amplified from gDNA and subcloned in BamHI and XbaI sites of pAFE25, creating *zapA-mCherry* flanked by BamHI and PstI sites; a ∼1kb fragment downstream of the *zapA* ORF was amplified from gDNA and cloned using 5’ PstI and 3’ SalI sites; the BamHI-PstI and PstI-SalI fragments from the resulting plasmids were 3-way ligated with BamHI-SalI cut pSR47S, creating pAFE324. pAFE25 was constructed by cloning a synthetic gene fragment harboring 5’ XbaI and 3’ PstI sites in the HincII site of pUC18 in the direction of *lacZ*α. For chromosomal *advA-GFP*, *advA-GFPmut3* was amplified from pAFE42 with 5’ BamHI and 3’ PstI sites; ∼1kb of sequence downstream of *advA* was amplified from gDNA with 5’ PstI and 3’ SalI sites; the fragments were digested and fused via 3-way ligation into BamHI and SalI sites of pSR47S resulting in pAFE65. To re-introduce *ftsW* or *ftsB* alleles, ∼2kb of sequence centered on each mutation was amplified from mutant gDNA with flanking BamHI and SalI sites and subcloned in pSR47S resulting in pAFE185 and pAFE186, respectively. For chromosomal *advA*(1-230), ∼2kb of partial *advA* and 5’ flanking sequence was amplified from gDNA with 5’ NotI and 3’ PstI sites and a stop codon following codon 230; ∼1kb of sequence 3’ of the *advA* stop codon was amplified with 5’ PstI and 3’ SalI sites; and fragments were 3-way ligated with pSR47S via NotI and SalI sites, creating pAFE272. Allelic exchange was performed using the sucrose counterselection method and verified by colony PCR [52].

#### GFP fusions

A Tc^r^ vector for Ara gene induction (pYDE344) was constructed by cloning the NsiI-EcoRI fragment from pVRL2 into pYDE153, such that *araC*-P_BAD_ replaces the *lacI-*P_T5lac_. (i) To construct *advA*-msGFP2 under Ara control, *advA* fragments were generated with 5’ BamHI and in-frame 3’ XbaI sites and subcloned in pAFE225. (ii) Deletions were constructed by PCR using the WT plasmid (pAFE229) as template. To delete parts of the AdvA C- or N-terminus, we replaced the full-length fragment with truncated PCR products using distinct 3’ or 5’ primers, respectively, the latter containing the Shine-Dalgarno and ATG sequence from pFPV25. For internal deletion, we used inverse PCR with primers containing a glycine codon and SpeI sites, followed by self-ligation via SpeI, such that Gly-Thr-Ser replaced the periplasmic domain. (iii) Point mutants were cloned with gBlocks. (iv) For untagged constructs, fragments were re-amplified using distinct primers with EcoRI (5’) and stop codon and PstI sites (3’). (v) For divisome proteins C-terminally fused to msGFP2, gene fragments were amplified from gDNA and subcloned into pAFE225. (vi) To make divisome proteins N-terminally fused to msGFP2, fragments were amplified from gDNA with 5’ SpeI and 3’ PstI sites and subcloned to pAFE309 (pAFE309 constructed by PCR-amplifying *msGFP2* using pAFE225 template and primers with 5’ SacI and 3’ SpeI sites and cloning the blunt-ended product in the HincII site of pUC18 in the direction of *lacZ*α). From the resulting plasmids, the EcoRI-PstI fragments (i-v) or SacI-PstI fragments (vi) were subcloned in pYDE334.

#### Reporter assay plasmids

For BACTH assay, genes were amplified from gDNA with 5’ SphI and 3’ XbaI sites (C-terminal fusions) or 5’ BamHI and 3’ EcoRI sites (N-terminal fusions), and constructs were subcloned into either pKNT25 and pUT18 (C-terminal fusions) or pKT25 and pUT18C (N-terminal fusions). BACTH plasmids were maintained in *E. coli* XL1-Blue at 30℃ before cotransforming *E. coli cya* strain BTH101. For AC/GC assay, genes were amplified from gDNA with 5’ NdeI and 3’ SalI or HindIII sites. Constructs were subcloned in pET23(a)+ before transforming *E. coli* CRP/CRP_G_ reporter strains.

### Western blot analysis

Cells were grown to optical density (OD) ∼1, and 1 OD unit was harvested by centrifugation. Pellets were resuspended in 50µl Laemmli buffer and heated at 95℃ for 5 min. Proteins were separated by 10% PAGE and transferred to low-fluorescence PVDF membranes. SYPRO Ruby blot stain (Invitrogen) was used for visualizing total protein following the manufacturer’s protocol. Blots were blocked (5% milk in TBST) and incubated with primary rabbit anti-GFP antibody (Invitrogen A11122, 1:2000) or rabbit AdvA^158-435^ antiserum (1:10,000) followed by secondary goat anti-rabbit IgG, HRP (Invitrogen 65-6120, 1:5000); or primary mouse CyaA antibody 3D1 (Santa Cruz Biotechnology SC-13582, 1:200, targeting CyaA T18) followed by secondary rabbit anti-mouse IgG, HRP (A27025, 1:5000). Blots were developed with Western Lightning Plus (Perkin Elmer) and imaged with a ChemiDoc MP (BioRad). Rabbit polyclonal AdvA antiserum was produced at the Pocono Rabbit Farm and Laboratory by immunization with purified recombinant AdvA^158-435^ using standard protocols.

### Colony formation tests

Bacteria were precultured for 3 h in standard or permissive LB medium (containing 100µM IPTG for gene swaps, or 5% sucrose for *advA*(1-230)) and washed 2x with PBS. Serial 10-fold dilutions in PBS were spotted on the noted solid medium. Colonies were imaged after overnight growth with transilluminated light on a ChemiDoc MP (BioRad).

### Microscopy

Gene swap and divisome tracking strains were precultured in LB with 100µM IPTG and washed as above, seeded at OD 0.02 in LB with 0.5% Ara, and imaged at 1 and 3 h (gene swap). For divisome tracking, cultures were seeded first in LB with 5% sucrose, incubated for 3 h, re-seeded at OD 0.02 in LB containing 5% sucrose and 0.5% Ara, and cells imaged after 1 h.

CRISPRi strains were precultured in LB for 3 h, seeded at OD 0.02 in LB with 50ng/ml aTc, and imaged after 3 h culture. A sample was mixed 1:1 with Slow-Fade Diamond (Thermo Fisher) and immobilized on agarose pads (1% agarose in PBS). Micrographs were acquired with 100x/1.4 phase-contrast objective on a Zeiss Axio Observer 7 microscope. Fluorescence micrographs used a Colibri 7 LED light source with 475nm LED and filter set 92 HE for GFP, and 587nm LED and filter set 45 for mCherry. Fiji [53] was used to determine cell boundaries from stacked GFP/mCherry/phase images. Cell length was quantified from phase images using MicrobeJ [54]. To analyze fluorescence profiles, images were background-subtracted and quantified by using MicrobeJ. Results were exported to .mat and demographs generated using a custom MATLAB (R2022b) script. Cells were excluded from analysis if their mean fluorescence was outside of the range defined by the lower quartile minus 1.5 times interquartile range (IQR) and the upper quartile plus 1.5 times IQR. Medial axis fluorescence profiles were then arranged by cell length and centered on Y=0.

### Bacterial two-hybrid assay

BACTH assays followed described protocols [30]. Overnight cultures of BTH101 cotransformants were spotted on solid BACTH assay medium (LB 1.5% agar [LBA] with Km 30µg/ml, Cb 50µg/ml, IPTG 500µM, and X-gal 40µg/ml). Plates were incubated overnight at 30°C followed by 24 h at room temperature and reporter color was imaged using dark-field illumination.

### Suppressor isolation and mapping by whole-genome sequencing

Colonies of AFA29 on LBA with 1mM IPTG and 5 µg/ml Tc were directly streaked on LBA plates or were inoculated into parallel tubes of LB, precultured 3h, and spread on multiple LBA plates. After overnight incubation, colonies were purified in parallel on LBA with or without 5 µg/ml Tc and incubated overnight. For isolates maintaining Tc^r^ (marker of pEGE305 providing extra copy of *lacI*) colonies from the LBA plate were inoculated into LB. The parent strain was also inoculated in LB with or without 100µM IPTG to provide reference controls.

When cultures reached OD of 0.3, samples were analyzed by fluorescence microscopy. For suppressor candidates that maintained a low GFP signal similar to the uninduced parent, cultures were continued until they reached saturation, stocked, and gDNA extracted (DNeasy kit Qiagen). gDNA was quantified by SYBR green assay. Illumina libraries were prepared using a modified small-volume Nextera tagmentation method and sequenced at TUCF-Genomics (single-end 100 bp, HiSeq 2500) [55, 56]. Reads were mapped to NZ_CP012004 and mutations identified using breseq [57].

### Expression, purification and crystallization of *A. baumannii* AdvA

The gene encoding ACX60_00475 from *A. baumannii ATCC 17978* (residues 158-435) was cloned by ligation independent cloning as described previously [58, 59] into Amp^r^ pMCSG53 vector, containing an N-terminal 6×His tag followed by a TEV protease cleavage site and genes for rare codons. *E. coli* BL21(DE3)(Magic) cells [58] were transformed with the plasmid and the overnight starter culture was grown in LB supplemented with 130 μg/ml Amp and 50 μg/ml Km at 37°C and 220 rpm. The next day, 3 L of M9 medium (High Yield M9 SeMet media, Medicilon Inc.) supplemented with 200 μg/ml Amp and 50 μg/ml Km were inoculated at 1:100 dilution with the overnight culture and incubated at 37°C and 220 rpm.

Expression was induced at OD=1.8-2 with 0.5 mM IPTG and the culture was further incubated at 25°C and 200 rpm for 14 h. The cells were harvested by centrifugation (6,000 x *g*, 10 min), re-suspended (5 ml of buffer/1g of cells) in lysis buffer (50 mM Tris pH 8.3, 500 mM NaCl, 10% glycerol, 0.1% IGEPAL CA-630) and frozen at −30°C until purification.

Frozen pellets were thawed and sonicated at 50% amplitude, in 5 s x 10 s cycle for 20 min at 4°C. The lysate was cleared by centrifugation (18,000 x *g*, 40 min) at 4°C, the supernatant was collected. The protein was purified by IMAC and size exclusion chromatography on an ÅKTAxpress system (GE Healthcare) as described with some modifications [60]. The cell extract was loaded into a His-Trap FF (Ni-NTA) column in loading buffer (10 mM Tris-HCl pH 8.3, 500 mM NaCl, 1 mM Tris (2-carboxyethyl) phosphine (TCEP), and 5% glycerol) and the column was washed with 10 column volumes (cv) of loading buffer and 10 (cv) of washing buffer (10 mM Tris-HCl pH 8.3, 1M NaCl, 25 mM imidazole, 5% glycerol). Protein was eluted with elution buffer (10 mM Tris pH 8.3, 500 mM NaCl, 1 M imidazole), loaded onto a Superdex 200 26/600 column, separated in loading buffer, collected and analyzed by SDS-PAGE. 6xHis-tag was cleaved by recombinant TEV protease in ratio 1:20 (protease:protein) overnight at room temperature. The cleaved protein was separated from recombinant TEV protease, un-cleaved protein and 6xHis tag peptide by Ni-NTA-affinity chromatography using loading buffer followed by loading buffer with 25mM imidazole. The cleaved protein was collected in loading buffer with 25mM imidazole. Both fractions were analyzed by PAGE for purity and 6xHis tag cleavage.

The protein was concentrated to 8.8 and 16.6 mg/ml in loading buffer without NaCl and with 500 mM NaCl, respectively. Both protein samples were used to set up 2-μl crystallization drops (1μl protein:1μl reservoir solution) in 96-well plates (Corning) using commercial Classics II, PACT, and JSCG+ (QIAGEN) crystallization screens. Diffraction quality crystals grown from the condition with 0.1M Bis-Tris (pH 6.5) and 20% (w/v) PEG 5000 MME (Classics II, condition 46) were flash cooled in liquid nitrogen for data collection.

### Structure Determination, Refinement and Validation

Diffraction quality crystals were screened and several data sets were collected. The data sets were collected at beam line 21ID-F of the Life Sciences-Collaborative Access Team (LS-CAT) at the Advanced Photon Source (APS), Argonne National Laboratory. Images were indexed, integrated, and scaled using HKL-3000 [61]. The data set with the 1.65 Å resolution was used for structure solution and refinement. The structure was determined by the Single Anomalous Dispersion (SAD) method using anomalous signal from the Se-Met protein derivative. Crystal belongs to the orthorhombic space group P2_1_2_1_2_1_ with one protein chain in the asymmetric unit. The initial model (residues 227-419) was built using the HKL3000 structure solution package. Data quality, structure refinement and the final model statistics are shown in Table 1. The model was refined using REFMAC [62] and additional residues were manually added during visual inspections in Coot [63] to yield the final structure spanning residues 226-420. The water molecules were generated automatically using ARP/wARP [64], ligands (PEG and EDO) were fit into electron density maps manually and the structure was further refined in REFMAC. The final stages of refinement were carried out with the Translation–Libration–Screw (TLS) group corrections, which were generated by the TLS Motion Determination (TLSMD) server [65, 66]. The quality of the model during refinement and the final validation of the structure was done using MolProbity [67] (http://molprobity.biochem.duke.edu/). The structure was deposited to the PDB (https://www.rcsb.org/) with the assigned PDB code 7SUA. Structural alignments and molecular graphics procedures were performed with ChimeraX [68].

**Table.1.** Data quality and refinement statistics. Values in parentheses are for the outer shell.

| PDB Accession Code | 7sua |
| --- | --- |
| **Data Collection** |  |
| Space group | $P2_12_12_1$ |
| Unit cell parameters (Å; °) | $a = 50.76; b = 59.78; c = 75.63$<br>$\alpha = \beta = \gamma = 90.00$ |
| Resolution range (Å) | 30.00 - 1.65 (1.68 - 1.65) |
| No. of reflections | 27,958 (1,354) |
| $R_{\text{merge}}$ (%) | 6.9 (68.7) |
| Completeness (%) | 99.5 (99.0) |
| $\langle I/\sigma(I) \rangle$ | 24.3 (2.4) |
| Multiplicity | 4.3 (4.1) |
| Wilson $B$ factor | 22.7 |
| **Refinement** |  |
| Resolution range (Å) | 27.06 - 1.65 (1.70 - 1.65) |
| Completeness (%) | 99.2 (94.8) |
| No. of reflections | 26,532 (1,920) |
| $R_{\text{work}}/R_{\text{free}}$, (%) | 17.6/20.8 (26.5/24.6) |
| Protein chains/atoms | 1/1,664 |
| Ligand/Solvent atoms | 11/251 |
| Mean temperature factor (Å<sup>2</sup>) | 25.8 |
| **Coordinate Deviations** |  |
| R.m.s.d. bonds (Å) | 0.006 |
| R.m.s.d. angles (°) | 1.365 |
| **Ramachandran plot** |  |
| Favored (%) | 100.0 |
| Allowed (%) | 0.0 |
| Outside allowed (%) | 0.0 |

### Enzyme assays

AdvA^158-435^ was also purified without cleavage of tag and without SeMet for enzyme assays. *E. coli* BL21 (DE3) pLysS cells were transformed with the pMCSG53-*advA*(158-435) plasmid; the overnight starter culture was in LB with Cm and Cb; and 500ml of Terrific Broth (TB) with 50µg/ml Cb and 25µg/ml Cm was used as protein production medium. After protein induction as described above, cultures were centrifuged and cell pellets frozen at -80°C. Pellets were thawed, resuspended in 5ml lysis buffer/1g of cells, and 15 µL benzonase (Sigma) was added per 50 mL to reduce viscosity. Cells were lysed by two passes through a French press, and lysates were clarified by centrifugation (16,000 x g, 1 h) at 4°C. Using gravity flow, 2.5 mL bed volume of Ni-NTA column was washed with 3 mL and then 10 mL of lysis buffer. Lysate was passed through column, followed by wash with 3x 10 mL of washing buffer. Protein was eluted with 5 mL of elution buffer with 250mM imidazole, then 5 mL with 500mM imidazole. The two elution fractions were merged, dialyzed against three changes of 1x loading buffer without TCEP at 4°C. After dialysis changes, TCEP was added to 1 mM final concentration. Protein aliquots were stored at 5.7 mg/ml concentration at -80°C.

NTP hydrolysis reactions were performed in 100 µL volumes containing NTP hydrolysis reaction buffer (25 mM HEPES, 150 mM KCl, 2 mM MgCl2, and 2 mM TCEP at pH 7.4.) and 0.5 mM NTP. NTP stocks (100 mM) were prepared by dissolving NTP powder (ATP, GTP, CTP, or UTP; Jena Bioscience) in 25 mM HEPES, pH 7.4. For measuring pyrophosphate release, 5 µM of 6xHis-tagged AdvA^158-435^ protein or equivalent volume of AdvA loading buffer was added to reactions. Reactions were mixed by pipetting, then incubated at 37°C for 1 h. Pyrophosphate was measured using the MAK168 fluorogenic PPi sensor kit (Sigma-Aldrich) per the manufacturer instructions, with PPi standards prepared in NTP hydrolysis reaction buffer.

Fluorescence was measured using a BioTek Synergy H1 microplate reader. For measuring phosphate release, reactions were prepared as above, except control reactions were set up with addition of 5 µM of apyrase (NEB). Reactions were diluted two-fold by addition of 100 µL reaction buffer and phosphate concentration measured using the Sigma-Aldrich Malachite Green Phosphate Assay Kit (MAK307) per the manufacturer instructions, with phosphate standard solutions prepared in NTP hydrolysis reaction buffer and absorbance at 620 nm detected on a BioTek Synergy H1 microplate reader.

To detect AC/GC activities of AdvA *in vivo*, the *E. coli* CRP/CRP_G_ reporter system was used as described [40]. *E. coli* BL21[DE3] *cya crp* strains harbored a CRP (the cAMP sensor CRP, or dual cAMP/cGMP sensor CRP_G_) and a heterologous NC, both IPTG-inducible from plasmids. NC controls included insertless vector (negative control), *A. baumannii* CavA (AC), and *Azospirillum* sp. B510 GuaA (GC). Colonies forming overnight on LBA containing Tc 10µg/ml and Cb 50 µg/ml were patched on reporter plates (LBA containing Tc 10µg/ml, Cb 50 µg/ml, X-gal 40 µg/ml, and IPTG at 25 or 50 µg/ml). Plates were incubated overnight at 37°C, followed by room temperature for one day, and imaged using white-light transillumination.

For ELISAs, strains were precultured in permissive medium (LB or LB supplemented with 5% sucrose) for 3 h, washed 2x with PBS, seeded in LB at OD 0.02, and cultured to OD 0.5. 1 ml of OD 0.5 was pelleted (3000xg, 3 min), resuspended with 1 ml 0.1 M-HCl-0.5% Triton X-100, and incubated for 10 min with end-over-end rotation at room temperature [69]. Samples were centrifuged (3000xg, 3 min) and supernatants were collected and stored at -20℃ prior to use. cAMP and cGMP ELISA kits (Enzo) were used per manufacturer’s specifications.

### Protein structural predictions and homology analysis

AlphaFold3 [37] was used for protein structural predictions. Dali [34] and Foldseek [70] were used for structural similarity searches. PDBsum was used to generate protein topology maps [71]. AdvA orthologs were identified using BLASTp [72] using query sequence WP_002000748.1, database refseq_select, maximum E value of 0.01, and minimum coverage of 75%. FastTree [73] was used to generate a phylogeny from the BLASTp hits. Custom scripts were used to assign BLOSUM62 matrix scores to positions in orthologs relative to WP_002000748.1 in an Interactive Tree of Life (iTOL)-compatible format. The phylogeny and heatmap data were visualized using iTOL [74].

## Acknowledgements

This work was supported by Northeastern University startup funds, Northeastern University PEAK Fellowships, the Dyson Foundation, and by HHS/NIH/NIAID grant R01AI162996 and contracts HHSN272201700060C and 75N93022C00035. This research also used resources of the Advanced Photon Source, a U.S. Department of Energy (DOE) Office of Science User Facility operated for the DOE Office of Science by Argonne National Laboratory under Contract No. DE-AC02-06CH11357. Use of the LS-CAT Sector 21 was supported by the Michigan Economic Development Corporation and the Michigan Technology Tri-Corridor (Grant 085P1000817). Access to LS-CAT is supported by the Northwestern Structural Biology Facility, which is funded in part by the Robert H. Lurie Comprehensive Cancer Center grant NCI P30CA060553. The content is solely the responsibility of the authors and does not necessarily represent the official views of the National Institutes of Health. This manuscript is the result of funding in whole or in part by the National Institutes of Health (NIH). It is subject to the NIH Public Access Policy. Through acceptance of this federal funding, NIH has been given a right to make this manuscript publicly available in PubMed Central upon the Official Date of Publication, as defined by NIH. We thank Mark Gomelsky for providing AC/GC reporter plasmids, Yunfei Dai for cloning the pYDE334 gene swap vector, and Jade Law, Abigail Thomas, Emma Young, Anika Fernandes, Erin Murphy, and Shantih Whiteford for technical assistance.

## Notes

### Competing Interest Statement

The authors have declared no competing interest.

### Summary of Updates

Updated and reordered figures; updated supplemental material including addition of Table S4; updated manuscript title and text.

