## Supplemental Figures and Tables for "Control of cell division by an *Acinetobacter baumannii* protein with a nucleotidyl cyclase-like domain"

### Table of Contents

Fig. S1. Evaluation of gene-swap system under control conditions
Fig. S2. Western blot analysis of AdvA variants in *A. baumannii*
Fig. S3. Additional analysis of AdvA truncations/deletions: (A) untagged versions behave identically to GFP-tagged constructs; and (B) non-complementing constructs lack dominant-negative effects
Fig. S4. Western blot analysis of AdvA depletion using the conditional P(IPTG)-*advA* allele
Fig. S5. Sucrose permits growth of bacteria with the conditional P(IPTG)-*advA* allele.
Fig. S6. Location and additional suppressor functions of amino acid substitutions in *A. baumannii* FtsB and FtsW that rescue AdvA$^-$ lethality
Fig. S7. Analysis of controls for two-hybrid assays
Fig. S8. Structural modeling analysis of predicted AdvA dimers
Fig. S9. Atypical topographical features of the AdvA CHD-like domain: lack of canonical AC/GC active site residues and presence of conserved tip
Fig. S10. AdvA CHD-like domain shows no detectable cyclase or NTPase activity under conditions tested *in vivo* and *in vitro*
Fig. S11. AdvA function is preserved despite substitutions to positions aligning with the canonical AC/GC active site

Table S1. Strains used in this study
Table S2. Plasmids used in this study
Table S3. Synthetic oligonucleotides and gene fragments used in this study
Table S4. AlphaFold3 confidence values for predicted AdvA assemblies

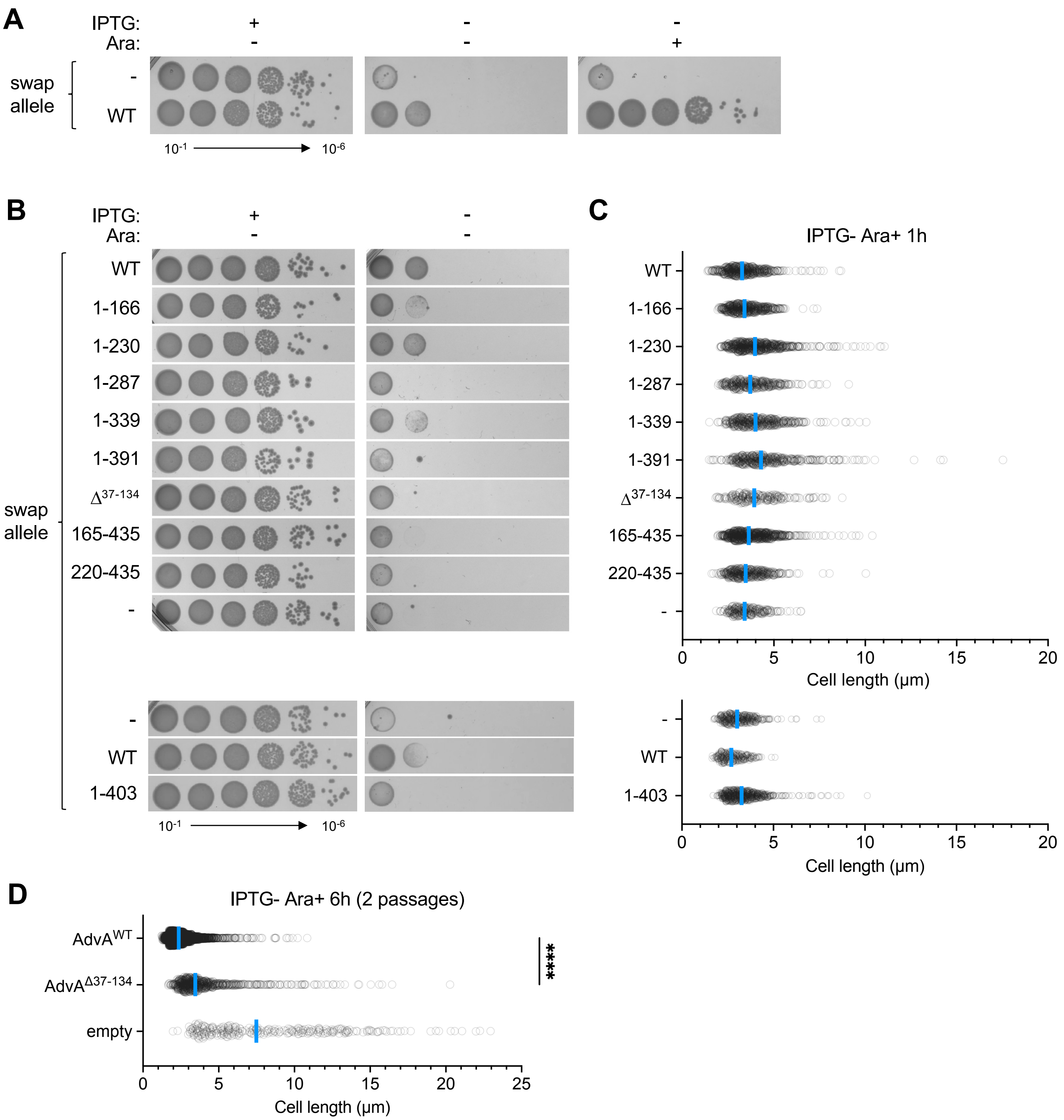


**Fig S1. Evaluation of gene-swap system under control conditions.** (A) Colony growth of *A. baumannii* AdvA allele swap strains harboring insertless P(Ara) vector or vector with WT *advA* swap gene. Solid medium contained IPTG 0 or 500 µM to control expression of the native *advA* gene, and Ara 0 or 0.5% to control expression of the plasmid-borne swap allele. (B) Colony growth of allele swap strains from Fig 1 on control medium (with or without IPTG 500 µM). Images representative of n=3 independent cultures. (C) Microscopy analysis of the *A. baumannii* AdvA swap strains from Fig 1 after 1 h in LB IPTG 0, Ara 0.5%. Cell length was determined from phase-contrast micrographs as in Fig 1 ($n \geq 234$ cells across two independent cultures per strain). Each symbol represents one cell; blue line, median; -, strain contained an insertless vector (no swap allele). (D) Cell length of the *A. baumannii* AdvA$^{\Delta 37\text{-}134}$ swap strain compared to controls after an additional passage in IPTG 0, Ara 0.5% swap medium (6 h total). After the first 3 h passage, cells were diluted in swap medium to OD 0.02 and grown an additional 3 h. Cell lengths were determined from phase-contrast microscopy ($n \geq 286$ cells per strain, pooled from 3 independent cultures; ****, $P<0.0001$, Kruskal-Wallis test.

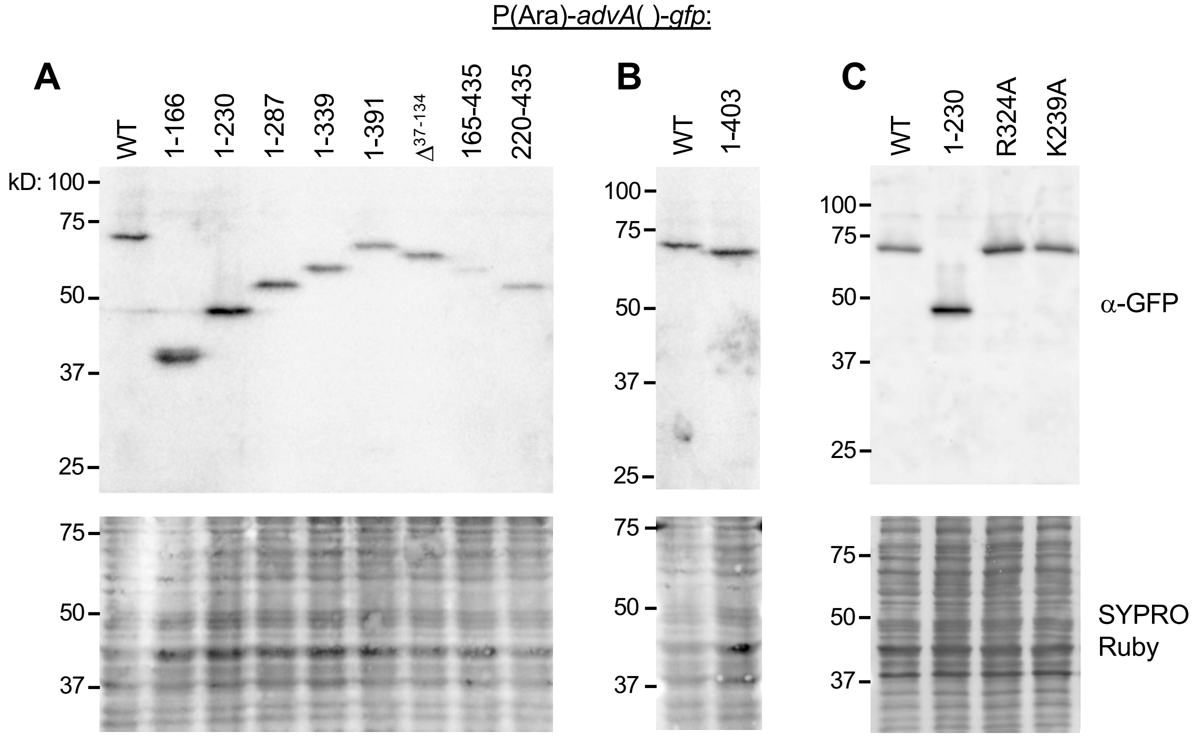


**Fig S2. Western blot analysis of AdvA variants in *A. baumannii*.** Gene swap strains harboring the indicated AdvA truncation/deletion construct (A, B) or point mutant (C) were cultured in LB Ara 0.5% as in Fig 1. Samples were collected at OD 1 and analyzed by SDS-PAGE and Western blot using anti-GFP primary antibody (top) and SYPRO Ruby to detect total protein (bottom).

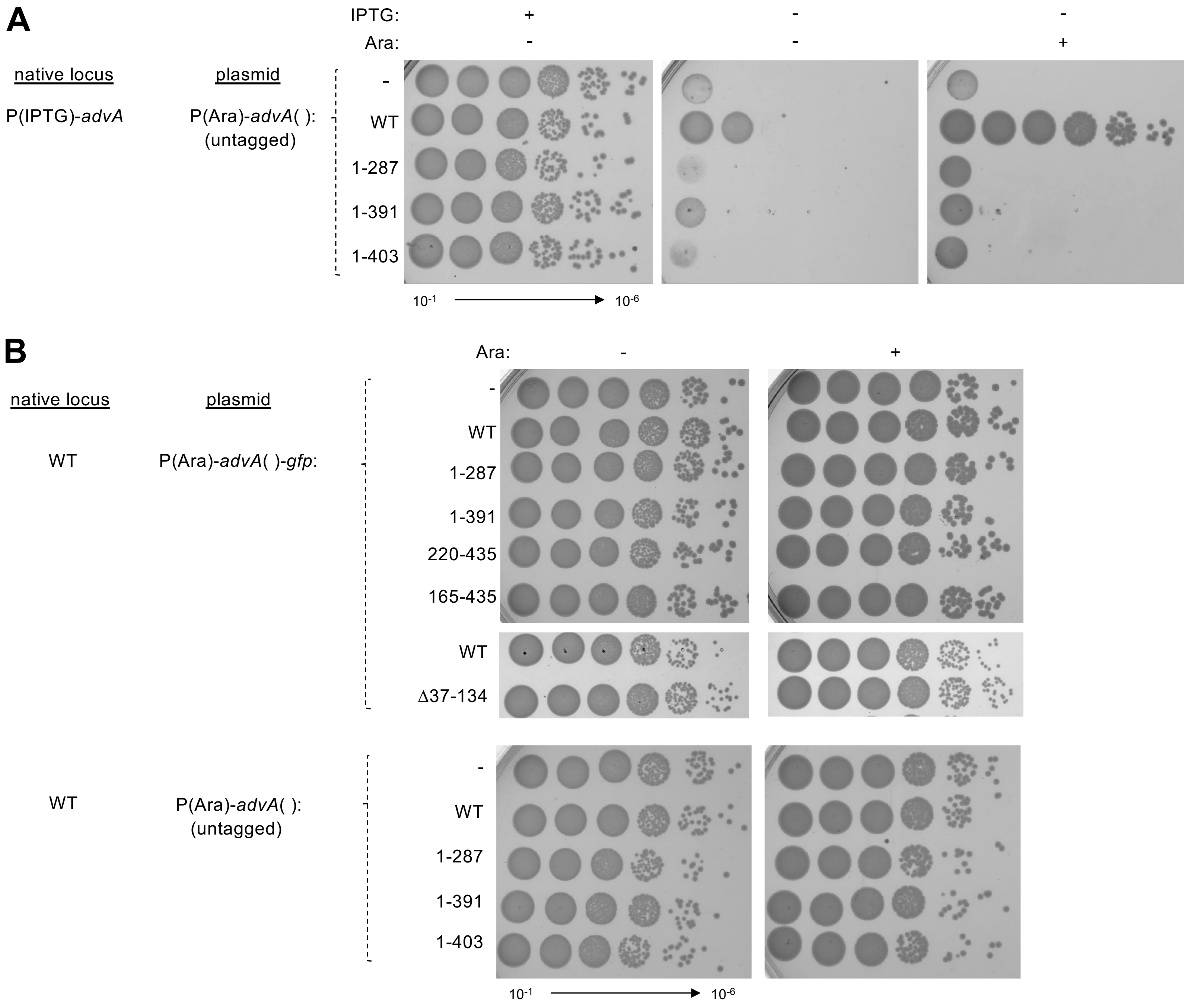


**Fig S3. Additional analysis of AdvA truncations/deletions: (A) untagged versions behave identically to GFP-tagged constructs; and (B) non-complementing constructs lack dominant-negative effects.** (A) Untagged versions of the non-complementing AdvA variants were tested in the *A. baumannii* gene-swap system for colony formation. Strains were precultured in IPTG 500 µM, dilutions were spotted on LBA with or without IPTG 500 µM or Ara 0.5%, and colony growth was analyzed as in Fig 1. (B) The AdvA-msGFP2 fusions from Fig 1 (top) and the untagged AdvA constructs from panel A (bottom) were tested in a WT (AdvA+) *A. baumannii* background for colony formation. Strains were precultured in LB, dilutions were spotted on LBA with or without Ara 0.5%, and colony growth was analyzed as in A. Images representative of n=3 independent cultures.

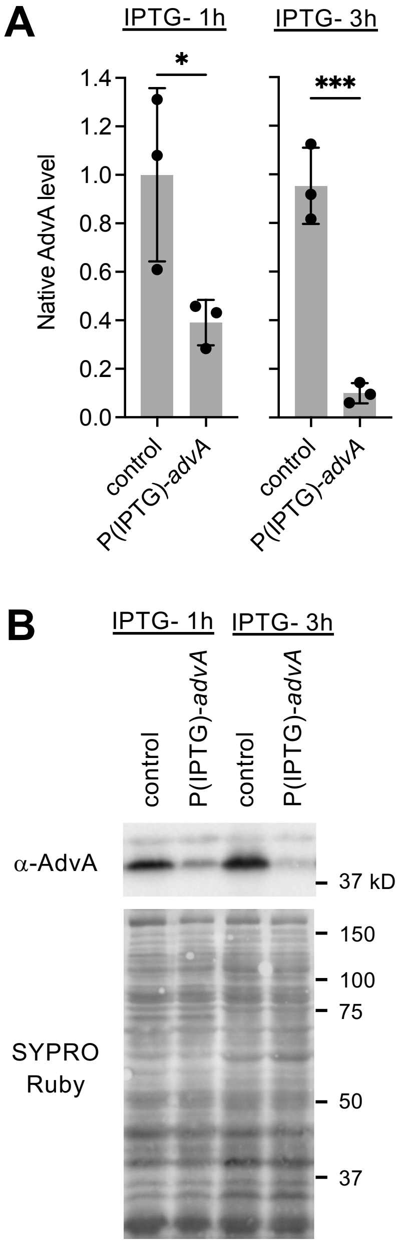


**Fig S4. Western blot analysis of AdvA depletion using the conditional P(IPTG)-*advA* allele.** AdvA protein levels were measured in *A. baumannii* strains harboring the conditional *advA* allele [P(IPTG)-*advA*] compared to the WT control. Strains also harbored an insertless plasmid vector (pYDE334), such that no AdvA-GFP swap construct is present. WT control was precultured in LB and P(IPTG)-*advA* was precultured in LB 100 μM IPTG for 3 h. Cells were washed twice with PBS and reinoculated into fresh LB. At 1 h and 3 h, 1 OD unit was collected and analyzed by SDS-PAGE and western blot using anti-AdvA antiserum. (A) Quantification of AdvA protein levels at 1 h and 3 h. AdvA levels were normalized by dividing AdvA band intensity by total protein intensity determined from SYPRO staining and are shown relative to control at 1 h. Bars show mean ± s.d. (n=3) * $P < 0.05$, *** $P < 0.001$, unpaired t test. (B) Representative western blot.

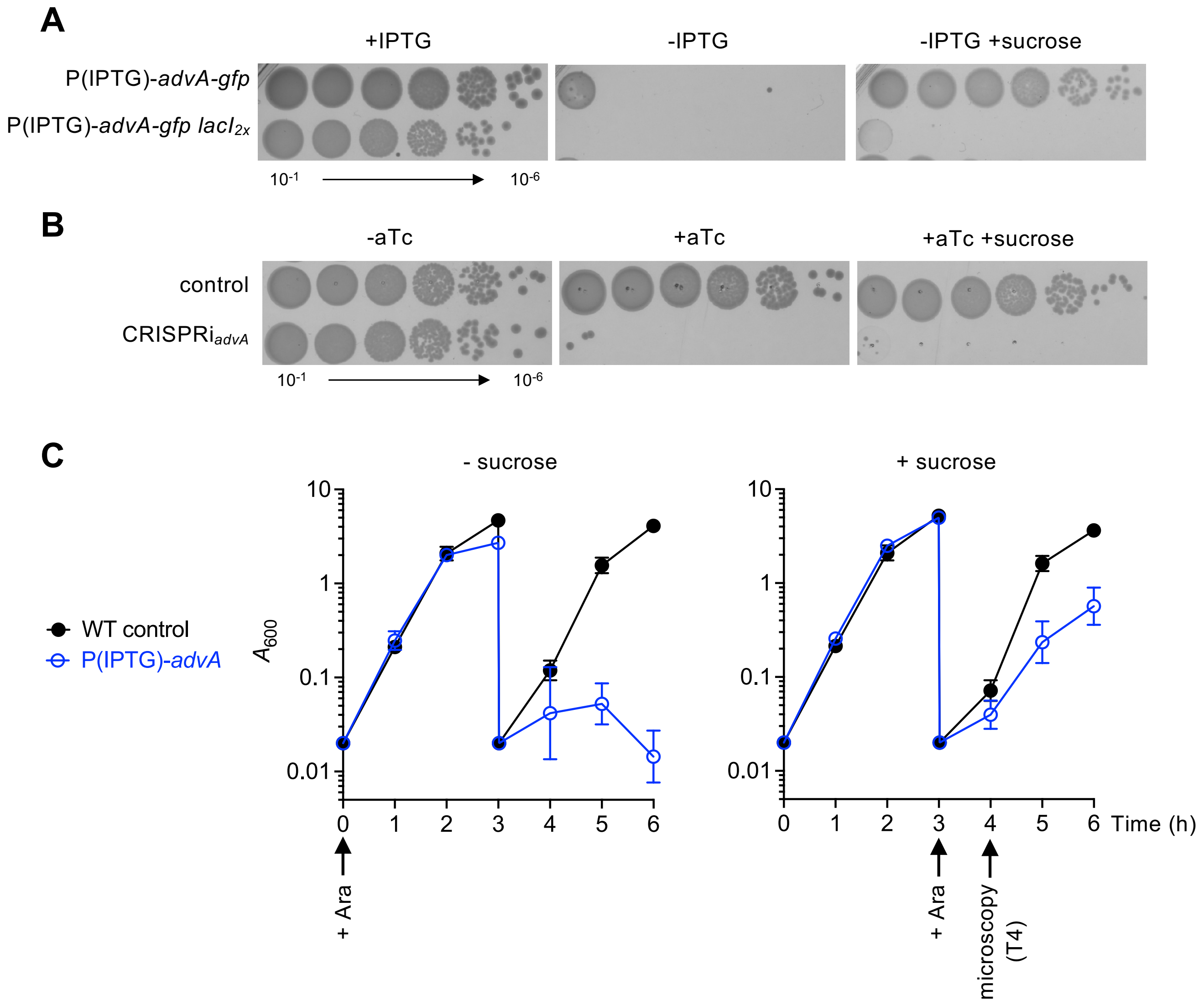


**Fig S5. Sucrose permits growth of bacteria with the conditional P(IPTG)-*advA* allele.** (A) Colony growth of *A. baumannii* conditional *advA* strains [P(IPTG)-*advA-gfp*] and [P(IPTG)-*advA-gfp* $lacI_{2x}$] on LBA with or without 500 µM IPTG or 10% sucrose. Strains were precultured in LB with 100 µM IPTG for 3 h and washed before spotting dilutions. (B) Colony growth of CRISPRi strains harboring control non-targeting sgRNA or *advA*-targeting sgRNA on LBA with or without CRISPRi inducer (50 ng/ml aTc) or 10% sucrose (B). Images representative of n=3 independent cultures (A, B). (C) Time course of WT vs AdvA-depleted bacterial growth in the presence and absence of sucrose. WT and P(IPTG)-*advA* strains with insertless P(Ara) vector (pYDE334) were precultured in LB (WT) or LB with 100 µM IPTG [P(IPTG)-*advA*] for 3 h. At T0, cells were washed twice with PBS and diluted to OD 0.02 in LB with 0.5% Ara (left) or 5% sucrose osmoprotectant (right). Medium lacked IPTG to deplete native AdvA. After 3 h culture, bacteria were diluted again to OD 0.02 in LB with 0.5% Ara (-sucrose, left; +sucrose, right), followed by continued culture for 3 h. Arrows indicate time of addition of Ara inducer and time point used for microscopy in Fig. 3. Data points show geometric mean ± s.d. (n=3).

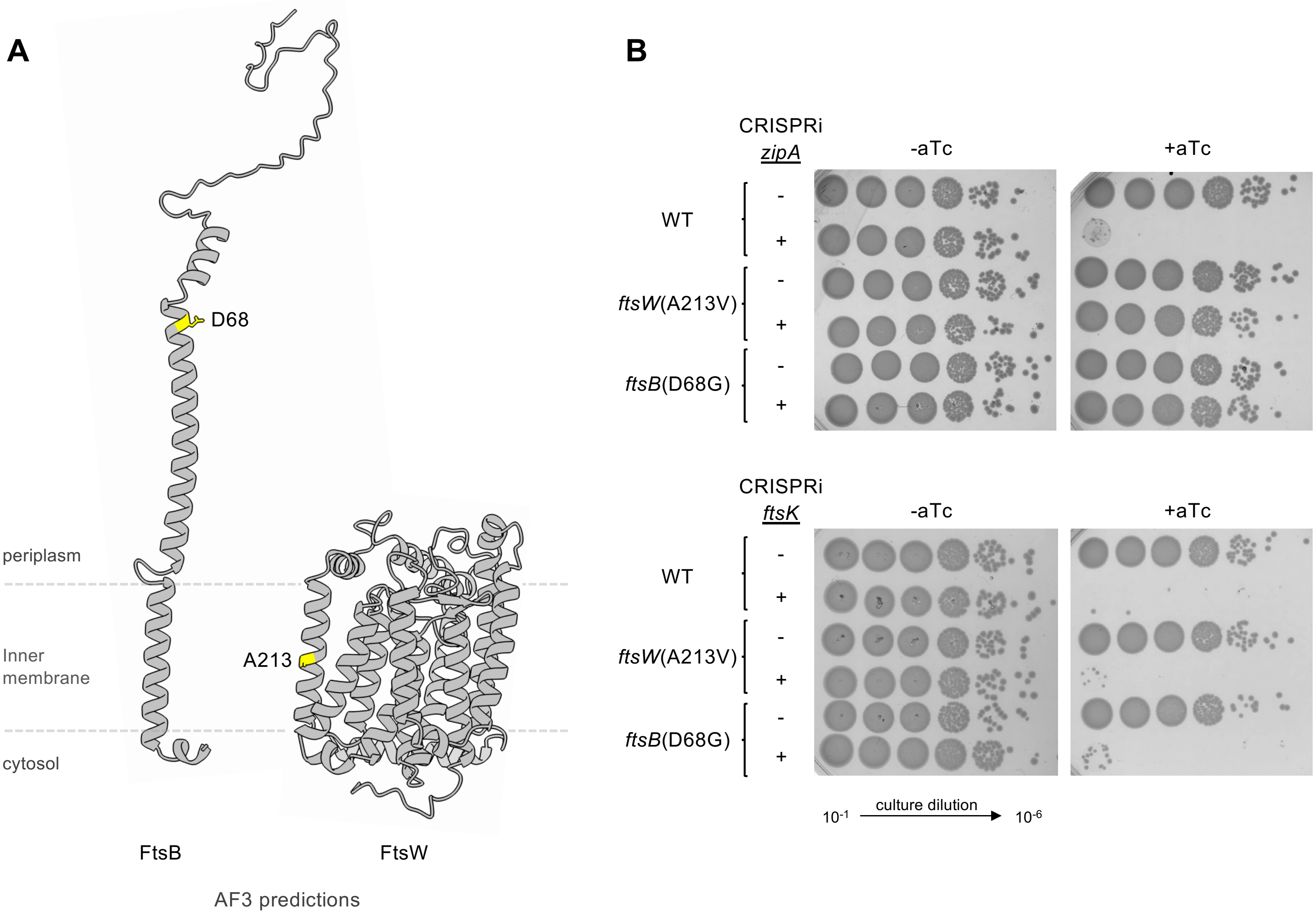


**Fig S6. Location and additional suppressor functions of amino acid substitutions in *A. baumannii* FtsB and FtsW that rescue AdvA$^{-}$ lethality.** (A) Ribbon diagrams show AlphaFold3 (AF3) predicted structures for FtsB (pTM = 0.42) and FtsW (pTM 0.92). Predicted topology relative to the inner membrane is indicated. Positions and amino acid side chains affected in suppressor mutants are shown in yellow. (B) Analysis of additional suppressor interactions using CRISPRi. Strains with the indicated genotype (left) harbored non-targeting control sgRNA (-) or an sgRNA targeting *zipA* or *ftsK* (+). Serial dilutions were spotted on solid medium with or without CRISPRi inducer (aTc, 50ng/ml). Images representative of n=3 independent cultures.

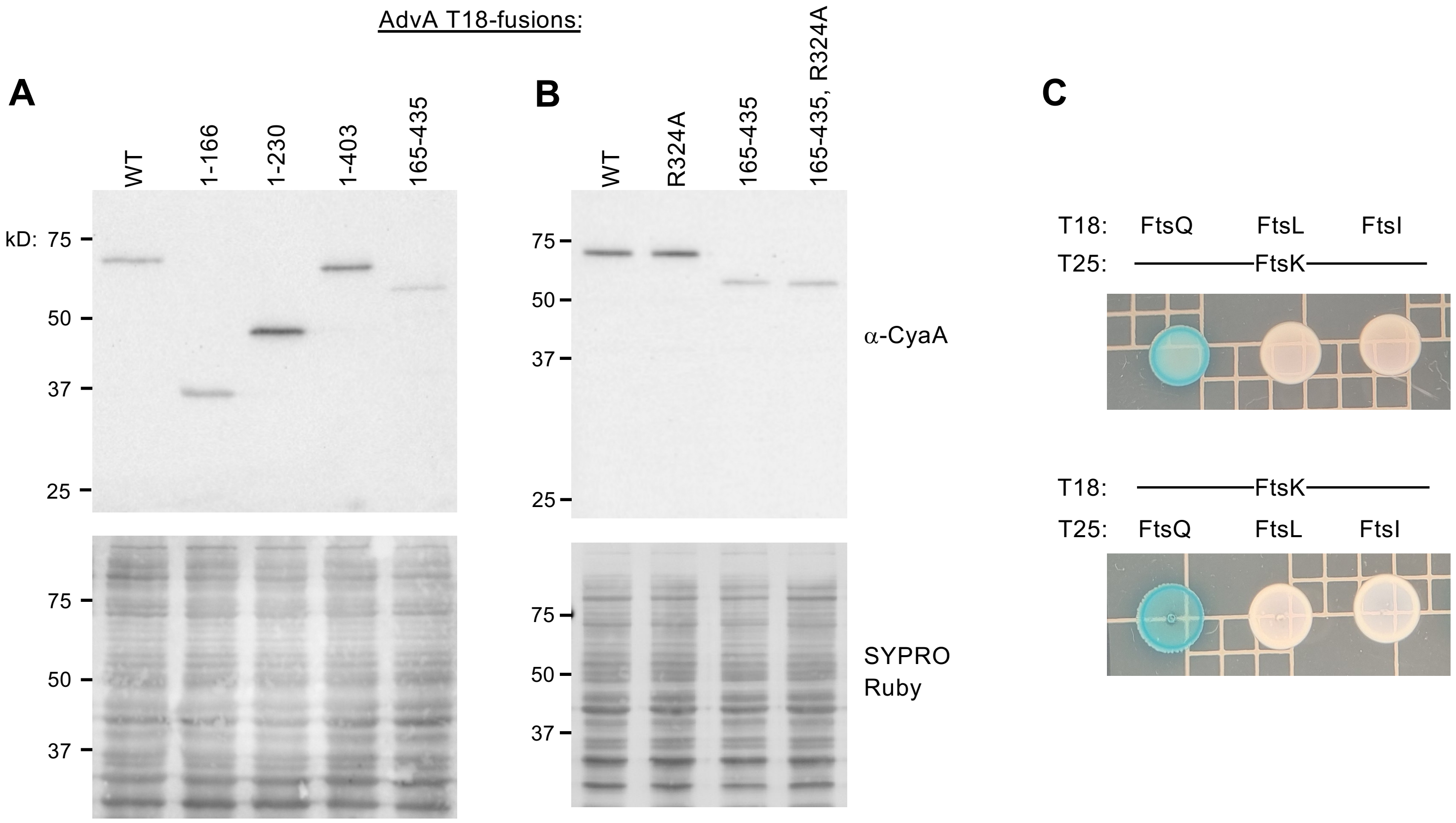


**Fig S7. Analysis of controls for two-hybrid assays.** (A-B) Western blot analysis in *E. coli* of T18 fusions to AdvA variants, including truncations with full-length control (A), and R324A point mutants and controls (B). BACTH strains harboring one of these fusions and an insertless pKNT25 plasmid were cultured in LB with Km (30 μg/ml), Cb (50 μg/ml), and IPTG (500 μM) at 30°C. Samples were collected at OD 1 and analyzed by SDS-PAGE and Western blot using anti-CyaA primary antibody (top) and SYPRO Ruby for total protein (bottom). (C) BACTH analysis in *E. coli* of interactions between *A. baumannii* FtsK and candidate *A. baumannii* divisome partners. Two-hybrid reporter analysis was performed as in Fig. 5.

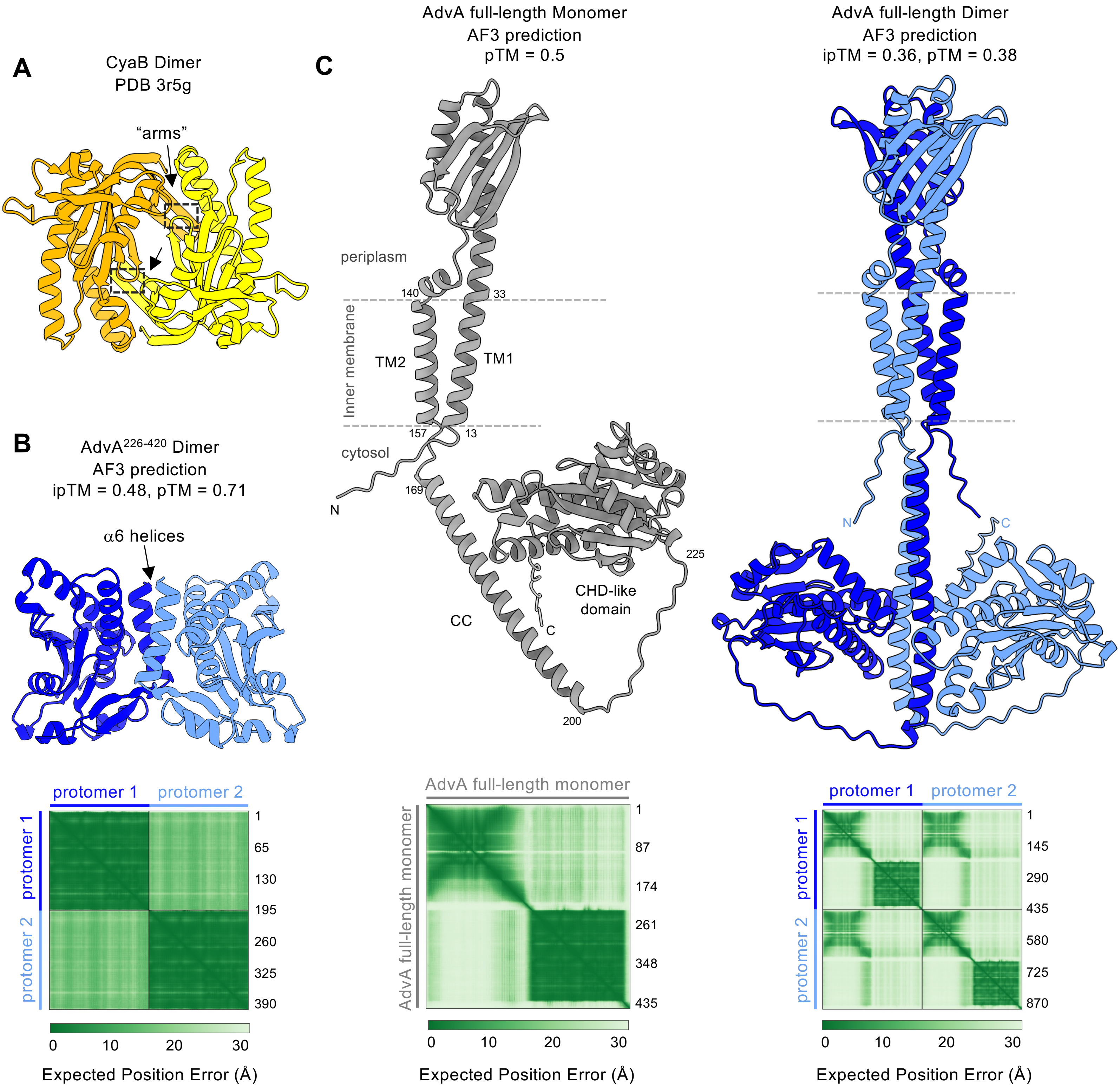


**Fig S8. Structural modeling analysis of predicted AdvA dimers.** (A) The CyaB dimer crystal structure (PDB: 3r5g, chains A,B) features a wreath-like organization dependent on the dimerization arms (arrows) and containing a central cleft that provides the active sites (dotted rectangles). (B, C) AlphaFold3 (AF3) structural models of theoretical homodimers of the AdvA CHD-like domain (B) and full-length protein (C, right), with individual protomers given in dark and light blue. Full-length protein monomer model is given in gray (C, left), with regions corresponding to the AdvA domains in Fig 6A labeled. Predicted membrane topology is based on the DeepTMHMM model. With each AF3 structural prediction, global ipTM and/or pTM confidence values are listed and the corresponding interface predicted aligned error (PAE) plot is shown below each model. Aligned residue number is given at the right of each plot.

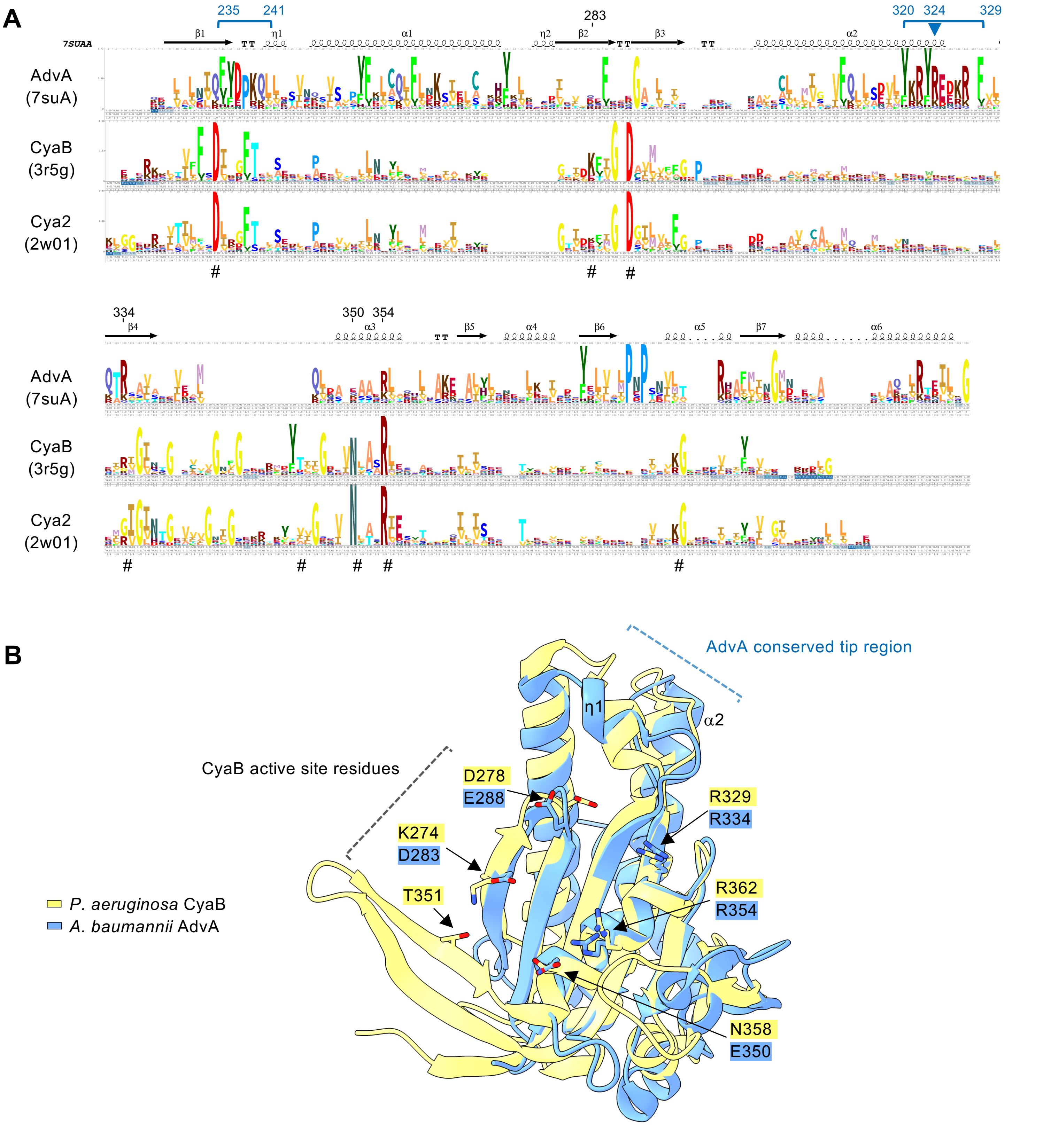


**Fig S9. Atypical topographical features of the AdvA CHD-like domain: lack of canonical AC/GC active site residues and presence of conserved tip.** (A) Stacked sequence logos show conserved residues at structurally aligned positions between AdvA$^{226-420}$ (PDB: 7sua), CyaB AC domain (PDB: 3r5g), and the GC Cya2 from *Synechocystis PCC6803* (PDB: 2w01), generated using DALI. Height of a stack of letters indicates sequence conservation at that position within each protein family; letter height reflects amino acid frequency at the position. Above logos is the secondary structure and residue numbers of AdvA$^{226-420}$. Blue brackets, conserved residues that contribute to an exposed tip at one end of AdvA; arrowhead, residue R324 in AdvA; #, active site residues in AC/GCs [1,2]. (B) Structural alignment of AdvA$^{226-420}$ with CyaB showing side chains and labels for 6 of the conserved catalytic residues in CyaB (yellow) and corresponding AdvA residues (blue).

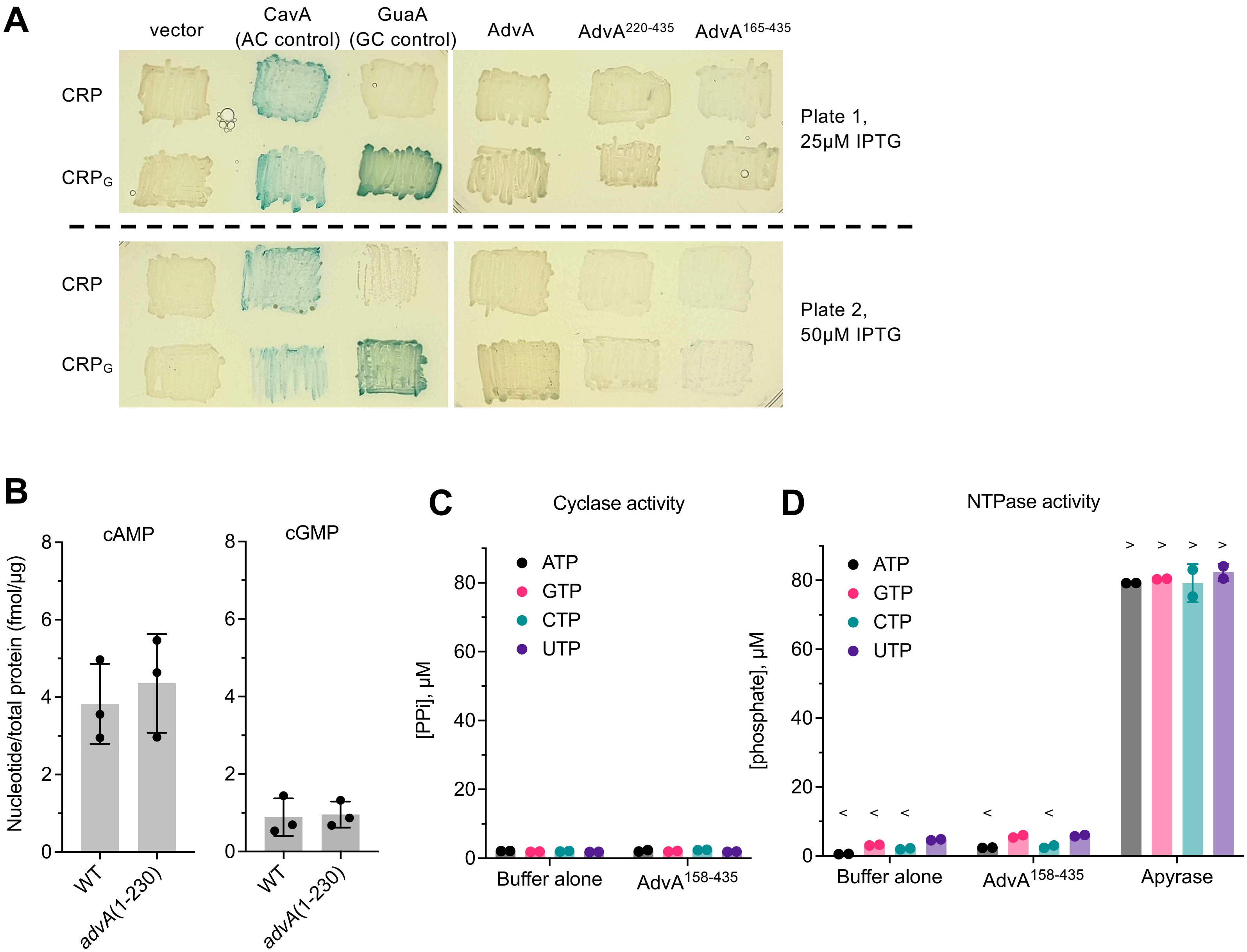


**Fig S10. AdvA CHD-like domain shows no detectable cyclase or NTPase activity under conditions tested *in vivo* and *in vitro*.** (A) *E. coli* CRP/CRP$_G$ reporter system was used to detect AC/GC activities of AdvA constructs. *E. coli* BL21[DE3] *cya crp* strains expressed WT CRP (sensing only cAMP) or a mutant CRP, CRP$_G$ (sensing both cAMP and cGMP) from a plasmid, and expressed a heterologous NC from a second plasmid. Expression of both components is controlled by IPTG. Blue color develops from CRP or CRP$_G$-dependent LacZ after growth on reporter plates with the indicated concentration of IPTG. Constructs encoding CavA from *A. baumannii* and GuaA from *Azospirillum* sp. B510 served as positive controls for AC and GC activity, respectively. The two adjacent images above or below the dotted line were from the same plate. Images representative of n=2 independent cultures. (B) ELISAs were used to examine effect of loss of AdvA CHD-like domain on *A. baumannii* cellular cAMP and cGMP levels. *A. baumannii* WT control and a mutant lacking the domain [*advA*(1-230) at native locus] were cultured in LB to OD 0.5, and cAMP and cGMP in cell lysates were measured by ELISA. Values were normalized to total protein level determined from BCA assay. Bars show means ± s.d. (n=3). (C-D) Evaluation of AdvA CHD-like domain NC and NTPase activity *in vitro*. Purified recombinant AdvA$^{158\text{-}435}$ containing N-terminal His-tag and cleavage site, or buffer alone, was used in NC assay (C) or NTPase assay (D) with the indicated nucleotide substrate. Apyrase was also used as a positive control for NTPase activity. Data were interpolated from phosphate standard curve, with < indicating values were below the lowest standard value, and > above the highest standard value. Bars show mean ± s.d. (n=2).

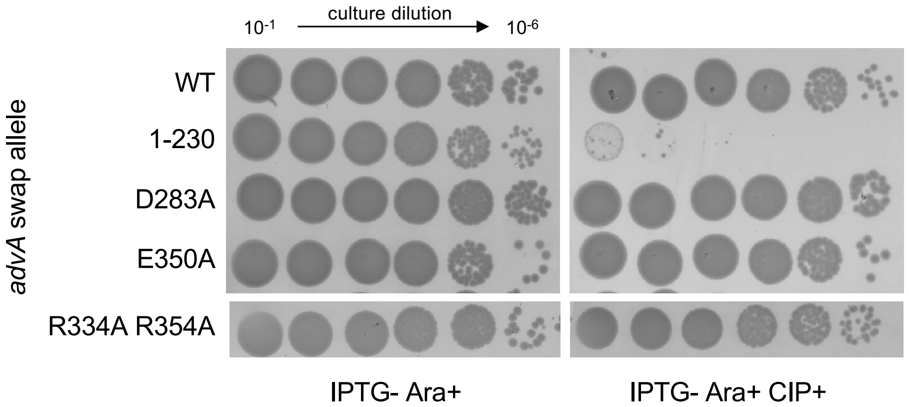


**Fig S11. AdvA function is preserved despite substitutions to positions aligning with the canonical AC/GC active site.** AdvA swap constructs were analyzed for effect on *A. baumannii* colony growth in the presence of fluoroquinolone antibiotic. Swap genes were expressed as fusions to msGFP2 via Ara induction as in Fig 1. Dilutions were spotted on LBA with 0.5% Ara and with or without CIP at 0.0625 µg/ml, and analyzed for colony formation. Images representative of n=3 independent cultures.

**Table S1. Strains used in this study**

| Strain ID | Designation | Genotype or description | Reference /source |
|---|---|---|---|
| ***A. baumannii strains (all ATCC 17978 background)*** | | | |
| AFA2 | WT ATCC 17978 | cerebrospinal fluid isolate, AbaAL44+ ("UN") type | [3], ATCC |
| AFA108 | P(IPTG)-*advA* | *advA*::*lacIq*-$P_{T5lac}$-*advA* | This work |
| AFA116 | P(IPTG)-*advA* insertless vector | AFA108 with pYDE334 | This work |
| AFA117 | P(IPTG)-*advA* P(Ara)-*advA*(1-166)-*msGFP2* | AFA108 with pAFE261 | This work |
| AFA118 | P(IPTG)-*advA* P(Ara)-*advA*(1-230)-*msGFP2* | AFA108 with pAFE262 | This work |
| AFA119 | P(IPTG)-*advA* P(Ara)-*advA*(220-435)-*msGFP2* | AFA108 with pAFE263 | This work |
| AFA120 | P(IPTG)-*advA* P(Ara)-*advA*(165-435)-*msGFP2* | AFA108 with pAFE264 | This work |
| AFA121 | P(IPTG)-*advA* P(Ara)-*advA*(WT)-*msGFP2* | AFA108 with pAFE267 | This work |
| AFA137 | P(IPTG)-*advA* P(Ara)-*advA*(1-287)-*msGFP2* | AFA108 with pAFE327 | This work |
| AFA138 | P(IPTG)-*advA* P(Ara)-*advA*(1-339)-*msGFP2* | AFA108 with pAFE328 | This work |
| AFA139 | P(IPTG)-*advA* P(Ara)-*advA*(1-391)-*msGFP2* | AFA108 with pAFE329 | This work |
| AFA286 | P(IPTG)-*advA* P(Ara)-*advA*(∆37-134)-*msGFP2* | AFA108 with pAFE547 | This work |
| AFA291 | P(IPTG)-*advA* P(Ara)-*advA*(1-403)-*msGFP2* | AFA108 with pAFE563 | This work |
| AFA319 | P(IPTG)-*advA* P(Ara)-*advA*(WT) untagged | AFA108 with pAFE597 | This work |
| AFA320 | P(IPTG)-*advA* P(Ara)-*advA*(1-287) untagged | AFA108 with pAFE598 | This work |
| AFA321 | P(IPTG)-*advA* P(Ara)-*advA*(1-391) untagged | AFA108 with pAFE599 | This work |
| AFA304 | P(IPTG)-*advA* P(Ara)-*advA*(1-403) untagged | AFA108 with pAFE581 | This work |
| AFA197 | P(Ara)-*advA*(WT)-*msGFP2* | AFA2 with pAFE267 | This work |
| AFA195 | P(Ara)-*advA*(220-435)-*msGFP2* | AFA2 with pAFE263 | This work |
| AFA196 | P(Ara)-*advA*(165-435)-*msGFP2* | AFA2 with pAFE264 | This work |
| AFA284 | P(Ara)-*advA*(∆37-134)-*msGFP2* | AFA2 with pAFE547 | This work |
| AFA316 | P(Ara)-*advA*(WT) untagged | AFA2 with pAFE597 | This work |
| AFA317 | P(Ara)-*advA*(1-287) untagged | AFA2 with pAFE598 | This work |
| AFA318 | P(Ara)-*advA*(1-391) untagged | AFA2 with pAFE599 | This work |
| AFA322 | P(Ara)-*advA*(1-403) untagged | AFA2 with pAFE581 | This work |
| AFA131 | *zapA-mCherry* | *zapA-mCherry* | This work |
| AFA132 | *zapA-mCherry* P(IPTG)-*advA* | *zapA-mCherry lacIq*-$P_{T5lac}$-*advA* | This work |
| AFA298 | *zapA-mCherry* insertless vector | AFA131 with pYDE334 | This work |
| AFA299 | *zapA-mCherry* P(IPTG)-*advA* insertless vector | AFA132 with pYDE334 | This work |
| AFA134 | *zapA-mCherry zipA-msGFP2* | AFA131 with pAFE317 | This work |
| AFA136 | *zapA-mCherry* P(IPTG)-*advA lacIq zipA-msGFP2* | AFA132 with pAFE317 | This work |
| AFA141 | *zapA-mCherry msGFP2-ftsW* | AFA131 with pAFE326 | This work |
| AFA143 | *zapA-mCherry* P(IPTG)-*advA msGFP2-ftsW* | AFA132 with pAFE326 | This work |
| AFA144 | *zapA-mCherry msGFP2-ftsN* | AFA131 with pAFE333 | This work |
| AFA145 | *zapA-mCherry* P(IPTG)-*advA msGFP2-ftsN* | AFA132 with pAFE333 | This work |
| AFA153 | *zapA-mCherry ftsK-msGFP2* | AFA131 with pAFE339 | This work |
| AFA155 | *zapA-mCherry* P(IPTG)-*advA ftsK-msGFP2* | AFA132 with pAFE339 | This work |
| AFA301 | *zapA-mCherry msGFP2-ftsB* | AFA131 with pAFE577 | This work |
| AFA302 | *zapA-mCherry* P(IPTG)-*advA msGFP2-ftsB* | AFA132 with pAFE577 | This work |
| YDA04 | WT CRISPRi parent | *attTn7::tetR-tetP-dcas9-rrnBT1-T7Te* $Gm^r$ | [4] |
| YDA07/08 | WT CRISPRi-control | YDA04 with pYDE007 | [4] |
| YDA011/012 | WT CRISPRi-*advA* | YDA04 with pYDE317-B | [4] |
| AFA28 | P(IPTG)-*advA-GFP* | *lacIq*-$P_{T5lac}$-*advA-GFPmut3* | This work |
| AFA29 | P(IPTG)-*advA-GFP* $lacI_{2x}$ | AFA28 with pEGE305 | This work |
| AFA45-16 | P(IPTG)-*advA-GFP* $lacI_{2x}$ *ftsW*(A213V) | AFA29 with spontaneous mutation conferring IPTG-independence *ftsW*(A213V) | This work |
| AFA45-18 | P(IPTG)-*advA-GFP* $lacI_{2x}$ *ftsB*(D68G) | AFA29 with spontaneous mutation conferring IPTG-independence *ftsB*(D68G) | This work |
| AFA59 | *ftsW*(A213V) CRISPRi parent | *attTn7::tetR-tetP-dcas9-rrnBT1-T7Te ftsW*(A213V) | This work |
| AFA60 | *ftsB*(D68G) CRISPRi parent | *attTn7::tetR-tetP-dcas9-rrnBT1-T7Te ftsB*(D68G) | This work |
| AFA62 | *ftsW*(A213V) CRISPRi-control | AFA59 with pYDE007 | This work |

| | | | |
|---|---|---|---|
| AFA63 | *ftsW*(A213V) CRISPRi-*advA* | AFA59 with pYDE317-B | This work |
| AFA65 | *ftsB*(D68G) CRISPRi-control | AFA60 with pYDE007 | This work |
| AFA66 | *ftsB*(D68G) CRISPRi-*advA* | AFA60 with pYDE317-B | This work |
| AFA33 | WT CRISPRi-*zipA* | YDA04 with pAFE112 | This work |
| AFA75 | *ftsW*(A213V) CRISPRi-*zipA* | AFA59 with pAFE112 | This work |
| AFA76 | *ftsB*(D68G) CRISPRi-*zipA* | AFA60 with pAFE112 | This work |
| AFA31 | WT CRISPRi-*ftsK* | YDA04 with pAFE117 | This work |
| AFA77 | *ftsW*(A213V) CRISPRi-*ftsK* | AFA59 with pAFE117 | This work |
| AFA78 | *ftsB*(D68G) CRISPRi-*ftsK* | AFA60 with pAFE117 | This work |
| AFA257 | P(IPTG)-*advA* P(Ara)-*advA*(D283A)-*msGFP2* | AFA108 with pAFE421 | This work |
| AFA260 | P(IPTG)-*advA* P(Ara)-*advA*(E350A)-*msGFP2* | AFA108 with pAFE427 | This work |
| AFA282 | P(IPTG)-*advA* P(Ara)-*advA*(R334A R354A)-*msGFP2* | AFA108 with pAFE531 | This work |
| AFA354 | P(IPTG)-*advA* P(Ara)-*advA*(K239A)-*msGFP2* | AFA108 with pAFE655 | This work |
| AFA341 | P(IPTG)-*advA* P(Ara)-*advA*(R324A)-*msGFP2* | AFA108 with pAFE645 | This work |
| AFA122 | *advA*(1-230) | *advA*(1-230) | This work |
| ***E. coli strains*** | | | |
| EGE1 | DH5α | *supE44 ΔlacU169* (*ϕ80lacZΔM15*) *hsdR17 recA1 endA1 gyrA96 thi-1 relA1* | [5] |
| EGE4 | DH5α λpir | DH5α (λpir) *tet*::Mu *recA* | [6] |
| AFE81 | XL1-Blue | *recA1 endA1 gyrA96 thi-1 hsdR17 supE44 relA1 lac* [*F´ proAB lacIq ZΔM15 Tn10 (Tc$^r$)*] | Agilent |
| NRE81 | BL21 (DE3) pLysS | F- *ompT hsdSB*(rB- mB-) *gal dcm* (DE3) pLysS (Cm$^r$) | Novagen |
| AFE6 | BTH101 | *F-, cya-99, araD139, galE15, galK16, rpsL1 (Str$^r$), hsdR2, mcrA1, mcrB1.* | [7] |
| AFE9 | BTH101 (-) insertless vectors | BTH101 with pKT25, pUT18C | [7] |
| AFE460 | BTH101 AdvA$^{WT}$-T18, T25-FtsA | BTH101 with pAFE458 and pAFE146 | This work |
| AFE461 | BTH101 AdvA$^{WT}$-T18, ZipA-T25 | BTH101 with pAFE458 and pAFE122 | This work |
| AFE462 | BTH101 AdvA$^{WT}$-T18, FtsK-T25 | BTH101 with pAFE458 and pAFE139 | This work |
| AFE463 | BTH101 AdvA$^{WT}$-T18, T25-FtsQ | BTH101 with pAFE458 and pAFE197 | This work |
| AFE464 | BTH101 AdvA$^{WT}$-T18, T25-FtsL | BTH101 with pAFE458 and pAFE198 | This work |
| AFE465 | BTH101 AdvA$^{WT}$-T18, T25-FtsB | BTH101 with pAFE458 and pAFE188 | This work |
| AFE466 | BTH101 AdvA$^{WT}$-T18, T25-FtsW | BTH101 with pAFE458 and pAFE190 | This work |
| AFE467 | BTH101 AdvA$^{WT}$-T18, T25-FtsI | BTH101 with pAFE458 and pAFE199 | This work |
| AFE468 | BTH101 AdvA$^{WT}$-T18, T25-FtsN | BTH101 with pAFE458 and pAFE200 | This work |
| AFE479 | BTH101 AdvA$^{1-166}$-T18, T25-FtsA | BTH101 with pAFE477 and pAFE146 | This work |
| AFE480 | BTH101 AdvA$^{1-166}$-T18, ZipA-T25 | BTH101 with pAFE477 and pAFE122 | This work |
| AFE481 | BTH101 AdvA$^{1-166}$-T18, FtsK-T25 | BTH101 with pAFE477 and pAFE139 | This work |
| AFE482 | BTH101 AdvA$^{1-166}$-T18, T25-FtsQ | BTH101 with pAFE477 and pAFE197 | This work |
| AFE483 | BTH101 AdvA$^{1-166}$-T18, T25-FtsL | BTH101 with pAFE477 and pAFE198 | This work |
| AFE484 | BTH101 AdvA$^{1-166}$-T18, T25-FtsB | BTH101 with pAFE477 and pAFE188 | This work |
| AFE485 | BTH101 AdvA$^{1-166}$-T18, T25-FtsW | BTH101 with pAFE477 and pAFE190 | This work |
| AFE486 | BTH101 AdvA$^{1-166}$-T18, T25-FtsI | BTH101 with pAFE477 and pAFE199 | This work |
| AFE487 | BTH101 AdvA$^{1-166}$-T18, T25-FtsN | BTH101 with pAFE477 and pAFE200 | This work |
| AFE503 | BTH101 AdvA$^{165-435}$-T18, T25-FtsA | BTH101 with pAFE299 and pAFE146 | This work |
| AFE504 | BTH101 AdvA$^{165-435}$-T18, ZipA-T25 | BTH101 with pAFE299 and pAFE122 | This work |
| AFE505 | BTH101 AdvA$^{165-435}$-T18, FtsK-T25 | BTH101 with pAFE299 and pAFE139 | This work |
| AFE506 | BTH101 AdvA$^{165-435}$-T18, T25-FtsQ | BTH101 with pAFE299 and pAFE197 | This work |
| AFE507 | BTH101 AdvA$^{165-435}$-T18, T25-FtsL | BTH101 with pAFE299 and pAFE198 | This work |
| AFE508 | BTH101 AdvA$^{165-435}$-T18, T25-FtsB | BTH101 with pAFE299 and pAFE188 | This work |
| AFE509 | BTH101 AdvA$^{165-435}$-T18, T25-FtsW | BTH101 with pAFE299 and pAFE190 | This work |
| AFE510 | BTH101 AdvA$^{165-435}$-T18, T25-FtsI | BTH101 with pAFE299 and pAFE199 | This work |
| AFE511 | BTH101 AdvA$^{165-435}$-T18, T25-FtsN | BTH101 with pAFE299 and pAFE200 | This work |
| AFE538 | BTH101 AdvA$^{1-230}$-T18, T25-FtsA | BTH101 with pAFE532 and pAFE146 | This work |
| AFE539 | BTH101 AdvA$^{1-230}$-T18, ZipA-T25 | BTH101 with pAFE532 and pAFE122 | This work |
| AFE540 | BTH101 AdvA$^{1-230}$-T18, FtsK-T25 | BTH101 with pAFE532 and pAFE139 | This work |
| AFE541 | BTH101 AdvA$^{1-230}$-T18, T25-FtsQ | BTH101 with pAFE532 and pAFE197 | This work |

| | | | |
|---|---|---|---|
| AFE542 | BTH101 AdvA$^{1-230}$-T18, T25-FtsL | BTH101 with pAFE532 and pAFE198 | This work |
| AFE543 | BTH101 AdvA$^{1-230}$-T18, T25-FtsB | BTH101 with pAFE532 and pAFE188 | This work |
| AFE544 | BTH101 AdvA$^{1-230}$-T18, T25-FtsW | BTH101 with pAFE532 and pAFE190 | This work |
| AFE545 | BTH101 AdvA$^{1-230}$-T18, T25-FtsI | BTH101 with pAFE532 and pAFE199 | This work |
| AFE546 | BTH101 AdvA$^{1-230}$-T18, T25-FtsN | BTH101 with pAFE532 and pAFE200 | This work |
| AFE586 | BTH101 AdvA$^{1-403}$-T18, T25-FtsA | BTH101 with pAFE583 and pAFE146 | This work |
| AFE587 | BTH101 AdvA$^{1-403}$-T18, ZipA-T25 | BTH101 with pAFE583 and pAFE122 | This work |
| AFE588 | BTH101 AdvA$^{1-403}$-T18, FtsK-T25 | BTH101 with pAFE583 and pAFE139 | This work |
| AFE589 | BTH101 AdvA$^{1-403}$-T18, T25-FtsQ | BTH101 with pAFE583 and pAFE197 | This work |
| AFE590 | BTH101 AdvA$^{1-403}$-T18, T25-FtsL | BTH101 with pAFE583 and pAFE198 | This work |
| AFE591 | BTH101 AdvA$^{1-403}$-T18, T25-FtsB | BTH101 with pAFE583 and pAFE188 | This work |
| AFE592 | BTH101 AdvA$^{1-403}$-T18, T25-FtsW | BTH101 with pAFE583 and pAFE190 | This work |
| AFE593 | BTH101 AdvA$^{1-403}$-T18, T25-FtsI | BTH101 with pAFE583 and pAFE199 | This work |
| AFE594 | BTH101 AdvA$^{1-403}$-T18, T25-FtsN | BTH101 with pAFE583 and pAFE200 | This work |
| AFE600 | BTH101 AdvA$^{1-403}$-T18, AdvA$^{WT}$-T25 | BTH101 with pAFE457 and pAFE583 | This work |
| AFE601 | BTH101 AdvA$^{WT}$-T18, AdvA$^{1-403}$-T25 | BTH101 with pAFE582 and pAFE458 | This work |
| AFE570 | BTH101 AdvA$^{WT}$-T18, AdvA$^{WT}$-T25 | BTH101 with pAFE457 and pAFE458 | This work |
| AFE573 | BTH101 AdvA$^{165-435}$-T18, AdvA$^{165-435}$-T25 | BTH101 with pAFE298 and pAFE299 | This work |
| AFE609 | BTH101 AdvA$^{1-166}$-T18, AdvA$^{1-166}$-T25 | BTH101 with pAFE476 and pAFE477 | This work |
| AFE610 | BTH101 AdvA$^{1-230}$-T18, AdvA$^{1-230}$-T25 | BTH101 with pAFE492 and pAFE532 | This work |
| AFE620 | BTH101 AdvA$^{1-403}$-T18, AdvA$^{1-403}$-T25 | BTH101 with pAFE582 and pAFE583 | This work |
| AFE569 | BTH101 AdvA$^{WT}$-T18, vector | BTH101 with pAFE458 and pKNT25 | This work |
| AFE568 | BTH101 AdvA$^{WT}$-T25, vector | BTH101 with pAFE457 and pUT18 | This work |
| AFE572 | BTH101 AdvA$^{165-435}$-T18, vector | BTH101 with pAFE299 and pKNT25 | This work |
| AFE571 | BTH101 AdvA$^{165-435}$-T25, vector | BTH101 with pAFE298 and pUT18 | This work |
| AFE607 | BTH101 AdvA$^{1-166}$-T18, vector | BTH101 with pAFE477 and pKNT25 | This work |
| AFE653 | BTH101 AdvA$^{1-166}$-T25, vector | BTH101 with pAFE476 and pUT18 | This work |
| AFE608 | BTH101 AdvA$^{1-230}$-T18, vector | BTH101 with pAFE532 and pKNT25 | This work |
| AFE654 | BTH101 AdvA$^{1-230}$-T25, vector | BTH101 with pAFE492 and pUT18 | This work |
| AFE606 | BTH101 AdvA$^{1-403}$-T18, vector | BTH101 with pAFE583 and pKNT25 | This work |
| AFE605 | BTH101 AdvA$^{1-403}$-T25, vector | BTH101 with pAFE582 and pUT18 | This work |
| AFE621 | BTH101 T18-FtsA, AdvA$^{WT}$-T25 | BTH101 with pAFE457 and pAFE147 | This work |
| AFE622 | BTH101 ZipA-T18, AdvA$^{WT}$-T25 | BTH101 with pAFE457 and pAFE142 | This work |
| AFE623 | BTH101 FtsK-T18, AdvA$^{WT}$-T25 | BTH101 with pAFE457 and pAFE140 | This work |
| AFE624 | BTH101 T18-FtsQ, AdvA$^{WT}$-T25 | BTH101 with pAFE457 and pAFE201 | This work |
| AFE625 | BTH101 T18-FtsL, AdvA$^{WT}$-T25 | BTH101 with pAFE457 and pAFE202 | This work |
| AFE626 | BTH101 T18-FtsB, AdvA$^{WT}$-T25 | BTH101 with pAFE457 and pAFE189 | This work |
| AFE627 | BTH101 T18-FtsW, AdvA$^{WT}$-T25 | BTH101 with pAFE457 and pAFE191 | This work |
| AFE628 | BTH101 T18-FtsI, AdvA$^{WT}$-T25 | BTH101 with pAFE457 and pAFE203 | This work |
| AFE629 | BTH101 T18-FtsN, AdvA$^{WT}$-T25 | BTH101 with pAFE457 and pAFE204 | This work |
| AFE630 | BTH101 T18-FtsA, AdvA$^{1-403}$-T25 | BTH101 with pAFE582 and pAFE147 | This work |
| AFE631 | BTH101 ZipA-T18, AdvA$^{1-403}$-T25 | BTH101 with pAFE582 and pAFE142 | This work |
| AFE632 | BTH101 FtsK-T18, AdvA$^{1-403}$-T25 | BTH101 with pAFE582 and pAFE140 | This work |
| AFE633 | BTH101 T18-FtsQ, AdvA$^{1-403}$-T25 | BTH101 with pAFE582 and pAFE201 | This work |
| AFE634 | BTH101 T18-FtsL, AdvA$^{1-403}$-T25 | BTH101 with pAFE582 and pAFE202 | This work |
| AFE635 | BTH101 T18-FtsB, AdvA$^{1-403}$-T25 | BTH101 with pAFE582 and pAFE189 | This work |
| AFE636 | BTH101 T18-FtsW, AdvA$^{1-403}$-T25 | BTH101 with pAFE582 and pAFE191 | This work |
| AFE637 | BTH101 T18-FtsI, AdvA$^{1-403}$-T25 | BTH101 with pAFE582 and pAFE203 | This work |
| AFE638 | BTH101 T18-FtsN, AdvA$^{1-403}$-T25 | BTH101 with pAFE582 and pAFE204 | This work |
| CKE37 | BTH101 FtsK-T25, T18-FtsQ | BTH101 with pAFE139 and pAFE201 | This work |
| CKE38 | BTH101 FtsK-T25, T18-FtsL | BTH101 with pAFE139 and pAFE202 | This work |
| CKE39 | BTH101 FtsK-T25, T18-FtsI | BTH101 with pAFE139 and pAFE203 | This work |
| CKE44 | BTH101 FtsK-T18, T25-FtsQ | BTH101 with pAFE140 and pAFE197 | This work |
| CKE45 | BTH101 FtsK-T18, T25-FtsL | BTH101 with pAFE140 and pAFE198 | This work |
| CKE46 | BTH101 FtsK-T18, T25-FtsI | BTH101 with pAFE140 and pAFE199 | This work |
| MSE159 | BTH101 AdvA$^{R324A}$-T18, ZipA-T25 | BTH101 with pMSE147 and pAFE122 | This work |
| MSE178 | BTH101 AdvA$^{165-435,R324A}$-T18, ZipA-T25 | BTH101 with pMSE170 and pAFE122 | This work |
| MSE197 | BTH101 ZipA-T18, AdvA$^{R324A}$-T25 | BTH101 with pAFE142 and pMSE150 | This work |

| | | | |
|---|---|---|---|
| MSE198 | BTH101 ZipA-T18, $AdvA^{165-435,\ R324A}$-T25 | BTH101 with pAFE142 and pMSE171 | This work |
| MSE221 | BTH101 AdvA-T18, RodA-T25 | BTH101 with pAFE458 and pYDE195 | This work |
| MSE222 | BTH101 AdvA-T18, T25-MreC | BTH101 with pAFE458 and pYDE188 | This work |
| MSE223 | BTH101 AdvA-T18, ElsS-T25 | BTH101 with pAFE458 and pYDE185 | This work |
| MSE224 | BTH101 $AdvA^{1-166}$-T18, T25-RodA | BTH101 with pAFE477 and pYDE195 | This work |
| MSE225 | BTH101 $AdvA^{1-166}$-T18, T25-MreC | BTH101 with pAFE477 and pYDE188 | This work |
| MSE226 | BTH101 $AdvA^{1-166}$-T18, ElsS-T25 | BTH101 with pAFE477 and pYDE185 | This work |
| MSE227 | BTH101 AdvA-T18, T25-PBP2 | BTH101 with pAFE458 and pYDE164 | This work |
| MSE228 | BTH101 AdvA-T18, MreD-T25 | BTH101 with pAFE458 and pYDE182 | This work |
| MSE229 | BTH101 $AdvA^{1-166}$-T18, T25-PBP2 | BTH101 with pAFE477and pYDE164 | This work |
| MSE230 | BTH101 $AdvA^{1-166}$-T18, MreD-T25 | BTH101 with pAFE477 and pYDE182 | This work |
| AFE441 | BL21[DE3] *cya crp* | *fhuA2* [*lon*] *ompT gal* (λ *DE3*) [*dcm*] Δ*hsdS crp*::*Kmr cyaA*::$Sp^r$ | [8] |
| AFE442 | BL21[DE3] *cya crp* CRP | BL21[DE3] *cya crp* with pRK-CRP | [8] |
| AFE443 | BL21[DE3] *cya crp* $CRP_G$ | BL21[DE3] *cya crp* with pRK-$CRP_G$ | [8] |
| AFE559 | BL21[DE3] *cya crp* CRP vector | AFE442 with pET23a(+) | This work |
| AFE560 | BL21[DE3] *cya crp* $CRP_G$ vector | AFE443 with pET23a(+) | This work |
| AFE561 | BL21[DE3] *cya crp* CRP CavA | AFE442 with pAFE566 | This work |
| AFE562 | BL21[DE3] *cya crp* $CRP_G$ CavA | AFE443 with pAFE566 | This work |
| AFE489 | BL21[DE3] *cya crp* CRP pETguaA | AFE442 with pAFE445 | This work |
| AFE490 | BL21[DE3] *cya crp* $CRP_G$ pETguaA | AFE443 with pAFE445 | This work |
| AFE533 | BL21[DE3] *cya crp* CRP $AdvA^{220-435}$ | AFE442 with pGPE254 | This work |
| AFE534 | BL21[DE3] *cya crp* $CRP_G$ $AdvA^{220-435}$ | AFE443 with pGPE254 | This work |
| AFE535 | BL21[DE3] *cya crp* CRP $AdvA^{165-435}$ | AFE442 with pAFE528 | This work |
| AFE536 | BL21[DE3] *cya crp* $CRP_G$ $AdvA^{165-435}$ | AFE443 with pAFE528 | This work |
| AFE550 | BL21[DE3] *cya crp* CRP AdvA | AFE442 with pAFE548 | This work |
| AFE551 | BL21[DE3] *cya crp* $CRP_G$ AdvA | AFE443 with pAFE548 | This work |

### Table S2. Plasmids used in this study

| Plasmid | Description | Reference |
|---|---|---|
| pUC18 | *oriColE1 MCS* $Cb^r$ cloning vector | [9] |
| ***Allelic exchange*** | | |
| pSR47S | Conditionally replicating allele exchange plasmid (oriTRP4 oriR6K *sacB* $Km^r$) | [10] |
| pAFE25 | pUC18 with mCherry C-terminal fusion construct | This work |
| pAFE104 | pSR47S with *lacI*q-$P_{T5lac}$-*advA* [P(IPTG)-*advA*] allele exchange construct | This work |
| pAFE324 | pSR47S with *zapA-mCherry* allele exchange construct | This work |
| pAFE65 | pSR47S with *advA-GFPmut3* allele exchange construct | This work |
| pAFE42 | pUC18::*advA-GFPmut3* | This work |
| pAFE43 | pEGE305::*advA-GFPmut3* | This work |
| pAFE185 | pSR47S with *ftsW*(A213V) allele exchange construct | This work |
| pAFE186 | pSR47S with *ftsB*(D68G) allele exchange construct | This work |
| pAFE272 | pSR47S with *advA*(1-230) allele exchange construct | This work |
| ***Gene swap/localization*** | | |
| pAFE225 | pUC18 containing msGFP2 C-terminal fusion construct | [11] |
| pAFE309 | pUC18 containing msGFP2 N-terminal fusion construct | This work |
| pEGE305 | P(IPTG) shuttle vector (*ori-pBR322 ori-pWH1277 bla*::*lacIq*-$P_{T5lac}$ $Tc^r$) | [12] |
| pYDE153 | P(IPTG) shuttle vector (*ori-pBR322 ori-pWH1277 bla*::*lacI*q-$P_{T5lac}$-MCS, $Tc^r$) | [4] |
| pYDE334 | pYDE153 with *araC-*$P_{BAD}$ replacing *lacI*q-$P_{T5lac}$ | This work |
| pAFE229 | pUC18::*advA*(WT)-*msGFP2* | This work |
| pAFE226 | pEGE305::*advA*(WT)-*msGFP2* | This work |
| pAFE261 | pYDE334::*advA*(1-166)-*msGFP2* | This work |
| pAFE262 | pYDE334::*advA*(1-230)-*msGFP2* | This work |
| pAFE263 | pYDE334::*advA*(220-435)-*msGFP2* | This work |
| pAFE264 | pYDE334::*advA*(165-435)-*msGFP2* | This work |
| pAFE267 | pYDE334::*advA*(WT)-*msGFP2* | This work |
| pAFE327 | pYDE334::*advA*(1-287)-*msGFP2* | This work |
| pAFE328 | pYDE334::*advA*(1-339)-*msGFP2* | This work |
| pAFE329 | pYDE334::*advA*(1-391)-*msGFP2* | This work |
| pAFE563 | pYDE334::*advA*(1-403)-*msGFP2* | This work |
| pAFE547 | pYDE334::*advA*(Δ37-134)-*msGFP2* | This work |
| pAFE317 | pYDE334::*zipA-msGFP2* | This work |
| pAFE326 | pYDE334::*msGFP2-ftsW* | This work |
| pAFE333 | pYDE334::*msGFP2-ftsN* | This work |
| pAFE339 | pYDE334::*ftsK-msGFP2* | This work |
| pAFE577 | pYDE334::*msGFP2-ftsB* | This work |
| pAFE597 | pYDE334::*advA*(WT) | This work |
| pAFE598 | pYDE334::*advA*(1-287) | This work |
| pAFE599 | pYDE334::*advA*(1-391) | This work |
| pAFE581 | pYDE334::*advA*(1-403) | This work |
| pAFE421 | pYDE334::*advA*(D283A)-*msGFP2* | This work |
| pAFE427 | pYDE334::*advA*(E350A)-*msGFP2* | This work |
| pAFE531 | pYDE334::*advA*(R334A R354A)-*msGFP2* | This work |
| pAFE645 | pYDE334::*advA*(R324A)-*msGFP2* | This work |
| pAFE655 | pYDE334::*advA*(K239A)-*msGFP2* | This work |
| ***BACTH*** | | |
| pKNT25 | Cloning and expression vector, pSU40 derivative with T25 domain of CyaA, MCS at the 3' start of T25, $Km^r$ | [7] |
| pUT18 | Cloning and expression vector, pUC19 derivative with T18 domain of CyaA, MCS at the 3' start of T18, $Cb^r$ | [7] |
| pKT25 | Cloning and expression vector, pSU40 derivative with T25 domain of CyaA, multiclonal sequence site (MCS) at the 3' end of T25, $Km^r$ | [7] |
| pUT18C | Cloning and expression vector, pUC19 derivative with T18 domain of Cya, MCS at the 3' end of T18, $Cb^r$ | [7] |
| pAFE458 | pUT18::*advA*(WT) | This work |
| pAFE146 | pKT25::*ftsA* | This work |
| pAFE122 | pKNT25::*zipA* | This work |

| | | |
|---|---|---|
| pAFE139 | pKNT25::*ftsK* | This work |
| pAFE197 | pKT25::*ftsQ* | This work |
| pAFE198 | pKT25::*ftsL* | This work |
| pAFE188 | pKT25::*ftsB* | This work |
| pAFE190 | pKT25::*ftsW* | This work |
| pAFE199 | pKT25::*ftsI* | This work |
| pAFE200 | pKT25::*ftsN* | This work |
| pAFE477 | pUT18::*advA*(1-166) | This work |
| pAFE299 | pUT18::*advA*(165-435) | This work |
| pAFE532 | pUT18::*advA*(1-230) | This work |
| pAFE457 | pKNT25::*advA*(WT) | This work |
| pAFE298 | pKNT25::*advA*(165-435) | This work |
| pAFE583 | pUT18::*advA*(1-403) | This work |
| pAFE582 | pKNT25::*advA*(1-403) | This work |
| pAFE476 | pKNT25::*advA*(1-166) | This work |
| pAFE492 | pKNT25::*advA*(1-230) | This work |
| pAFE147 | pUT18C::*ftsA* | This work |
| pAFE142 | pUT18::*zipA* | This work |
| pAFE140 | pUT18::*ftsK* | This work |
| pAFE201 | pUT18C::*ftsQ* | This work |
| pAFE202 | pUT18C::*ftsL* | This work |
| pAFE189 | pUT18C::*ftsB* | This work |
| pAFE191 | pUT18C::*ftsW* | This work |
| pAFE203 | pUT18C::*ftsI* | This work |
| pAFE204 | pUT18C::*ftsN* | This work |
| pMSE147 | pUT18::*advA*(R324A) | This work |
| pMSE150 | pKNT25::*advA*(R324A) | This work |
| pMSE170 | pUT18::*advA*(165-435, R324A) | This work |
| pMSE171 | pKNT25::*advA*(165-435, R324A) | This work |
| pYDE195 | pKT25::*rodA*(ACX60_RS05960) | [13] |
| pYDE164 | pKT25::*pbp2*(ACX60_RS13275) | [13] |
| pYDE188 | pKT25::*mreC*(ACX60_RS03610) | [13] |
| pYDE182 | pKNT25::*mreD*(ACX60_RS03615) | [13] |
| pYDE185 | pKNT25::*elsS*(ACX60_RS02860) | [13] |
| ***CRISPRi*** | | |
| pYDE007 | sgRNA delivery plasmid non-targeting control guide (ori-pBR322 ori-pWH1277 $P_{J23119}$-sgRNA*mrfp*-terminator, $Cb^r$) | [4] |
| pYDE317-B | pYDE007 derivative with sgRNA-*advA* | [4] |
| pAFE112 | pYDE007 derivative with sgRNA-*zipA* | This work |
| pAFE117 | pYDE007 derivative with sgRNA-*ftsK* | This work |
| ***AC/GC reporter assay*** | | |
| pET23a(+) | Bacterial expression vector, $Cb^r$ | Sigma |
| pAFE445 | pETguaA [pET23(a)+ encoding *Azospirillum* sp. B510 GuaA] | [8] |
| pAFE548 | pET23(a)+::*advA* | This work |
| pAFE528 | pET23(a)+::*advA*(165-435) | This work |
| pGPE254 | pET23(a)+::*advA*(220-435) | This work |
| pAFE566 | pET23(a)+::*cavA* (*A. baumannii* ACX60_RS09815) | This work |
| ***AdvA Purification*** | | |
| pMCSG53 | Vector with N-terminal $His_6$ tag and TEV recognition sequence; $Amp^r$ | [14] |
| pMCSG53-*advA* | pMCSG53::*advA*(158-435) | This work |

### Table S3. Synthetic oligonucleotides and gene fragments used in this study

| Primer name | Description | Sequence 5'--3' | RE site(s) |
|---|---|---|---|
| ***Allelic exchange*** | | | |
| NotI-U3387_F | Fwd for *advA* 5’ homology arm [P(IPTG)-*advA*, *advA*(1-230)] | CGCCAGCGGCCGCAATCGCAAGCATCGGTAAATC | NotI |
| newSphI-U3387R | Rev for *advA* 5’ homology arm [P(IPTG)-*advA*] | ACTTCGCATGCAATTATATTGAAATTCGATTATGATTCTTACA | SphI |
| lac-sphF | Fwd for P(IPTG)-*advA* 3’ homology arm | GCTGTCAAACATGAGAATTGCTCCGACTA | |
| lac_3387-salI_R | Rev for P(IPTG)-*advA* 3’ homology arm | CGCCGTCGACTGCTGCACTACTCGGGCTGT | SalI |
| AdvA_103982_PstI_R | Rev for *advA*(1-230) 5’ homology arm | ATTGCTGCAGTTATGCCAAGGTATTTTGGTCTGTAGACTCTAAGC | PstI |
| Pst-D3387-F | Fwd for *advA* 3’ homology arm | CAGCACTGCAGAAATTTATTGTGCGCATTTAAACCTAGCCATCC | PstI |
| Sal-D3387-R | Rev for *advA* 3’ homology arm | AGGACGTCGACCATGTACACGTACCGTTGTAACACCTTCT | SalI |
| ZapA_UpH_F | Fwd for *zapA* 5’ homology arm | CATCGGATCCGTGAAAATTCTGCTCTCTC | BamHI |
| ZapA_trlR | Rev for *zapA* 5’ homology arm [*zapA-mcherry*] | AATTCTAGACACTATTTGTTCAACATCTTCC | XbaI |
| ZapA_DownH_F | Fwd for *zapA* 3’ homology arm | ATACTGCAGACAATTCGCATAAAGCTTCATCAAAC | PstI |
| ZapA_DownH_R | Rev for *zapA* 3’ homology arm | GATTGTCGACGTGCAACTGACTACTATAAAGGG | SalI |
| ***Gene swap/localization*** | | | |
| 3387-bamF | Fwd for *advA* | CAACTAGGATCCGAATCATAATCGAATTTCAATATAATTGCCTTGG | BamHI |
| 3387-periR | Rev for *advA*(1-166), C fusion | CTAATCTAGAGCTACGTGTTGGGCGAGCAATG | XbaI |
| new3387-cytoF | Fwd for *advA*(165-435) | CCGGAATTCAGATTTAAGAAGGAGATATACATATGAGTCGTAGCGAATACTTAGCCCGTA | EcoRI |
| AdvA_287trlR | Rev for *advA*(1-287), C fusion | TCTAGATGCATGAAACTCGTCTACGAC | XbaI |
| AdvA_339trlR | Rev for *advA*(1-339), C fusion | TCTAGAATTACACACGGCGCTGC | XbaI |
| AdvA_391trlR | Rev for *advA*(1-391), C fusion | TCTAGAGACATTACTTGGGTTAGGC | XbaI |
| advA_Dperi_F | Fwd for *advA*(Δ37-134) | ACGGTTACTAGTACCCTGGTGTGTCGTTGCAATCACC | SpeI |
| advA_Dperi_R | Rev for *advA*(Δ37-134) | CAACCAACTAGTGGGGAAATTCTTCGTACCCAATG | SpeI |
| advA_1-403_R | Rev for *advA*(1-403), C fusion | ACATCTAGAGTTCATACCATTGATCATAAATGCATGACG | XbaI |
| AdvA_220F | Fwd for *advA*(220-435) | GCTAGAATTCTTTAAGAAGGAGATATACATATGAGCTTAGAGTCTACAGACC | EcoRI |
| advA_pstR_2.0 | Rev for *advA* untagged | GCTACTGCAGAATGCGCACAATAAATTTTTAAGATGCTGCACTACTCGG | PstI |
| AdvA_287_R | Rev for *advA*(1-287) untagged | TGTGACTGCAGAATGCGCACAATAAATTTTTATGCATGAAACTCGTCTACGAC | PstI |
| AdvA_391_R | Rev for *advA*(1-391) untagged | TACCCTGCAGAATGCGCACAATAAATTTTTAGACATTACTTGGGTTAGGC | PstI |
| AdvA_403_nontrlR | Rev for *advA*(1-403) untagged | CGAATCTGCAGAATGCGCACAATAAATTTTTAATTCATACCATTGATCATAAATGC | PstI |
| msGFP2_NtrlF | Fwd for msGFP2 | GAGCTCAGATTTAAGAAGGAGATATACATATGGATTCTACTGAATCTTTATTCACTG | SacI |
| msGFP2_NtrlR | Rev for msGFP2, N fusion | ACTAGTGCCGCCGCCGCCGCCAGAACCACCAGAACCACCAG | SpeI |
| ZipA_trlF | Fwd for *zipA* | GGATCCGAACAGGCAAAAGAATACGGGTTAGTTT | BamHI |
| ZipA_trlR | Rev for *zipA*, C fusion | TCTAGAGGCTGTTGCCTGTGCTGGAC | XbaI |
| FtsK_BamF | Fwd for *ftsK* | AATTGGATCCGTTACAGGACAGTATATGACTGCGGTG | BamHI |
| FtsK_XbaR | Rev for *ftsK*, C fusion | GTAGTCTAGAAACTAAAATATCGCGCTTACCGTTG | XbaI |
| FtsW_tlrF | Fwd for *ftsW*, N fusion | ACTAGTATGGCAGGCTTAGCTCAGAC | SpeI |
| FtsW_tlrR | Rev for *ftsW* | CTGCAGCGCTTCATGGTTTAGAAGTTTG | PstI |
| FtsN_tlrF | Fwd for *ftsN*, N fusion | ACTAGTGTGTTTGGCAAAACGCAACGCG | SpeI |
| FtsN_tlrR | Rev for *ftsN* | CTGCAGCTTAATCTTTATTTGCGTTTAATCACAATAGAGTCAATAC | PstI |
| FtsB_trlF | Fwd for *ftsB*, N fusion | ACTAGTATGTTAGAAGTATTTCGTTCCACTTCAAGCAAAC | SpeI |
| FtsB_trlR | Rev for *ftsB* | CTGCAGCTGCCCATAATTTAATCTGGGATG | PstI |
| ***BACTH*** | | | |
| new_advA_sphF_2.0 | Fwd for *advA* [BACTH C fusion] | GCATGCGAATCATAATCGAATTTCAATATAATTGCCTTGG | SphI |
| 3387-Xba-R | Rev for *advA* [BACTH C fusion] | TAAATTCTAGAGATGCTGCACTACTCGGGCTGTC | XbaI |
| new_advA_peri_B2H_2.0_R | Rev for *advA*(1-166) [BACTH C fusion] | GGCTTCTAGACTGCTACGTGTTGGGCGAGCAATG | |
| advA_1_230_B2H_2.0_R | Rev for *advA*(1-230) [BACTH C fusion] | TTGATCTAGACATGCCAAGGTATTTTGGTCTGTAGACTCTAAGC | XbaI |
| AdvAcyto_SphIF | Fwd for *advA*(165-435) [BACTH C fusion] | GCATGCAGATTTAAGAAGGAGATATACATATGAGTCGTAGCGAATACTTAGCCCGTA | SphI |
| AdvA_403_B2H_R | Rev for *advA*(1-403) [BACTH C fusion] | ACATCTAGACTGTTCATACCATTGATCATAAATGCATGACG | XbaI |
| FtsAF-BACTH | Fwd for *ftsA* [BACTH N fusion] | TGGTGGATCCTATGAGTGAAGCTGTTCCCTCAGTTG | BamHI |
| FtsAR-BACTH | Rev for *ftsA* [BACTH N fusion] | ATAAGAATTCACATCCTAAAAAATGGCTTTAAGTTTGC | EcoRI |
| ZipA-Sph-F | Fwd for *zipA* [BACTH C fusion] | GGAGGCATGCTATGGAAATCAATACGATTATTGGCATCGTTGTTG | SphI |
| ZipA-Xba-R | Rev for *zipA* [BACTH C fusion] | AAATTCTAGAATGGCTGTTGCCTGTGCTGGACGATA | XbaI |
| FtsK-B2H_F | Fwd for *ftsK* [BACTH C fusion] | AGGAGCATGCCATGACTGCGGTGTCAAGTGTTTATGCACAG | SphI |
| FtsK-B2H_R | Rev for *ftsK* [BACTH C fusion] | GTAGTCTAGACGAACTAAAATATCGCGCTTACCGTTGG | XbaI |
| FtsQ-bamF | Fwd for *ftsQ)* [BACTH N fusion] | ACGGGGATCCTATGGCACAACTTCCGGCATC | BamHI |

| | | | |
|---|---|---|---|
| FtsQ-ecoR | Rev for *ftsQ* [BACTH N fusion] | AAGCGAATTCTTATGGCTTTGCTTTTGTACCACCTG | EcoRI |
| FtsL-bamF | Fwd for *ftsL* [BACTH N fusion] | AAGGGGATCCTATGAAAAGCAGTGATGAAATCG | BamHI |
| FtsL-ecoR | Rev for *ftsL* [BACTH N fusion] | AGCAGAATTCTTACTTATTTTGCTCTGAGGTC | EcoRI |
| FtsB-bamF | Fwd for *ftsB* [BACTH N fusion] | CTCTGGATCCTATGTTAGAAGTATTTCGTTCC | BamHI |
| FtsB-ecoR | Rev for *ftsB* [BACTH N fusion] | TGCCGAATTCTTAATCTGGGATGTCTGGTG | EcoRI |
| FtsW-bamF | Fwd for *ftsW* [BACTH N fusion] | AGGAGGATCCAATGGCAGGCTTAGCTCAGACC | BamHI |
| FtsW-ecoR | Rev for *ftsW* [BACTH N fusion] | GCTTGAATTCTTAGAAGTTTGATTCTTCCCTCTCAGG | EcoRI |
| FtsI-bamF | Fwd for *ftsI* [BACTH N fusion] | GGCCGGATCCTATGGTAGATAAGCGAACAAAGCAA | BamHI |
| FtsI-ecoR | Rev for *ftsI* [BACTH N fusion] | CAGTGAATTCTTACCTGCGAATAGGATTTTCTGGAGTATTAAG | EcoRI |
| FtsN-bamF | Fwd for *ftsN* [BACTH N fusion] | AATAGGATCCTGTGTTTGGCAAAACGCAACG | BamHI |
| FtsN-ecoR | Rev for *ftsN* [BACTH N fusion] | TACTGAATTCTTATTTGCGTTTAATCACAATAGAGTCAATAC | EcoRI |
| ***CRISPRi*** | | | |
| sgr-*zipA* | Fwd for sgRNA-*zipA* | TTACACTAGTGTCTAGCGAGGGTTCAGCATGGTTGTTTTAGAGCTAG<br>AAATAGCAAG | SpeI |
| sgr-*ftsK* | Fwd for sgRNA-*ftsK* | TTACACTAGTATCACTTGAAATATGCATCCAGCCGTTTTAGAGCTAG<br>AAATAGCAAG | SpeI |
| sgr-rev | Rev for sgRNAs | AAGTGGGCCCAGATCTAAGCTTCAAAAAAAGCACCGAC | ApaI |
| ***AC/GC reporter assay*** | | | |
| advA_GC_F | Fwd for *advA* [pET insert] | AATACATATGCCTTGGAAGTTTAACTTGATTGCG | NdeI |
| advA_GC_R | Rev for *advA* [pET insert] | GCTAGTCGACAATGCGCACAATAAATTTTTAAGATGCTGC | SalI |
| advA_cyto_GC_F | Fwd for *advA*(165-435) [pET insert] | TCGCCATATGCGTAGCGAATACTTAGCCCGTA | NdeI |
| advA_220_GC_F | Rev for *advA*(220-435) [pET insert] | ACCACATATGAGCTTAGAGTCTACAGACCAA | NdeI |
| gcS_GC_ampF | Fwd for *cavA* [pET insert] | AGGAGGCTAGCCAATTAGAGAGGTTTATAGATCG | NheI |
| gcS_GC_ampR | Rev for *cavA* [pET insert] | GCAGCAAGCTTTATTTATATAATTTATACCTGATCTTGTTCTTC | HindIII |
| ***Protein purification*** | | | |
| ACX60_00475-F | Fwd for *advA*(158-435) [pMCSG53 insert] | TACTTCCAATCCAATGCCCGTGCCATTGCTCGCCCAACA | |
| ACX60_00475-R | Ref for *advA*(158-435) [pMCSG53 insert] | TTATCCACTTCCAATGTTAAGATGCTGCACTACTCGGGCT | |
| **Synthetic gene fragment name** | **Sequence 5'--3'** | | **RE site(s)** |
| mCherry-codonopt-1 | GGTTCGTCTAGAGGTGGTGGTGGCATGGTTTCTAAAGGTGAAGAAGATAACATGGCTATTATTAAAG<br>AGTTTATGCGTTTCAAAGTTCACATGGAAGGTTCTGTTAACGGTCACGAGTTTGAAATTGAAGGTGAAGGT<br>GAAGGTCGTCCATATGAAGGTACTCAAACTGCTAAATTAAAAGTTACTAAAGGTGGTCCATTACCATTCGC<br>TTGGGATATTTTATCTCCACAATTCATGTATGGTTCTAAAGCTTATGTTAAACACCCAGCTGATATTCCAGA<br>CTATTTGAAATTGTCATTCCCAGAAGGTTTCAAATGGGAACGTGTTATGAACTTCGAAGATGGTGGTGTTG<br>TTACTGTTACTCAAGATTCTTCTTTACAAGATGGTGAGTTCATTTATAAGGTGAAGTTACGTGGTACTAACT<br>TCCCATCTGATGGTCCAGTTATGCAAAAGAAAACTATGGGTTGGGAAGCTTCTTCTGAACGTATGTATCCA<br>GAAGATGGTGCATTAAAAGGTGAAATTAAACAACGTTTAAAACTTAAAGATGGTGGTCATTATGATGCAGA<br>AGTGAAAACTACTTATAAAGCCAAGAAACCAGTGCAGTTGCCAGGTGCTTATAACGTTAACATTAAACTTG<br>ACATTACTTCTCACAACGAAGATTATACTATTGTTGAACAATATGAACGTGCTGAAGGTCGTCACTCTACT<br>GGTGGTATGGATGAATTGTATAAATAATGATCCAGACCTGCAGTTCTGG | | XbaI, PstI |
| AdvA R324A | CAACTAGGATCCGAATCATAATCGAATTTCAATATAATTGCCTTGGAAGTTTAACTTGATTGCGCCTAGAC<br>AAGGGCTATTTGCTAGCCTACTGATCATTAGTTTTGCATTGCATACCTTTTTATTGGTGATTGCAACGACA<br>CACCAGCTCAATGAAAACCGTGCGAGTCAAGGTCAGCTCATGACCAGTCAACTCGTTGCAGACAGCTTAT<br>CTGAGCTTGAACCTGCCAATACGGTCTCTTTAGCTCTGATTGCTAATCGCTATGCGACCAATCCAAGTGT<br>GGCATCTATCCGTATTCTTGACGCCAATAAACAAGTTTTGGCAACAAGCGGCATGTCAAAAACTCGCCAA<br>GGTGAAATTTTTGTTCGTGATGCGCTTCAAAACGAGAAAAAAGTCGGTTCTATTGAAATCACACTAATTCA<br>ACCAAGTATTGGGGAAATTCTTCGTACCCAATGGCTTGCGATTTTAGCCTCCCTGTTCTTACATGTTTTATT<br>ATGGCTAGCTTACCGTGCCATTGCTCGCCCAACACGTAGCGAATACTTAGCCCGTATTAACGAAGAAAAT<br>CGTTTAAAACATGAAATTCAAGAGTTAACTCAAGCTCTCGCGCTTGAAAAACAAAATACAGTGACCTTGGT<br>TGCTCAAGCTCAGCAGCAAGCTAAAGCTAAACCAATTGTTCGCTCACAACCAGAAAAAAGCTTAGAGTCT<br>ACAGACCAAAATACCTTGGCACTTAATATTCAATTTTATGACCCTAAACAGTTATTAAGTAGTGTTAACCAG<br>TCTGTTTCTGTACCGTACTTTAAACTTTGTCAGCTCTTCTTAAACAAGAGTATTGAGCTATGTACAAAGCAT<br>TATCATTTGAAAGCGACTGATATTGATGTCGTAGACGAGTTTCATGCAGAGGGCGCAACACTTGCCATTT<br>CGACTTCACATCCACACGCTGTAGAGTGTCTACTTATGGTAGGTACGGTTTTCCAATTGCTTTCAGACGTT<br>TTATATAAGCGTTACGCTGAAGATAAACGCTTTGCGCTACAAACTCGCAGCGCCGTGTGTAATGCCGTAG<br>AAGCAATGCAAATTGATGCCAAAGAAGCTGCTCAACGTTTAGCGCAACATCTTCATGCGAAAGAATCTGC<br>ACTCTACCTTGATAATGAACAGCTTAAAGCAATTCAAGACAGCTATCAGCTTGTTGCAATGCCTAACCCAA<br>GTAATGTCATGACTCGTCATGCATTTATGATCAATGGTATGAATGCTGAATGTGCAGAGCTTGCACAAAAT<br>ATTCGAACTGAAATTTTGATGGGTAAAAAATCAATTCCTCAAAATGACAGCCCGAGTAGTGCAGCATCTAG<br>AAAATT | | BamHI, XbaI |
| AdvA K239A | CAACTAGGATCCGAATCATAATCGAATTTCAATATAATTGCCTTGGAAGTTTAACTTGATTGCGCCTAGAC<br>AAGGGCTATTTGCTAGCCTACTGATCATTAGTTTTGCATTGCATACCTTTTTATTGGTGATTGCAACGACA<br>CACCAGCTCAATGAAAACCGTGCGAGTCAAGGTCAGCTCATGACCAGTCAACTCGTTGCAGACAGCTTAT<br>CTGAGCTTGAACCTGCCAATACGGTCTCTTTAGCTCTGATTGCTAATCGCTATGCGACCAATCCAAGTGT<br>GGCATCTATCCGTATTCTTGACGCCAATAAACAAGTTTTGGCAACAAGCGGCATGTCAAAAACTCGCCAA<br>GGTGAAATTTTTGTTCGTGATGCGCTTCAAAACGAGAAAAAAGTCGGTTCTATTGAAATCACACTAATTCA<br>ACCAAGTATTGGGGAAATTCTTCGTACCCAATGGCTTGCGATTTTAGCCTCCCTGTTCTTACATGTTTTATT<br>ATGGCTAGCTTACCGTGCCATTGCTCGCCCAACACGTAGCGAATACTTAGCCCGTATTAACGAAGAAAAT<br>CGTTTAAAACATGAAATTCAAGAGTTAACTCAAGCTCTCGCGCTTGAAAAACAAAATACAGTGACCTTGGT<br>TGCTCAAGCTCAGCAGCAAGCTAAAGCTAAACCAATTGTTCGCTCACAACCAGAAAAAAGCTTAGAGTCT<br>ACAGACCAAAATACCTTGGCACTTAATATTCAATTTTATGACCCTGCTCAGTTATTAAGTAGTGTTAACCAG<br>TCTGTTTCTGTACCGTACTTTAAACTTTGTCAGCTCTTCTTAAACAAGAGTATTGAGCTATGTACAAAGCAT<br>TATCATTTGAAAGCGACTGATATTGATGTCGTAGACGAGTTTCATGCAGAGGGCGCAACACTTGCCATTT | | BamHI, XbaI |

| | | |
|---|---|---|
| | CGACTTCACATCCACACGCTGTAGAGTGTCTACTTATGGTAGGTACGGTTTTCCAATTGCTTTCAGACGTT<br>TTATATAAGCGTTACCGTGAAGATAAACGCTTTGCGCTACAAACTCGCAGCGCCGTGTGTAATGCCGTAG<br>AAGCAATGCAAATTGATGCCAAAGAAGCTGCTCAACGTTTAGCGCAACATCTTCATGCGAAAGAATCTGC<br>ACTCTACCTTGATAATGAACAGCTTAAAGCAATTCAAGACAGCTATCAGCTTGTTGCAATGCCTAACCCAA<br>GTAATGTCATGACTCGTCATGCATTTATGATCAATGGTATGAATGCTGAATGTGCAGAGCTTGCACAAAAT<br>ATTCGAACTGAAATTTTGATGGGTAAAAAATCAATTCCTCAAAATGACAGCCCGAGTAGTGCAGCATCTAG<br>AAAATT | |
| AdvA D283A | CAACTAGGATCCGAATCATAATCGAATTTCAATATAATTGCCTTGGAAGTTTAACTTGATTGCGCCTAGAC<br>AAGGGCTATTTGCTAGCCTACTGATCATTAGTTTTGCATTGCATACCTTTTTATTGGTGATTGCAACGACA<br>CACCAGCTCAATGAAAACCGTGCGAGTCAAGGTCAGCTCATGACCAGTCAACTCGTTGCAGACAGCTTAT<br>CTGAGCTTGAACCTGCCAATACGGTCTCTTTAGCTCTGATTGCTAATCGCTATGCGACCAATCCAAGTGT<br>GGCATCTATCCGTATTCTTGACGCCAATAAACAAGTTTTGGCAACAAGCGGCATGTCAAAAACTCGCCAA<br>GGTGAAATTTTTGTTCGTGATGCGCTTCAAAACGAGAAAAAAGTCGGTTCTATTGAAATCACACTAATTCA<br>ACCAAGTATTGGGGAAATTCTTCGTACCCAATGGCTTGCGATTTTAGCCTCCCTGTTCTTACATGTTTTATT<br>ATGGCTAGCTTACCGTGCCATTGCTCGCCCAACACGTAGCGAATACTTAGCCCGTATTAACGAAGAAAAT<br>CGTTTAAAACATGAAATTCAAGAGTTAACTCAAGCTCTCGCGCTTGAAAAACAAAATACAGTGACCTTGGT<br>TGCTCAAGCTCAGCAGCAAGCTAAAGCTAAACCAATTGTTCGCTCACAACCAGAAAAAAGCTTAGAGTCT<br>ACAGACCAAAATACCTTGGCACTTAATATTCAATTTTATGACCCTAAACAGTTATTAAGTAGTGTTAACCAG<br>TCTGTTTCTGTACCGTACTTTAAACTTTGTCAGCTCTTCTTAAACAAGAGTATTGAGCTATGTACAAAGCAT<br>TATCATTTGAAAGCGACTGATATTGATGTCGTAGCTGAGTTTCATGCAGAGGGCGCAACACTTGCCATTTC<br>GACTTCACATCCACACGCTGTAGAGTGTCTACTTATGGTAGGTACGGTTTTCCAATTGCTTTCAGACGTTT<br>TATATAAGCGTTACCGTGAAGATAAACGCTTTGCGCTACAAACTCGCAGCGCCGTGTGTAATGCCGTAGA<br>AGCAATGCAAATTGATGCCAAAGAAGCTGCTCAACGTTTAGCGCAACATCTTCATGCGAAAGAATCTGCA<br>CTCTACCTTGATAATGAACAGCTTAAAGCAATTCAAGACAGCTATCAGCTTGTTGCAATGCCTAACCCAAG<br>TAATGTCATGACTCGTCATGCATTTATGATCAATGGTATGAATGCTGAATGTGCAGAGCTTGCACAAAATA<br>TTCGAACTGAAATTTTGATGGGTAAAAAATCAATTCCTCAAAATGACAGCCCGAGTAGTGCAGCATCTAGA<br>AAATT | BamHI,<br>XbaI |
| AdvA E350A | CAACTAGGATCCGAATCATAATCGAATTTCAATATAATTGCCTTGGAAGTTTAACTTGATTGCGCCTAGAC<br>AAGGGCTATTTGCTAGCCTACTGATCATTAGTTTTGCATTGCATACCTTTTTATTGGTGATTGCAACGACA<br>CACCAGCTCAATGAAAACCGTGCGAGTCAAGGTCAGCTCATGACCAGTCAACTCGTTGCAGACAGCTTAT<br>CTGAGCTTGAACCTGCCAATACGGTCTCTTTAGCTCTGATTGCTAATCGCTATGCGACCAATCCAAGTGT<br>GGCATCTATCCGTATTCTTGACGCCAATAAACAAGTTTTGGCAACAAGCGGCATGTCAAAAACTCGCCAA<br>GGTGAAATTTTTGTTCGTGATGCGCTTCAAAACGAGAAAAAAGTCGGTTCTATTGAAATCACACTAATTCA<br>ACCAAGTATTGGGGAAATTCTTCGTACCCAATGGCTTGCGATTTTAGCCTCCCTGTTCTTACATGTTTTATT<br>ATGGCTAGCTTACCGTGCCATTGCTCGCCCAACACGTAGCGAATACTTAGCCCGTATTAACGAAGAAAAT<br>CGTTTAAAACATGAAATTCAAGAGTTAACTCAAGCTCTCGCGCTTGAAAAACAAAATACAGTGACCTTGGT<br>TGCTCAAGCTCAGCAGCAAGCTAAAGCTAAACCAATTGTTCGCTCACAACCAGAAAAAAGCTTAGAGTCT<br>ACAGACCAAAATACCTTGGCACTTAATATTCAATTTTATGACCCTAAACAGTTATTAAGTAGTGTTAACCAG<br>TCTGTTTCTGTACCGTACTTTAAACTTTGTCAGCTCTTCTTAAACAAGAGTATTGAGCTATGTACAAAGCAT<br>TATCATTTGAAAGCGACTGATATTGATGTCGTAGACGAGTTTCATGCAGAGGGCGCAACACTTGCCATTT<br>CGACTTCACATCCACACGCTGTAGAGTGTCTACTTATGGTAGGTACGGTTTTCCAATTGCTTTCAGACGTT<br>TTATATAAGCGTTACCGTGAAGATAAACGCTTTGCGCTACAAACTCGCAGCGCCGTGTGTAATGCCGTAG<br>AAGCAATGCAAATTGATGCCAAAGCTGCTGCTCAACGTTTAGCGCAACATCTTCATGCGAAAGAATCTGC<br>ACTCTACCTTGATAATGAACAGCTTAAAGCAATTCAAGACAGCTATCAGCTTGTTGCAATGCCTAACCCAA<br>GTAATGTCATGACTCGTCATGCATTTATGATCAATGGTATGAATGCTGAATGTGCAGAGCTTGCACAAAAT<br>ATTCGAACTGAAATTTTGATGGGTAAAAAATCAATTCCTCAAAATGACAGCCCGAGTAGTGCAGCATCTAG<br>AAAATT | BamHI,<br>XbaI |
| AdvA R334A R354A | CAACTAGGATCCGAATCATAATCGAATTTCAATATAATTGCCTTGGAAGTTTAACTTGATTGCGCCTAGAC<br>AAGGGCTATTTGCTAGCCTACTGATCATTAGTTTTGCATTGCATACCTTTTTATTGGTGATTGCAACGACA<br>CACCAGCTCAATGAAAACCGTGCGAGTCAAGGTCAGCTCATGACCAGTCAACTCGTTGCAGACAGCTTAT<br>CTGAGCTTGAACCTGCCAATACGGTCTCTTTAGCTCTGATTGCTAATCGCTATGCGACCAATCCAAGTGT<br>GGCATCTATCCGTATTCTTGACGCCAATAAACAAGTTTTGGCAACAAGCGGCATGTCAAAAACTCGCCAA<br>GGTGAAATTTTTGTTCGTGATGCGCTTCAAAACGAGAAAAAAGTCGGTTCTATTGAAATCACACTAATTCA<br>ACCAAGTATTGGGGAAATTCTTCGTACCCAATGGCTTGCGATTTTAGCCTCCCTGTTCTTACATGTTTTATT<br>ATGGCTAGCTTACCGTGCCATTGCTCGCCCAACACGTAGCGAATACTTAGCCCGTATTAACGAAGAAAAT<br>CGTTTAAAACATGAAATTCAAGAGTTAACTCAAGCTCTCGCGCTTGAAAAACAAAATACAGTGACCTTGGT<br>TGCTCAAGCTCAGCAGCAAGCTAAAGCTAAACCAATTGTTCGCTCACAACCAGAAAAAAGCTTAGAGTCT<br>ACAGACCAAAATACCTTGGCACTTAATATTCAATTTTATGACCCTAAACAGTTATTAAGTAGTGTTAACCAG<br>TCTGTTTCTGTACCGTACTTTAAACTTTGTCAGCTCTTCTTAAACAAGAGTATTGAGCTATGTACAAAGCAT<br>TATCATTTGAAAGCGACTGATATTGATGTCGTAGACGAGTTTCATGCAGAGGGCGCAACACTTGCCATTT<br>CGACTTCACATCCACACGCTGTAGAGTGTCTACTTATGGTAGGTACGGTTTTCCAATTGCTTTCAGACGTT<br>TTATATAAGCGTTACGCTGAAGATAAAGCTTTTGCGCTACAAACTCGCAGCGCCGTGTGTAATGCCGTAG<br>AAGCAATGCAAATTGATGCCAAAGAAGCTGCTCAACGTTTAGCGCAACATCTTCATGCGAAAGAATCTGC<br>ACTCTACCTTGATAATGAACAGCTTAAAGCAATTCAAGACAGCTATCAGCTTGTTGCAATGCCTAACCCAA<br>GTAATGTCATGACTCGTCATGCATTTATGATCAATGGTATGAATGCTGAATGTGCAGAGCTTGCACAAAAT<br>ATTCGAACTGAAATTTTGATGGGTAAAAAATCAATTCCTCAAAATGACAGCCCGAGTAGTGCAGCATCTAG<br>AAAATT | BamHI,<br>XbaI |

**Table S4. AlphaFold3 confidence values for predicted AdvA assemblies.**

| Assembly | Interface | Chain-pair ipTM (mean, range) | PAE_min (Å) |
|---|---|---|---|
| $AdvA^{226-420}$ + $ZipA^{171-339}$ | AdvA-ZipA | 0.89 (0.89–0.89) | 1.0 |
| AdvA full-length dimer | AdvA-AdvA | 0.36 (0.36–0.36) | 3.3 |
| $AdvA^{226-420}$ (CHD-like domain) dimer | AdvA-AdvA | 0.38 (0.27–0.48) | 13.9 |
| $AdvA^{1-166}$ dimer | AdvA-AdvA | 0.56 (0.56–0.57) | 3.8 |
| $AdvA^{1-166}$ dimer + FtsN (full-length) | AdvA-FtsN | 0.20 (0.16–0.24) | 10.0 |
| | AdvA-AdvA | 0.60 (0.54–0.63) | 2.6 |
| $AdvA^{1-166}$ dimer + $FtsN^{1-133}$ (without SPOR) | AdvA-FtsN | 0.25 (0.22–0.28) | 8.0 |
| | AdvA-AdvA | 0.68 (0.68–0.69) | 1.8 |
| $AdvA^{1-166}$ dimer + FtsL + FtsB | AdvA-FtsL | 0.13 (0.12–0.16) | 15.8 |
| | AdvA-FtsB | 0.12 (0.11–0.14) | 16.1 |
| | AdvA-AdvA | 0.48 (0.44–0.51) | 4.3 |
| | FtsL-FtsB | 0.45 (0.43–0.48) | 3.8 |

Each assembly was modeled with AlphaFold3 using five independent seeds. Confidence scores are reported for the top-ranked model from each seed. Shown are interface predicted template modeling (ipTM) scores and minimum predicted aligned error (PAE_min) for the indicated chain pair across the seeds, extracted from the AlphaFold3 confidence outputs. For assemblies containing the AdvA dimer, the two AdvA protomer-partner pair ipTMs were within 0.04 of each other in all cases, and the more favorable score of the two was reported. ipTM values >0.8 indicate a confident interface and values <0.6 indicate low confidence (AlphaFold Server documentation).
